# Functional coexistence theory: a mechanistic framework linking biodiversity to ecosystem function

**DOI:** 10.1101/2024.05.05.591902

**Authors:** Joe Wan, Po-Ju Ke, Iris Hordijk, Lalasia Bialic-Murphy, Thomas W. Crowther

**Author notes:** These authors contributed equally (Joe Wan, Po-Ju Ke).

## Abstract

Theory and experiments show that diverse ecosystems often have higher levels of function (for instance, biomass production), yet it remains challenging to identify the biological mechanisms responsible. We synthesize developments in coexistence theory into a general theoretical framework linking community coexistence to ecosystem function. Our framework, which we term functional coexistence theory, identifies three components determining the total function of a community of coexisting species. The first component directly corresponds to the niche differences that enable pairwise species coexistence, and to the complementarity component from the additive partition of biodiversity effects. The second component measures whether higher functioning species also have higher competitive fitness, providing a missing link between the additive partition’s selection effect and modern coexistence theory’s concept of equalization. The third component is least well-studied: reducing functional imbalances between species increases niche difference’s positive effect on function. Using a mechanistic model of resource competition, we show that our framework can identify how traits drive the effect of competition on productivity, and confirm our theoretical expectations by fitting this model to data from a classic plant competition experiment. Furthermore, we apply our framework to simulations of communities with multiple ecosystem functions or more than two species, demonstrating that relationships between niche, fitness, and function also predict total function beyond the case studied by classical theory. Taken together, our results highlight fundamental links between species coexistence and its consequences for ecosystem function, providing an avenue towards a predictive theory of community–ecosystem feedbacks.

## 1 Introduction

All living systems obey the same set of physical laws, yet any individual ecosystem encompasses a unique assembly of organisms and interactions. This fundamental contrast is embodied by a traditional division within ecology: ecosystem ecology focuses on the flow of energy and nutrients as common currencies, while community ecology aims to explain the diversity of organisms. However, understanding ecosystems requires ecologists to acknowledge the fundamental links between these aspects: ecosystem flows affect community composition; in turn, ecological communities control ecosystem cycles of energy and nutrients. Thus, general theories of ecosystems must account for the feedback between ecosystem and community processes. As human activity simultaneously perturbs global element cycles and threatens local biodiversity, understanding such feedback is a fundamental ecological challenge with enormous practical consequences for understanding and mitigating global change.

One successful body of research, termed *biodiversity–ecosystem function*, studies this feedback by asking how diversity at the community level affects function at the ecosystem level (e.g. biomass production, nutrient cycling, or ecosystem services). This field has combined manipulative experiments (Hector, Bazeley-White, et al. 2002; Hooper, Chapin III, et al. 2005) and theoretical analyses (Connolly et al. 2013; Loreau and Hector 2001; Turnbull et al. 2013) to highlight the effect of biodiversity on ecosystem processes such as primary productivity, nutrient cycling, and ecosystem services (Hector, Bazeley-White, et al. 2002; Isbell et al. 2017). The effect of biodiversity on ecosystem function can be partitioned into two components: *complementarity*, which measures whether species function better on average within communities versus growing alone (e.g., due to underlying niche differentiation) and *selection*, which measures whether higher-functioning species disproportionately dominate a community (Loreau and Hector 2001). This approach, termed the *additive partition* of biodiversity effects, and subsequent related frameworks (Bannar-Martin et al. 2018; Connolly et al. 2013; Fox 2005; Liang, Zhou, et al. 2015) have been applied to a variety of experiments and empirical studies. Taking advantage of this theoretical–empirical synthesis, a cross-scale perspective has emerged (Cardinale, Hillebrand, et al. 2009; Hooper, Adair, et al. 2012; O’Connor et al. 2017) emphasizing the positive effects that biodiversity often has on ecosystem function.

Nonetheless, the degree to which biodiversity promotes community-level functioning varies greatly between systems (O’Connor et al. 2017). While most work has focused on biomass production in terrestrial plants, the positive diversity–function relationships observed there may not generalize across ecosystem types (O’Connor et al. 2017) with different species pools, environmental conditions (Spaak, Baert, et al. 2017), or community structures (Hordijk et al. 2023). Indeed, in certain highly competitive systems, consistently negative diversity–function relationships may be the norm (Maynard et al. 2017). Furthermore, even within systems, biodiversity effects vary during community succession (Weis et al. 2007), suggesting that observed biodiversity effects may only be transient (Turnbull et al. 2013). Accordingly, though recent empirical (Gonzalez et al. 2020; Liang, Crowther, et al. 2016) and modeling work (Pavlick et al. 2013) has begun to focus on applying the insights of diversity–function studies at large scales, synthesizing a general predictive theory of ecosystem function remains challenging. Thus, an important current challenge for understanding and predicting community–ecosystem feedbacks is identifying the underlying ecological mechanisms—that is, interactions between species and their environment— through which diversity affects function (Hector, Bell, et al. 2009; Loreau 2010; Loreau, Sapijanskas, et al. 2012; Mouquet et al. 2002).

Just as the additive partition has provided a unifying tool for linking diversity to ecosystem function, a body of theory known as *modern coexistence theory* has provided a general framework for understanding and predicting the maintenance of diversity itself. As a quantitative currency for coexistence, the theory identifies two processes: *stabilization*, which prevents competitive exclusion by reducing species’ relative negative effects on each other (and thus is also termed *niche difference*), and *equalization*, which reduces competitive imbalances between species (termed *fitness differences*) such that stabilization can ensure coexistence (Chesson 2000; Ke and Letten 2018). In contrast to the additive partition approach, which was developed to test empirical hypotheses in biodiversity–ecosystem function experiments (Loreau and Hector 2001, 2019; Wagg et al. 2019), modern coexistence theory was first proposed to provide mechanistic predictions of coexistence in theoretical models (Chesson 2000). Indeed, its metrics have successfully been applied to predict how a variety of specific biological mechanisms contribute to coexistence in theoretical (Ke and Wan 2020; Letten, Ke, et al. 2017; Spaak, Ke, et al. 2023) and empirical studies (Godoy and Levine 2014; Johnson et al. 2022; Petry et al. 2018). Accordingly, studies have related niche and fitness measures from modern existence theory to ecosystem function (Carroll et al. 2011; Turnbull et al. 2013), though subsequent debate has questioned the generality and applicability of this approach (Loreau and Hector 2019; Loreau, Sapijanskas, et al. 2012; Pillai and Gouhier 2019; Wagg et al. 2019). Thus, despite calls to adopt a more mechanistic view of biodiversity–function relationships (Ratcliffe et al. 2017; Wang et al. 2024) and recent work comparing these relationships to niche and fitness metrics (Godoy, Gómez-Aparicio, et al. 2020), there is no general framework extending the predictive power of coexistence theory to address communities’ total function.

Building upon this emerging synthesis, we apply modern coexistence theory to provide a general mechanistic framework for biodiversity effects. Our approach, which we term *functional coexistence theory*, highlights the importance of considering species’ functional imbalances in tandem with their classical niche and fitness differences. Integrating these components, researchers can quantify the mechanisms governing coexistence between species in order to predict how the resulting community is likely to function. First, we use classic competition models to illustrate our framework (section “Extending modern coexistence theory to predict function”) by deriving conditions for one kind of biodiversity effect (transgressive overyielding). Accordingly, we identify three processes determining the total function of a community: stabilizing niche difference, fitness–function relationships, and functional equalization. Next, we place our functional coexistence framework within the context of the rich literature on biodiversity–ecosystem function to show that the two approaches are compatible despite their quantitative differences (section “Placing functional coexistence theory in the context of the biodiversity–function literature”). Moreover, we show how our framework can identify mechanistic drivers of ecosystem function (section “Linking functional coexistence theory to biological mechanism”). Using a general trait-based model of resource competition, we show how functional coexistence theory can predict the effect of traits on total function, and confirm these predictions by reinterpreting a classic plant competition experiment with our framework (Wedin and Tilman 1993). Finally, we demonstrate how our theory holds when expanded to study multifunctionality and multispecies communities (section “Beyond classic theory: applying functional coexistence theory to multiple functions and species”). Taken as a whole, our proposed framework clarifies the fundamental links between coexistence and ecosystem function. Thus, by synthesizing a mechanistic understanding of diversity–function relationships, our results can help predict how ecosystems, along with the key services they provide, will respond to change.

## 2 Extending modern coexistence theory to predict function

In this section, we illustrate how modern coexistence theory’s niche and fitness measures can be integrated with measures of species’ function in order to predict ecosystem function, beginning with a quantitative two-species framework frequently employed in empirical studies of coexistence. Just as modern coexistence theory classifies processes affecting coexistence into stabilizing and equalizing components (section “Modern coexistence theory: two components maintaining diversity”), our functional coexistence framework identifies three components contributing to the total function of communities. We illustrate these components by considering biomass production in the two-species model (Box 1 and section “Functional coexistence theory: three components driving overyielding”), focusing specifically on *transgressive overyielding*, which occurs when a community’s total function exceeds that of its most productive species (Loreau 2010).

### 2.1 Modern coexistence theory: two components maintaining diversity

Modern coexistence theory highlights that differences between species can affect coexistence in two ways: they may promote coexistence by helping all species in a community invade (i.e., recover from low abundance), or hinder coexistence by favoring certain species over others. Accordingly, species coexistence can be predicted from two metrics summarizing these roles: niche differences (ND) promote coexistence, while fitness differences (FD) hinder coexistence. Stated conceptually, coexistence occurs when niche differences are greater than fitness differences (ND *>* |FD|), allowing all species to attain positive invasion growth rates (Barabás et al. 2018). Following the framework of Ke and Letten (2018) and Letten, Ke, et al. (2017), we depict these requirements for a two-species system in Figure 1a.

**Figure 1:**
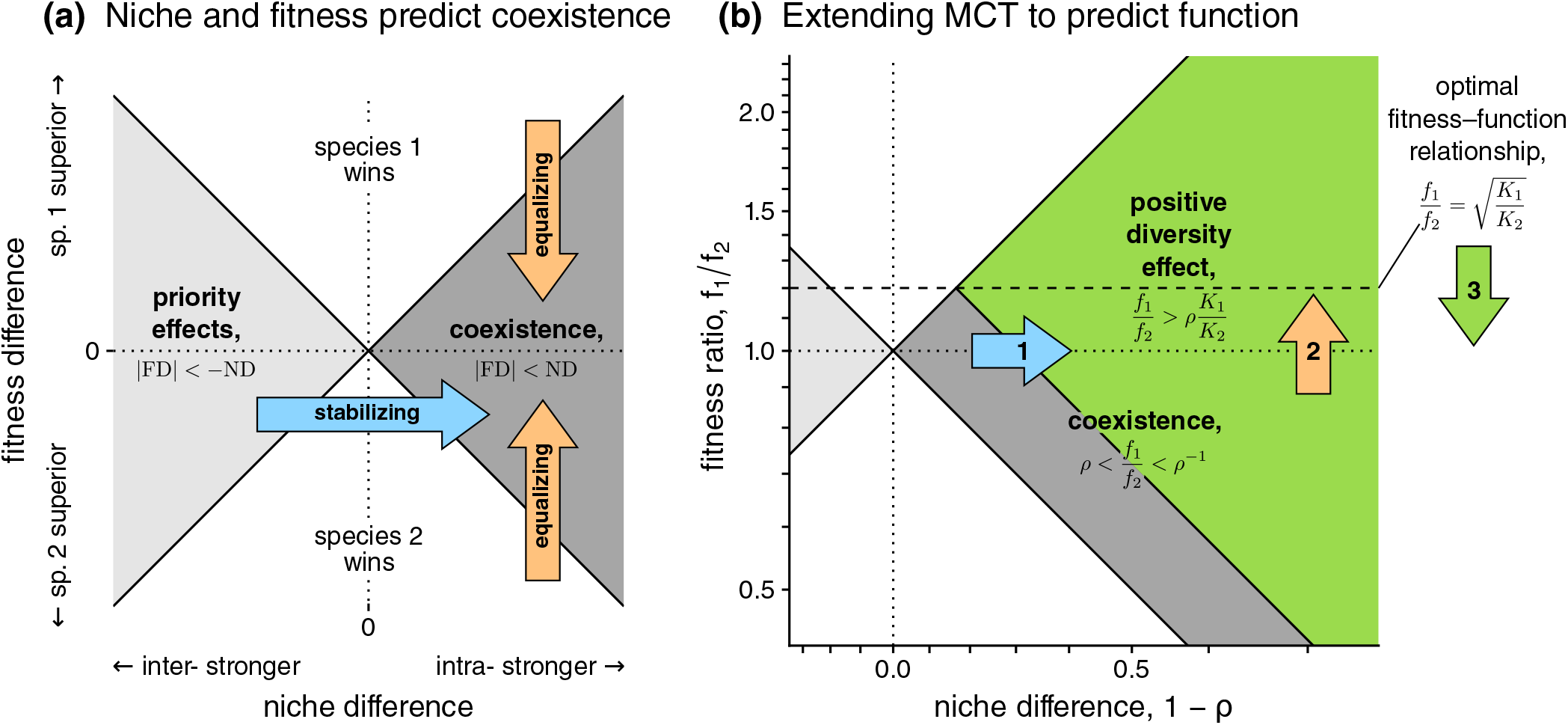
Modern coexistence theory and its functional extension. **(a) Modern coexistence theory: niche and fitness differences predict coexistence**. Coexistence outcomes between two species depend on niche difference, ND (horizontal axis), and fitness difference, FD (vertical axis; panel adapted from Mordecai 2011). Coexistence (dark gray) requires niche difference to be positive (ND *>* 0) and large enough to overcome fitness difference (*ND >* |FD|). Any process promoting coexistence can be partitioned into two components: stabilizing (blue arrow), i.e., increasing niche difference, and equalizing (orange arrows), i.e., decreasing the magnitude of fitness difference towards zero. Here, ND and FD are notated conceptually, but can be quantified for specific models, as discussed below. **(b) Functional coexistence theory: extending the modern coexistence framework to predict function**. We now use niche and fitness to predict whether species interactions cause a community to outperform the best single species, termed *transgressive overyielding*. To match panel (a), axes are logarithmically transformed (− log *ρ* and log *f*_1_/ *f*_2_), but we label corresponding values of the more familiar measures from the literature (1− *ρ* and *f*_1_/ *f*_2_). On top of the conditions for coexistence, a positive diversity effect (green region) only occurs when the higher yielding species (here, species 1) also has sufficiently high fitness (*f*_1_/ *f*_2_ *> ρ·K*_1_/*K*_2_) or equivalently, when niche difference and 1’s fitness advantage are in excess of those required for coexistence (ND −Δ *>* |FD− Δ|, where 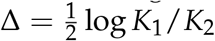. Accordingly, processes affecting function can be partitioned into effects on niche (arrow 1) and fitness (arrow 2), as previously, but also on functional imbalance between the species (arrow 3).

Thus, processes maintaining diversity can be classified according to these two components. The first, *stabilization*, increases niche differences (Figure 1a, blue arrow); to clarify their role in coexistence, niche differences are therefore sometimes termed *stabilizing niche differences*. The second, *equalization* (Figure 1a, orange arrows), makes species more similar in fitness, thereby reducing competitive hierarchy and preventing exclusion (Figure 1a, orange arrows). This stabilizing–equalizing framework does not directly quantify biological mechanism because its components do not directly correspond to concrete biological processes: Barabás et al. 2018; Loreau, Sapijanskas, et al. 2012. However, applied to mechanistic models, it provides a powerful tool for summarizing how coexistence can arise through processes ranging from abiotic interactions such resource use (Letten, Ke, et al. 2017; Song et al. 2019) to biotic interactions such as pollination (Johnson et al. 2022), mutualism (Kandlikar et al. 2019; Ke and Wan 2020), or disease (Mordecai 2011).

#### Box 1.

Linking modern coexistence theory to ecosystem function

For a class of commonly-used competition models, we can use the niche and fitness components of modern coexistence theory to calculate total ecosystem function. As a representative example, we consider conditions for transgressive overyielding in the classic Lotka–Volterra model, where the dynamics of species *i*’s population *N*_*i*_ follow:

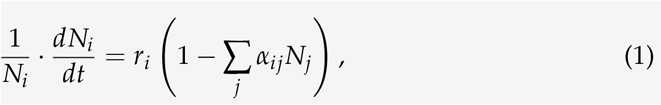

where *r*_*i*_ is species *i*’s intrinsic rate of increase and *α*_*ij*_ is the per-capita competitive effect of species *j* on species *i*. A more general analysis and full derivations are given in Appendix S1, and conditions for other outcomes in Appendix S2; we also show in Appendix S3 that the results can be extended to models with nonlinear competitive effects. Note that we discuss biomass here for simplicity, but that results are fully analogous for any function Φ instead of biomass *K*, as discussed in Appendix S8.

##### Modern coexistence theory: niche and fitness measures

In a two-species community, the niche and fitness components are (per Chesson and Kuang 2008):

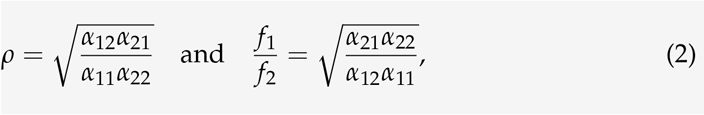

which respectively give the niche overlap and fitness ratio between species 1 and 2. Two conditions allow the species to stably coexist, each corresponding to one of the coexistence components in modern coexistence theory. First, species must experience niche differentiation: *ρ* < 1, ensuring that within-species competition is stronger than between-species competition. Second, species must be sufficiently similar in competitive ability:

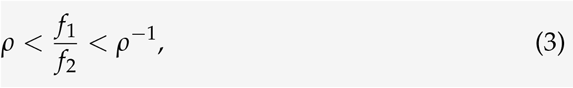

ensuring that the fitness ratio between species is not too imbalanced relative to niche differentiation. We illustrate these conditions in Figure 1a; since *ρ* and *f*_1_/ *f*_2_ are ratios, we take logarithms to obtain niche and fitness differences corresponding to the conceptual discussion in section 2.1 (ND = − log *ρ* and FD = log *f*_1_/ *f*_2_; Johnson et al. 2022; Yamamichi et al. 2022), where coexistence requires ND *>* |FD|, though we label the axes with the more familiar units of 1 − *ρ* and *f*_1_/ *f*_2_.

##### Fitness determines species’ contributions to total function

To link these measures to function, we first focus on 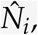the biomass that species *i* contributes to the community. We show that each species’ biomass is proportional to its intrinsic yield 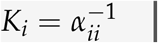 (i.e., carrying capacity) and to its scaled invasion growth rate *F*_*i*_ = 1 − *α*_*ij*_/*α*_*jj*_ (i.e., its invasion growth rate divided by *r*_*i*_). This gives a straightforward expression for biomass of species *i* at equilibrium,

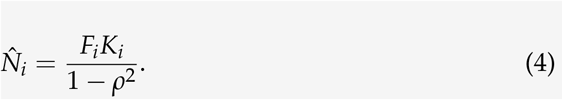

Here, as in more general versions of modern coexistence theory (Barabás et al. 2018), *F*_*i*_ measures a species’ fitness—its ability to persist under competition with the rest of its community. We apply *F*_*i*_ to simplify the derivation of our results and emphasize their link to invasion analysis (Grainger et al. 2019). Though the wide applicability of modern coexistence theory is underpinned by invasion analysis, which considers dynamics when one species is rare (Grainger et al. 2019), the theory gains considerable predictive power because invasion growth rates also predict a system’s long-term trajectory and properties (Arnoldi et al. 2022; Barabás et al. 2018). Accordingly, the traditionally-defined Lotka–Volterra fitness ratio *f*_1_/ *f*_2_ quantifies two species’ imbalance in *F* as 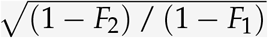 while niche overlap *ρ* quantifies competitive reduction in both species’ *F* as 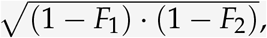, and we can apply the identity

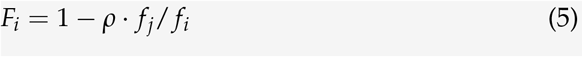

to relate the two sets of measures.

##### Degree of transgressive overyielding

Using equation 4 to write total biomass as 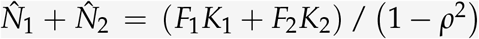, we can investigate the relative degree of transgressive overyielding, i.e., the difference between the biomass of the total community and that of its highest-yielding single species. Without loss of generality, we designate species 1 as the highest-yielding single species (*K*_1_ *> K*_2_). In order to understand the conditions promoting transgressive overyielding, we can rewrite the total biomass as

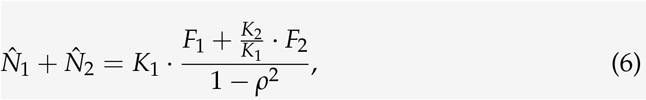

There are three ways to change the value of this expression relative to *K*_1_, the baseline for transgressive overyielding: (1) changing *stabilization*, i.e., how close *ρ* is to 0; (2) changing *fitness imbalance*, i.e., the relative magnitudes of *F*_1_ and *F*_2_ for a particular value of *ρ*; and (3) changing *yield imbalance*, i.e., how close the yield ratio *K*_2_/*K*_1_ is to 1. Note that these components are not fully independent due to the relationship between *ρ, F*_1_, and *F*_2_.

##### Conditions for overyielding

Our analysis allows us to derive simple conditions for transgressive overyielding. Solving the conditions under which total biomass (equation 6) is greater than the best intrinsic yield *K*_1_ (and rewriting *ρ* in terms of *F*_1_, *F*_2_) gives *F*_1_ *>* 1 − *K*_2_/*K*_1_. In other words, transgressive overyielding requires the higher yielding species to also have sufficiently high fitness—a *fitness–function relationship*. Rewriting this in terms of *ρ* and *f*_1_/ *f*_2_ (Appendix S1) gives the *functional coexistence theory* condition for transgressive overyielding:

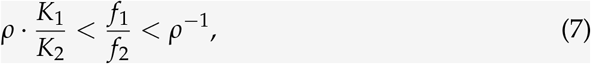

where *K*_1_/*K*_2_ *>* 1, and the upper bound of *ρ*^−1^ is due to the fact that coexistence is a prerequisite for transgressive overyielding. As niche overlap *ρ* decreases (i.e. species experience increasing niche differentiation), transgressive overyielding first becomes possible when *ρ*·*K*_1_/*K*_2_ = *f*_1_/ *f*_2_ = *ρ*^−1^ and thus at

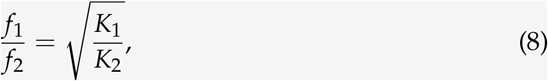

which, as we show in Appendix S1, is also more generally the fitness ratio maximizing total biomass. Illustrated in Figure 1b, these conditions are closely related to the coexistence condition from modern coexistence theory (equation 3). Simply put, transgressive overyielding requires that the niche difference and fitness advantage felt by the higher yielding species are in excess of those required for coexistence: in the conceptual notation of Section 2.1 and Figure 1a, ND − Δ *>* |FD − Δ|, where ND, FD are defined logarithmically as above, and 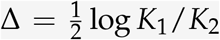 measures the yield imbalance to be overcome (Appendix S1).

#### 2.2 Functional coexistence theory: three components driving overyielding

In Box 1, we extend modern coexistence theory in order to include ecosystem function by relating its niche and fitness measures to the total function of the community. As a representative example, we consider the conditions under which the community’s biomass production shows *transgressive overyielding*; that is, when total biomass at equilibrium exceeds the biomass of each species growing alone (Loreau 2010). While we use the familiar Lotka–Volterra model as an illustration, our results (Appendix S1) rest upon a more general finding that in many models of competition, a species’ relative contribution to total biomass can be determined from two quantities: (1) its intrinsic yield *K*_*i*_, or biomass produced when growing alone, and (2) its fitness *F*_*i*_, or ability to persist under competition (Box 1: equation 4). This includes, in addition to the Lotka–Volterra model used in Box 1, many models with nonlinear competitive responses such as the Beverton–Holt model (Beverton and Holt 1957) used to study annual plant competition (Levine and HilleRisLambers 2009) and Tilman (1982)’s substitutable resource competition model (Letten, Ke, et al. 2017). Furthermore, this relationship is approximately true in an even broader class of models (Arnoldi et al. 2022), enabling further generalizations (for instance, nonlinear competitive responses, which we illustrate in Appendix S3).

Using this result, we combine the niche and fitness measures from modern coexistence theory (here, *ρ* and *f*_1_/ *f*_2_) with each species’ intrinsic yield (*K*_1_, *K*_2_) to fully predict total biomass and how it responds to community coexistence (Box 1: equation 7).

We identify three processes that enable transgressive overyielding, where the community outperforms its best single species (Loreau 2010), as depicted in Figure 1b. The first is simply stabilizing niche differences: increasing niche difference (decreasing *ρ* towards zero) tends to increase total biomass (arrow 1). The second concerns the relationship between fitness and function: transgressive overyielding occurs when the higher yielding species has a competitive advantage in excess of that needed for it to persist (arrow 2). The third component can be termed *functional equalization*: making the species more similar in intrinsic yield (decreasing *K*_1_/*K*_2_ towards 1) increases the potential for transgressive overyielding (arrow 3). In this section, we simulate how these components affect total biomass in the Lotka–Volterra model (Figure 2) and use these results to illustrate the interpretation of each process.

**Figure 2:**
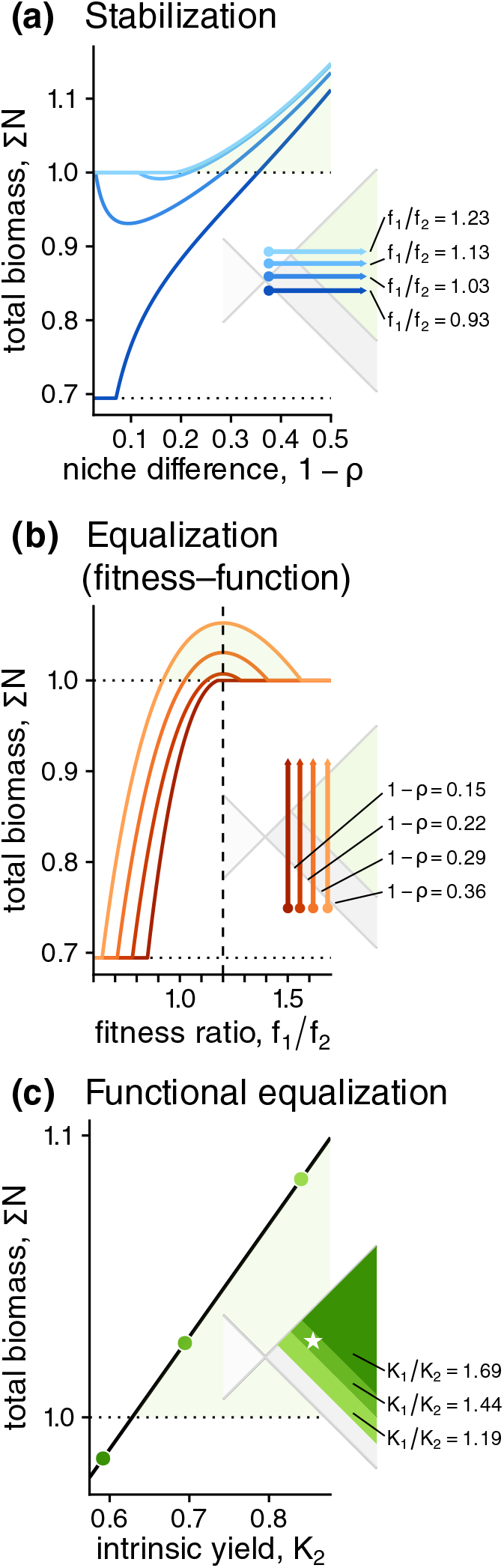
Illustrating causes for positive diversity effects using the Lotka–Volterra model. In each panel, we show the effect (solid lines/points) of varying a component (horizontal axis) on total biomass (vertical axis) as compared to the biomass of each species growing alone (dotted horizontal lines; set to 0.7 for the less-productive species 2 and 1.0 for the more-productive species 1). Insets show parameter values and positions on the coexistence space plot in Figure 1b. **(a) Stabilizing niche differences**. Increasing niche difference 1 − *ρ* eventually results in total biomass exceeding the yield of the best species (transgressive overyielding), regardless of fitness difference (line color), though biomass may decrease when niche difference is low. Note that the line for *f*_1_/ *f*_2_ = 1.23 overlaps or is slightly above that for *f*_1_/ *f*_2_ = 1.13. **(b) Fitness–function relationship**. Transgressive overyielding occurs and total biomass is maximized at the optimal fitness ratio 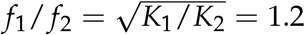 (vertical dotted line), provided niche difference (line color) is high enough to allow coexistence at this fitness ratio. **(c) Functional equalization**. Making species more equal in function by increasing productivity of the inferior species *K*_2_ while niche and fitness remain fixed (star in inset) increases total biomass by increasing the potential for transgressive overyielding (different green regions in inset). See Appendix S9 for parameter values.

##### Stabilizing niche differences

In the pairwise models we consider, niche difference (and equivalently, stabilization) can be interpreted as the tendency for intraspecific interactions to be more negative than interspecific ones: that is, for species to limit themselves more strongly than they limit each other. Confirming previous results (Carroll et al. 2011), we find that such niche differences tend to promote total biomass yield (Figure 2a). However, we caution that functional and competitive imbalances can complicate this relationship: when the higher-yielding species had only moderately higher fitness than its competitor (*f*_1_/ *f*_2_ = 1.03 and 1.13), increasing niche difference just enough to allow coexistence decreased total biomass. Thus, transgressive overyielding generally requires niche differentiation in excess of that simply required for coexistence (e.g., Figure 1b, where the transgressive boundary for overyielding lies to the right of the coexistence boundary). Nonetheless, regardless of the fitness ratio between coexisting species, sufficiently high niche difference always eventually enabled transgressive overyielding. Accordingly, we follow previous work in emphasizing that niche differentiation plays an essential role in allowing diversity to promote ecosystem function.

##### Fitness–function relationship

Modern coexistence theory highlights the role of competitive fitness (*F*_*i*_ or *f*_*i*_/ *f*_*j*_), the ability for a species to persist in a community (as measured by invasion analysis); indeed, without the context provided by fitness differences, predicting coexistence is impossible (Adler, HilleRisLambers, et al. 2007; Kandlikar et al. 2019). Going further, our functional framework highlights that fitness also determines the degree to which each species contributes to total ecosystem function. We find that transgressive overyielding requires precise relationships between fitness and function: namely, a species with a higher function (here, intrinsic yield for biomass *K*) must also have a sufficiently high competitive ability (as measured by *F*_*i*_ or *f*_*i*_/ *f*_*j*_; Figure 2b). In other words, higher-functioning species must have fitness in excess of that required for coexistence (by a factor of *K*_1_/*K*_2_; equation 7). Our simulations highlight that this component can be viewed as a version of modern coexistence theory’s equalization: regardless of niche difference, bringing fitness ratio towards its optimum value (vertical dashed line, 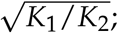; equation 8) always promoted transgressive overyielding, just as bringing it towards 1 would have promoted coexistence. Thus, our functional framework generalizes modern coexistence theory by showing that fitness differences also determine ecosystem function.

##### Functional equalization

Finally, we identify a driver of diversity effects with no direct equivalent from modern coexistence theory: functional equalization, which increases ecosystem function by reducing functional imbalances between species (e.g., low vs. high biomass production). As our simulations illustrate (Figure 2c), making coexisting species more equal in function always promotes transgressive overyielding because it reduces the opportunity for competition to select (i.e., increase the relative abundance of) functionally inferior species (i.e., the gray region in the inset becomes smaller). Functional equalization amplifies the effect of stabilizing niche differences: when species have equal function, transgressive overyielding always occurs as a consequence of stable coexistence. In this extreme, the previous fitness–function relationships become irrelevant because species do not differ in function, a scenario implicitly considered by classic experimental analyses designed for communities where species have similar intrinsic yields (e.g., the relative yield total approach: de Wit 1960). While functional imbalance has been discussed as a caveat for the interpretation of such studies (Schmid et al. 2008; Wagg et al. 2019), it has received little attention as an explanation of biodiversity effects in its own right; thus, we highlight its importance in predicting the total function of a community.

### 3 Placing functional coexistence theory in the context of the biodiversity–function literature

Although previous work has used concepts from coexistence theory to examine questions from the biodiversity–function literature, it has remained unclear whether these two approaches can be reconciled. In this section, we place functional coexistence theory within the context of the biodiversity–ecosystem function literature in order to show how it complements previous approaches. We begin by briefly summarizing the interpretation of the additive partition’s complementarity and selection components: though their respective links with niche and fitness have long been noted, the broader compatibility of the two frameworks has remained contentious (subsection “Previous attempts to synthesize diversity–function and theories of coexistence”). After quantitatively relating and niche and fitness measures to the complementarity and selection components (Box 2), we highlight how the perspective of functional coexistence theory resolves apparent contradictions between the theories (subsection “Comparing the niche–fitness and additive partition frameworks”). Although our presentation of functional coexistence theory has focused on transgressive overyielding, we then show how it can be applied to predict other outcomes, highlighting its flexibility as a predictive framework for total community function (subsection “Predicting different outcomes using niche, fitness, and function”).

#### 3.1 Previous attempts to synthesize diversity–function and theories of coexistence

The links between productivity and the processes allowing species to coexist have been noted since early efforts to quantify competition (de Wit 1960), culminating in quantitative descriptions of niche partitioning between species (e.g., MacArthur 1970). Building upon this perspective, studies from the biodiversity–ecosystem function literature (reviewed in Hooper, Chapin III, et al. 2005) have hypothesized that such niche partitioning effects may explain widely-observed positive effects from diversity manipulation experiments.

To synthesize the diversity of metrics and hypotheses from this field, Loreau and Hector (2001) proposed the *additive partition* of such biodiversity effects into two components. The first, *complementarity*, is an average indicating how much more species tend to yield in communities than growing alone, which can serve to quantify the role of niche partitioning and other interactions such as facilitation (Hooper, Chapin III, et al. 2005; Loreau 2004; Loreau, Sapijanskas, et al. 2012; Turnbull et al. 2013). The second component, *selection*, measures effects that depend on species identity by quantifying the tendency for species with higher intrinsic yield to contribute more to communities. As Loreau and Hector (2001) originally suggested (and later refined by Fox 2005), this selection component measures competitive differences in a manner analogous to fitness in evolutionary studies (Price 1995). Thanks to its generality, the additive partition has successfully summarized a large and diverse set of experimental studies (Cardinale, Matulich, et al. 2011). Nonetheless, as long noted (Hooper, Chapin III, et al. 2005; Loreau and Hector 2001; Mouquet et al. 2002), it does not identify specific biological processes driving biodiversity effects, nor does it predict how they might change with respect to time or environmental context.

More recent work has suggested that modern coexistence theory may help address limitations of the additive partition by helping to detect the biological mechanisms responsible for biodiversity effects (Carroll et al. 2011; Godoy, Gómez-Aparicio, et al. 2020; Turnbull et al. 2013). Indeed, the framework formalizes the same ecological concepts as the additive partition: like complementarity, niche difference measures processes reducing the importance of competition between species; like selection, fitness measures processes favoring one species over another (Adler, HilleRisLambers, et al. 2007). Accordingly, theoretical work has aimed to relate the approaches (Turnbull et al. 2013); towards this goal, Carroll et al. (2011) suggested the additive partition may misrepresent underlying mechanisms (e.g., resource partitioning), and proposed using niche difference as an alternative metric for diversity–function studies. However, a subsequent exchange questioned whether either approach appropriately indexes underlying mechanisms (Carroll et al. 2012; Loreau, Sapijanskas, et al. 2012), while more recent debate has stressed their different and potentially incompatible conceptual aims (Loreau and Hector 2019; Pillai and Gouhier 2019; Wagg et al. 2019). Thus, despite recent calls to harness ecological theory to identify mechanisms for biodiversity effects (Godoy, Gómez-Aparicio, et al. 2020; Ratcliffe et al. 2017; Wang et al. 2024), it remains unclear how to integrate the general insights offered by modern coexistence theory within the field of biodiversity–ecosystem function.

##### Box 2.

Relating functional coexistence to other frameworks for diversity effects

Loreau and Hector (2001) defined the *additive partition of biodiversity effects* by showing that Δ*Y*, the difference between observed total yield and expected yield *Y*_*E*_, can be written as

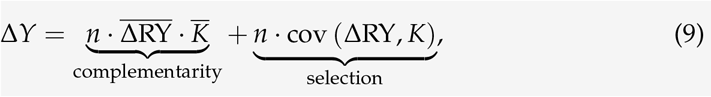

where *n* is the number of species, *K* is intrinsic yield or function when growing alone, RY_*i*_ is relative yield (a species’ yield within the community divided by its intrinsic yield), and 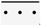, cov (*·*·*·*), and Δ·*·*·respectively denote these quantities’ mean, covariance, and deviation from experimenters’ expectations. Expected yield is the weighted average of intrinsic yields according to expected relative yields (*Y*_*E*_ = Σ_*i*_ RY_*E*,*i*_*K*_*i*_); a typical choice of RY_*E*,*i*_ is species’ proportions at the beginning of an experiment, but equation 9 is valid for any choice of expected relative yield. Here, following previous studies (Carroll et al. 2011; Loreau 2010), we consider changes relative to average intrinsic yield 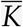 (corresponding to RY_*E*,*i*_ = 1/*n*); note that this differs from the derivation in Box 1, which focused on transgressive overyielding (i.e., relative to *K*_1_).

###### Relating the additive partition to niche and fitness

We relate the additive partition to niche and fitness measures for Box 1’s competition models (Appendix S4) by considering the coexistence equilibrium. Noting that RY_*i*_ is our 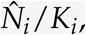, we find that complementarity is [Σ *F*/ 1− *ρ*^2^) − 1] *·*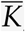, regardless of the choice of expected relative yield, and selection is *n*·cov (*F, K*) / (1 − *ρ*^2^), provided all expected relative yields are equal, i.e., RY_*E*,*i*_ = 1/*n*. In this case, expected yield is simply average intrinsic yield 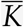 and we can write these expressions out in full for the equilibrium abundances of (coexisting) species 1 and 2 as:

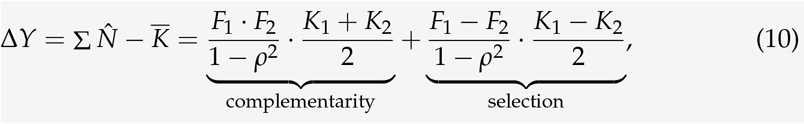

corresponding to previous results (Carroll et al. 2011; Loreau, Sapijanskas, et al. 2012) except that we have simplified the expression by keeping *F*_1_, *F*_2_. As previously noted by Carroll et al. (2011), these expressions have complicated relationships with *ρ* and *f*_1_/ *f*_2_; nonetheless, the form of equation 10 suggests that complementarity is related to the tendency of both *F*_1_ and *F*_2_ to be large, while selection is related to the difference between *F*_1_ and *F*_2_. We conform these expectations in Figure 3a–c; using equation 5, we can also show them to hold exactly (Appendix S4).

**Figure 3:**
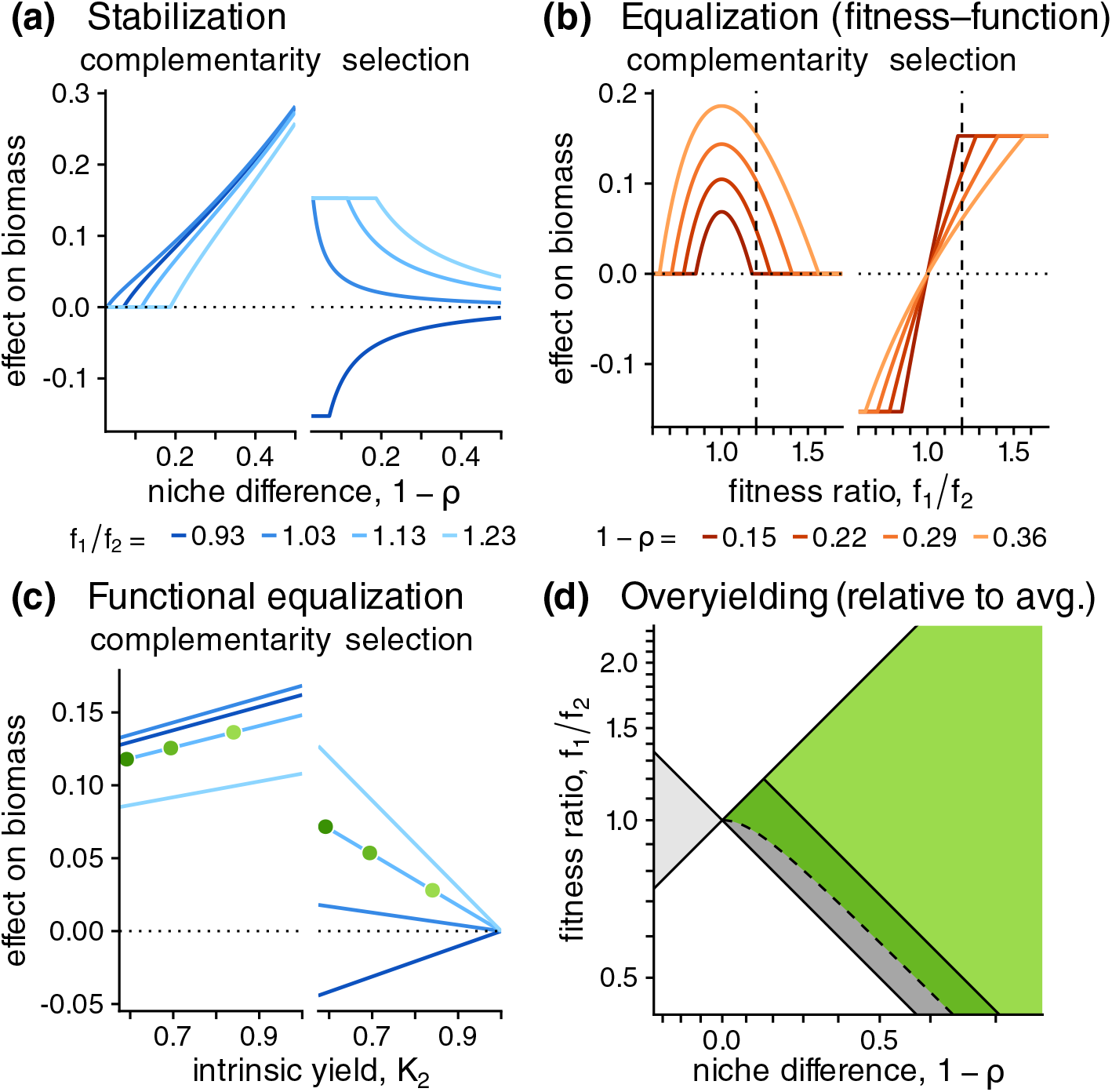
Linking functional coexistence theory components with previous approaches. **(a)– (c) Relating stabilization, fitness–function, and functional equalization to the additive partition**. Following the simulations in Figure 2 (matching parameter values and legends), we show how each of the components of functional coexistence theory are related to the additive partition by using species’ equilibrium biomass to calculate the complementarity and selection effects as in Box 2. Instead of only showing one set of niche and fitness values (as in Figure 2c), panel (c) uses the same four values of *f*_1_/ *f*_2_ (line color) as in panel (a). **(d) Conditions for overyielding relative to average intrinsic yield**. On top of the niche–fitness space in Figure 1b, we visualize the conditions for overyielding relative to 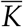, the average of species 1 and 2’s intrinsic yields (dark green region); the boundary (dashed line) is the full condition given in Box 2. This defines a curve that becomes nearly parallel to the transgressive overyielding and coexistence boundaries at high enough niche difference.

###### Overyielding relative to average intrinsic yield

As exemplified by the additive partition’s freely chosen *Y*_*E*_, there are many ways to quantify overyielding; Appendix S2 considers these in a fully general way. Here, we demonstrate the case where the community outperforms species’ average intrinsic yield 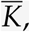, corresponding to the choice of RY_*E*,*i*_ = 1/*n* in the additive partition, used above and previously in the literature (Carroll et al. 2011; Loreau, Sapijanskas, et al. 2012). Solving for the condition under which equation 10 is positive gives the inequality

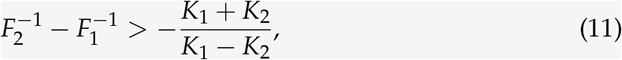

where the left hand side measures whether species 1 has a greater value of *F*_*i*_. This condition is illustrated as the dashed line in Figure 3d and can be rewritten in terms of *f*_1_/ *f*_2_ and *ρ* using equation 5 (see more complicated expression in Appendix S2).

#### 3.2 Comparing the niche–fitness and additive partition frameworks

In Box 2 (and Appendix S4), we calculate how complementarity and selection are related to the niche, fitness, and function components from functional coexistence theory, finding conceptual agreement between the frameworks despite quantitative differences. We show these in Figure 3 by calculating the complementarity and selection components for the same scenarios originally simulated in Figure 2; we also provide general proofs of these findings in Appendix S4. As we increased niche difference (Figure 3a), complementarity always increased with increasing niche difference, while selection did not change in a consistent direction: it either increased (*f*_1_/ *f*_2_ = −0.93) or decreased (other values of *f*_1_/ *f*_2_) depending on the underlying fitness difference. Meanwhile, increasing fitness ratio (Figure 3b) caused complementarity to increase until *f*_1_/ *f*_2_ = 1 and then decrease; this effect occurred at all niche difference values. On the other hand, regardless of niche difference, selection consistently increased with fitness ratio: it was negative when fitness favored the lower yielding species 2, increasing to 0 when *f*_1_/ *f*_2_ = 1, and becoming positive when the fitness ratio favored the higher yielding species 1. Finally, increasing the intrinsic yield of the lower yielding species *K*_2_ (Figure 3c) slightly increased complementarity (though this effect would disappear when standardizing by the average intrinsic yield: see general result in Appendix S4). Regardless of the fitness ratio, doing so also reduced the magnitude of selection, such that selection was always 0 when *K*_2_ = *K*_1_ = 1.

Thus, we conclude that the coexistence theory and additive partition components are closely linked: increased stabilization consistently corresponds to increased complementarity, while increasing the fitness ratio in favor of the higher yielding species consistently corresponds to increased selection. Meanwhile, reducing functional imbalance reduces selection effects, which vary in sign depending on species’ fitness, leaving complementarity, which is always positive for coexisting species. This confirms previous suggestions that stabilization and complementarity are closely linked (Carroll et al. 2011; Loreau 2004), and that selection is related to fitness (Fox 2005). Indeed, it is possible to show that selection can be interpreted as a niche difference metric for modern coexistence theory, closely related to previously proposed metrics based on arithmetic means of invasion growth rates (Barabás et al. 2018; Chesson 2003; Spaak, Ke, et al. 2023; Zhao et al. 2016); we show this result in Appendix S5. Furthermore, despite quantitative differences, the two approaches made similar inferences regarding the primary drivers of total function: the fact that increasing niche difference always eventually increases total yield enough to allow transgressive overyielding (Figure 2a) must necessarily be attributed to its positive effect on complementarity, not to its inconsistent effect on selection (Figure 3a). Similarly, the tendency of increasing fitness ratio in favor of species 1 to promote overyielding (Figure 2b) must be attributed to the selection effect, which always increased in this scenario, not complementarity, which decreased when fitness ratio rose above 1 (Figure 3b).

In contrast to previous investigations focusing on quantitative differences in the magnitudes of the components of the modern coexistence and additive partition frameworks (Carroll et al. 2011; Loreau, Sapijanskas, et al. 2012), our analysis shows qualitative correspondence between changes in each set of metrics. Indeed, studies aiming to identify mechanisms underpinning productivity changes along environmental gradients (e.g., Fridley 2002; Q.-G. Zhang and D.-Y. Zhang 2006) have focused predominantly on the sign of changes in additive partition or modern coexistence components, which we have shown are compatible between the two frameworks. Accordingly, in one of the few studies to compare the frameworks, Godoy, Gómez-Aparicio, et al. (2020) manipulated water availability for an annual plant community and found that niche differences and complementarity tended to simultaneously increase, while large fitness differences were associated with large selection effects.

#### 3.3 Predicting different outcomes using niche, fitness, and function

Loreau and Hector (2001)’s partition was developed to generalize different metrics studied by biodiversity experiments (e.g., relative yield total: de Wit 1960; transgressive overyielding: Schmid et al. 2008; approaches meant to address sampling effects: Loreau 1998), which either closely correspond to additive partition components (e.g., relative yield total is simply scaled complementarity: Loreau and Hector 2001) or can be obtained through an appropriate choice of expected relative yields RY_*E*,*i*_. We show here that our components (stabilizing niche difference, fitness–function relationships, and functional equalization) similarly generalize to a variety of outcomes and metrics considered in the literature (Appendix S4).

To demonstrate this generalizability, Box 2 highlights the condition for overyielding relative to the average intrinsic yield 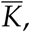, which also corresponds to the case used to analyze the additive partition. We graphically analyze this condition by visualizing it in the niche– fitness space alongside the condition for transgressive overyielding (Figure 3d). These requirements are less stringent than those for transgressive overyielding: for instance, equalizing fitness difference towards *f*_1_/ *f*_2_ = 1 always allows a coexisting community to overyield the average intrinsic yield (proven generally in Appendix S4). Nonetheless, the two conditions are closely related—in fact, the boundaries become parallel at high enough niche difference, and the distance between the boundaries is determined by imbalance in intrinsic yield (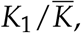, as opposed to *K*_1_/*K*_2_ for transgressive overyielding: Appendix S4). Put simply, all forms of overyielding require stabilization and a fitness advantage for the higher yielding competitor in excess of that required for coexistence alone, and the extent of this excess requirement is determined by the degree of imbalance in intrinsic yield.

Broadly speaking, we suggest that functional coexistence theory offers a generalizable theory for addressing the total function of communities. While the previous results of Carroll et al. (2011) focused on expressing the complementarity and selection components in terms of the niche and fitness difference, our focus is on using niche and fitness to provide quantitative predictions of total function. Accordingly, the present framework can predict of arbitrary forms of overyielding, as well as the conditions that maximize total function (as discussed above: equation 8, Figure 1b). Addressing previous warnings (e.g., Loreau, Sapijanskas, et al. 2012) that niche and fitness measures may not be suitable as quantitative predictors because they do not provide information about yield, functional coexistence theory incorporates intrinsic yield as the missing link enabling modern coexistence theory to satisfactorily predict species’ contribution to function (equation 4).

### 4 Linking functional coexistence theory to biological mechanism

Mechanistic models of competition offer a way towards a more complete understanding of ecosystem dynamics. In particular, the well-studied consumer–resource models provide an opportunity to unify community and ecosystem dynamics (Chase and Leibold 2003; Tilman 1982). Long used to elucidate the role of the niche in species coexistence (MacArthur 1970), these models predict the dynamics of competing species by representing their interactions with a set of shared resources (or limiting factors). From an ecosystem perspective, consumer–resource models often reflect fundamental constraints on nutrient cycling, resulting in more realistic dynamics (Gross 2008); from a community perspective, they can succinctly capture species dynamics using a minimum of measurements or parameters (Letten and Stouffer 2019). Therefore, such models may be able to simultaneously explain the composition and function of diverse communities, and have been applied to explore links between the biodiversity–ecosystem paradigm and other theories of biodiversity (Cardinale, Hillebrand, et al. 2009; Carroll et al. 2011; Turnbull et al. 2013).

In this section, we show how to quantitatively apply the functional coexistence framework to summarize mechanistic models, directly identifying biological mechanisms driving biodiversity effects. Using a trait-based model where species interfere with other species’ ability to use a limiting resource (Box 3), we show how species traits determine the niche, fitness, and function components of our framework and affect ecosystem function; in particular, we predict that resource level should not enable transgressive overyielding (subsection “Applying functional coexistence theory to a mechanistic model”). Using data from a classic plant competition experiment across a soil nitrogen gradient (Wedin and Tilman 1993), we fit the model and confirm our predictions regarding overyielding (subsection “Explaining productivity in a classic plant competition experiment”).

#### Box 3.

Identifying mechanisms for diversity effects in a consumer-resource model

We generalize a one-resource competition model from Tilman (1980) to an arbitrary number of species. Assuming that there is a single primary limiting factor *R* in the system, we then allow species to interfere with each other’s resource uptake in order to implicitly capture the effect of additional limiting factors. We consider *n* species, each with biomass *N*_*i*_, and a single limiting resource *R* (Figure 4a). The dynamics of the general model are given by the following equations:

**Figure 4:**
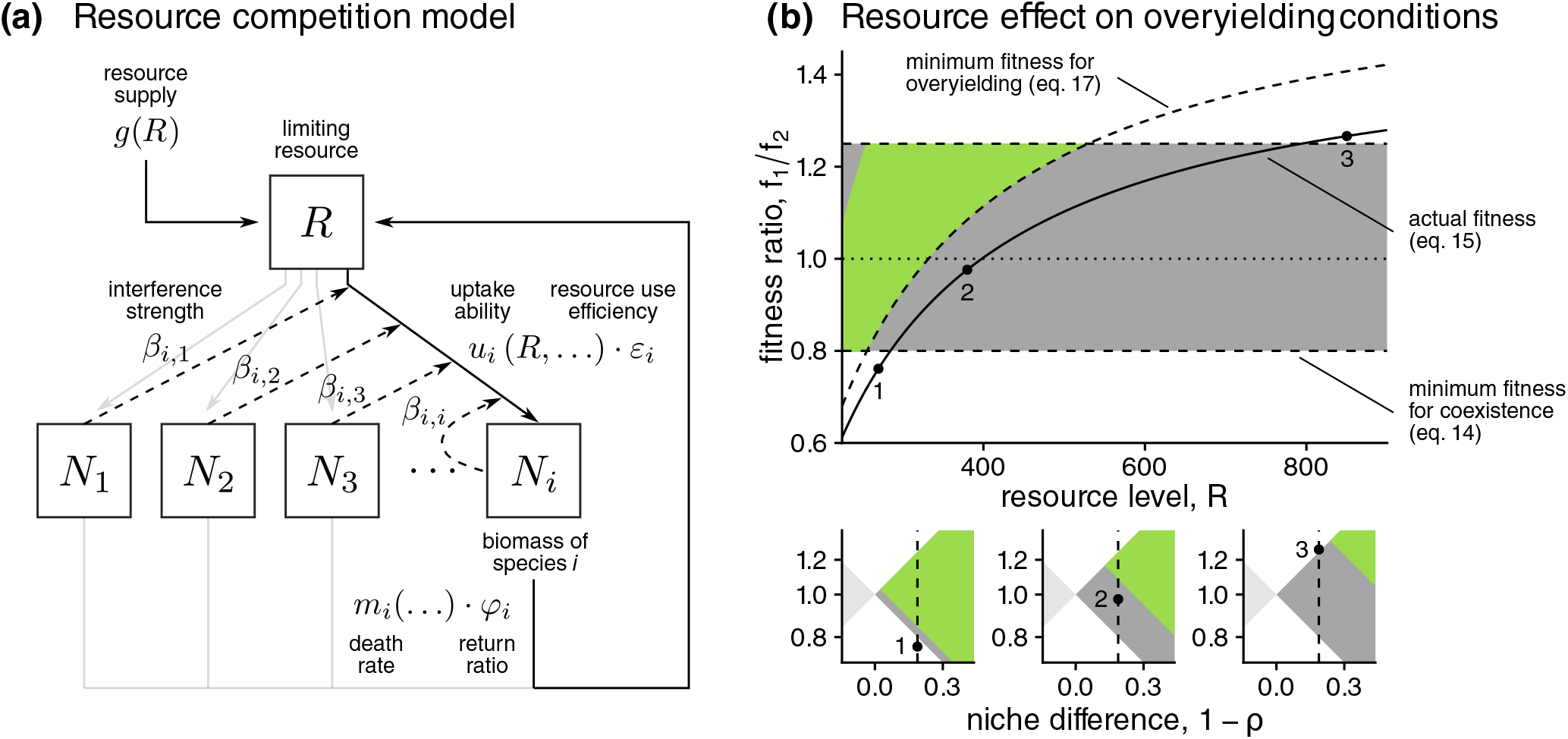
Applying the framework to a mechanistic competition model. We apply the functional coexistence framework to a general trait-based resource competition model. **(a) Model diagram**. A single limiting resource *R* is taken up by species *N*_1_, *N*_2_, *N*_3_, …, *N*_*i*_ differing in their uptake ability *v*_*i*_ and resource use efficiency *ε*_*i*_, while species-specific mortality μ_*i*_ returns resources to the pool. Furthermore, species interfere with each other, reducing their ability to take up resources according to interference strength *β*_*ij*_, which captures limitation by factors not explicitly represented in the model. **(b) Model predictions: changing resource level cannot drive overyielding**. For representative parameter values (Appendix S9), we show in the top panel how changing resource level affects fitness (solid line). In order to link this to coexistence outcomes, we shade the ratios at which coexistence (gray) and transgressive overyielding (green) could occur at each resource level. The minimum fitness ratio at which overyielding is possible (dashed curve) increases as resource level increases, while the conditions for coexistence (dashed horizontal lines) do not change. Regardless of resource level, the actual fitness ratio is never sufficiently high to allow overyielding. In the lower panels, we show how the communities at resource levels indicated 1–3 in the top diagram can be visualized in the niche and fitness space of Figure 1: points represent the actual niche and fitness difference, and the green region represents the changing requirements for overyielding.

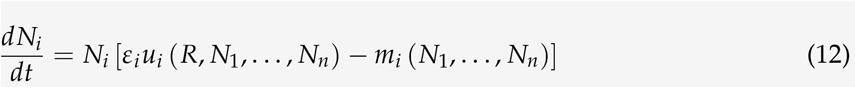

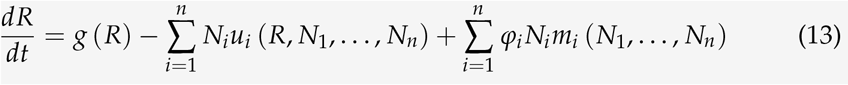

Here, a species’ growth depends on its resource use efficiency *ε*_*i*_ and its per-capita resource uptake *u*_*i*_ (a function of the abundance of the resource and of other species), and it experiences mortality according to some function *m*_*i*_. Resource dynamics are governed by some resource supply function *g*, uptake by consumers, and return from dead biomass, where φ_*i*_ is the resource returned per unit of species *i*’s dead biomass.

##### Linking the model to functional coexistence theory

To link our consumer–resource model to the general results above, we analyze a specific version of the model where species species *i*’s resource uptake is reduced by interference: 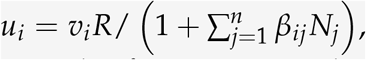, where *v*_*i*_ is *i*’s intrinsic uptake ability and *β*_*ij*_ is the strength of resource uptake interference by species *j* on species *i*; note that response to interference follows a functional form identical to that of competition in the Beverton–Holt model (Beverton and Holt 1957). Assuming constant species mortality (*m*_*i*_ = μ_*i*_) and a closed system (*g* (*R*) = 0) with complete resource return 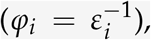, the total amount of resource in the system (i.e., in *R* and biomass) is constant and we can derive population dynamics as:

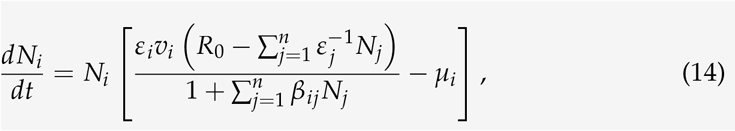

where the conserved quantity 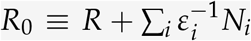 is the total amount of resource in the system (i.e., in the *R* pool or in biomass); we give the derivation in detail in Appendix S6 and show that it can also be interpreted as a first-order approximation to more complex resource dynamics. Since this form corresponds to the class of models considered in Box 1 and Appendix S1, we can derive the quantities necessary to apply functional coexistence theory (see full derivations in Appendix S6). Namely, we show that the coexistence components are

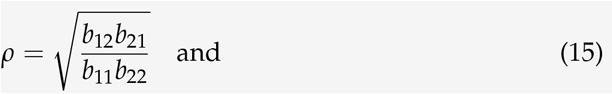

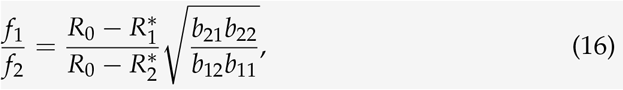

where 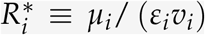 directly corresponds to Tilman (1980)’s *R*^*^, the minimum resource concentration at which species *i* can maintain positive population growth, and 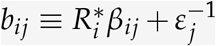 measures the competitive effect of species *j* on *i* via interference and the amount of resource it removes from the pool. On the other hand, intrinsic yield is

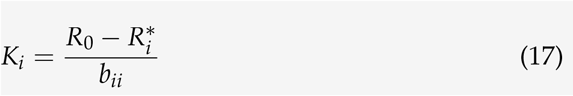

Note that *b*_*ij*_ is independent of total resource level, and thus a species’ actual sensitivity to competition (*sensu* Box 1) further depends on total resource level (Appendix S6), contributing the 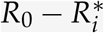 term to the expressions above.

##### Conditions for transgressive overyielding

Applying equation 7, we find straight-forwardly that transgressive overyielding requires

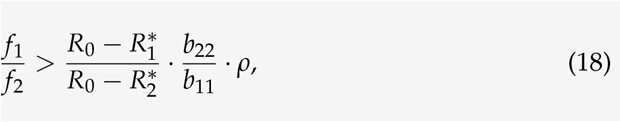

where only the first term depends on total resource level; in fact, the same ratio also determines the dependence of fitness on total resource level in equation 16. In other words, changing resource level always affects fitness ratio and yield ratio in the same way. Simplifying equation 18 to eliminate this factor shows that transgressive overyielding requires the higher yielding species to be more sensitive to conspecific than to heterospecific competitors (*b*_11_ *> b*_12_). In terms of the mechanistic traits of the model, this can be written

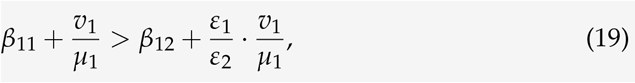

indicating that transgressive overyielding is favored if the higher yielding species experiences stronger interference from conspecifics (higher *β*_11_) or uses resources less efficiently (lower *ε*_1_).

#### 4.1 Applying functional coexistence theory to a mechanistic model

Defining a general trait-based resource competition model, we show that functional coexistence theory can be used to understand the drivers of ecosystem function in mechanistic models. In Box 3, we provide the model in mathematical detail and calculate the niche, fitness, and function measures needed to apply the functional coexistence framework. Closely related to previous models of interference competition (Amarasekare 2002) and facilitation (Gross 2008), our model (Figure 4a) considers an arbitrary number of species *N*_*i*_ competing for a single shared limiting resource *R*; in doing so, the model offers more mechanistic insight than classic phenomenological models of competition. Species differ in their ability *v*_*i*_ to obtain this resource, their resource use efficiency *ε*_*i*_, and in their mortality μ_*i*_, creating a competitive hierarchy in resource competition. Furthermore, species interfere with the resource uptake of conspecific and heterospecifics (*β*_*ii*_, *β*_*ij*_). Although the number of distinct resources limits the number of coexisting species in models of pure resource competition, this interference term allows an arbitrary number of species to coexist in the present model (Supplemental Figure S6.1). Stated conceptually, the limiting factors necessary for coexistence consist of the shared resource *R*, modelled mechanistically, and additional species interactions *β*_*ij*_, considered more phenomenologically. We suggest that this may be an appropriate mechanistic model for systems where species interact in diverse ways, but overall, interactions are strongly structured by competition for a single shared resource. For instance, in a plant system, *R* could represent space (e.g., in a forest ecosystem) or a limiting soil nutrient (e.g., nitrogen; Clark et al. 2018), while *β*_*ij*_ could represent more specific factors such allelopathy or shared pathogens (Ke and Wan 2020) that affect plants’ ability to compete for the shared resource. Similarly, in a fungal decomposer system, *R* could represent a common carbon substrate for which species compete, while *β*_*ij*_ could represent the effect of chemical interference (Tyc et al. 2017) or competition for other nutrients.

##### Modern coexistence theory: linking mechanistic and phenomenological perspectives on coexistence

Following the method of Letten, Ke, et al. (2017), we use the relationship between the mechanistic model and the Lotka–Volterra model to derive niche and fitness differences, thereby bridging mechanistic and phenomenological perspectives on coexistence. We find that the effect of competition can be broken down into two components. The first depends on resource level: exactly as in Tilman’s resource-ratio theory (Tilman 1982), resource competition ability is captured by the quantity 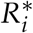 (i.e., the minimum resource level at which a species can maintain positive growth), and species *i*’s total sensitivity to competition is inversely related to 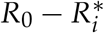 (i.e., the portion of the resource pool available to it; Appendix S6). The second component is a resource–independent quantity *a*_*ij*_ (Box 3), which measures the overall effect of one species on another through interference and monopolization of resources. Accordingly, as Letten, Ke, et al. (2017) found in a similar consumer–resource model, niche difference depends only on species traits (equation 15), but fitness difference also depends on total resource level (equation 16). In particular, by bringing the term 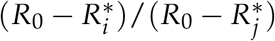 closer to unity, increasing total resource level reduces the importance of species’ differences in *R*^*^ and can act to equalize fitness differences in this model (Figure 4b, solid line), potentially allowing coexistence (gray region).

##### Functional coexistence theory: despite coexistence, changing resource level cannot drive overyielding

Applying our calculations for niche, fitness, and function, we can predict diversity–function relationships in the mechanistic model. Though each trait affects multiple components (Appendix S6), reflecting a general challenge in working with the components of modern coexistence theory (Barabás et al. 2018; Song et al. 2019), these components offer a considerably simpler picture of coexistence and its consequences. As an example, we calculate the effect of changing total resource level on the system (Figure 4b). As expected, doing so changes the fitness of the competing species (solid curve) but not the niche difference (horizontal dashed lines), driving a shift from the competitive exclusion of species 1 at low resource levels to coexistence or exclusion of species 2 at higher resource levels (Letten, Ke, et al. 2017). However, the system never shows transgressive overyielding (green): because increasing resource level increases a species’ intrinsic yield at the same time as its fitness, fitness never becomes higher than the condition imposed by yield imbalance (curved dashed line), preventing the system from entering the region where transgressive overyielding would occur (green). In fact, this is a fully general result for our model (Box 3; formulae in Figure 4b): provided species coexist, transgressive overyielding is determined solely by species’ intrinsic traits and varying total resource level cannot overturn the presence/absence of transgressive overyielding (equation 19). In the general terms of our functional coexistence framework, resource level can affect fitness, but its parallel effect on yield keeps the fitness–function relationship fixed.

##### Identifying biological mechanisms for overyielding

We have shown that functional coexistence theory provides a useful tool for predicting the outcome of the resource competition model. Going one step further, we highlight that it also clarifies the actual biological mechanisms for these outcomes. For instance, an extensive body of work (Fridley 2002; Godoy, Gómez-Aparicio, et al. 2020; Ratcliffe et al. 2017; Turnbull et al. 2013; Q.-G. Zhang and D.-Y. Zhang 2006) has sought to identify whether changes in resource limitation can explain variation in diversity–function relationships. However, our model suggests that resource limitation alone cannot change the relationship between fitness and intrinsic yield unless other forms of competition also change (i.e., our model’s *β*_*ij*_). Thus, we suggest that contrasting findings regarding the effect of resource gradients on diversity–function relationships can be reconciled by understanding that these results reflect changes in the nature of competition, which may be system specific, rather than some general effect of resource limitation itself. Meanwhile, another classic question from the biodiversity–ecosystem function literature concerns the apparent rarity of transgressive overyielding (Schmid et al. 2008), especially given theoretical expectations that the general condition should not be highly restrictive: higher functioning species should be more limited by conspecific than by heterospecific competitors (Loreau 2004). Translating the condition in our model to specific conditions on species traits (equation 19), we identify one potential mechanistic explanation: the same traits that confer high intrinsic yield (e.g., low sensitivity to conspecific interference *β*_*ii*_ and high resource use efficiency *ε*_*i*_) tend to make species less limited by conspecifics (e.g., *b*_*ii*_ and the left-hand side of equation 19).

#### 4.2 Explaining productivity in a classic plant competition experiment

In order to demonstrate how functional coexistence theory can help integrate theory and experiment, we test the theoretical predictions of our resource competition model by fitting our resource competition model to biomass data from an experiment quantifying plant competition across a soil nitrogen gradient (Figure 5). Working in an extensively studied grassland system (Cedar Creek, Minnesota, USA), the classic study of Wedin and Tilman (1993) competed four pairs of grass species while experimentally manipulating soil nitrogen, the nutrient shown to limit productivity in this system. We selected this study because it directly manipulated limiting resources (corresponding to *R*_0_ in our model); furthermore, extensive mechanistic data collected by the authors alongside their competition experiment provide an opportunity to validate our biological inferences. We applied the functional coexistence framework to investigate overyielding between the only species pair that showed robust coexistence: the grasses *Poa pratensis* and *Agropyron repens*. Using measurements of the species’ biomass production in monocultures, we first parameterized each species’ *R*^*^ and the resource–independent intraspecific interaction parameter *b*_*ii*_; next, since detailed time series data was not available, we fit the resource– independent interspecific interaction parameter *b*_*ij*_ to biomass in competition treatments. We then used the fitted parameters to quantify transgressive overyielding and the niche, fitness, and function measures (Figure 5; full methods and parameter fits in Appendix S7).

**Figure 5:**
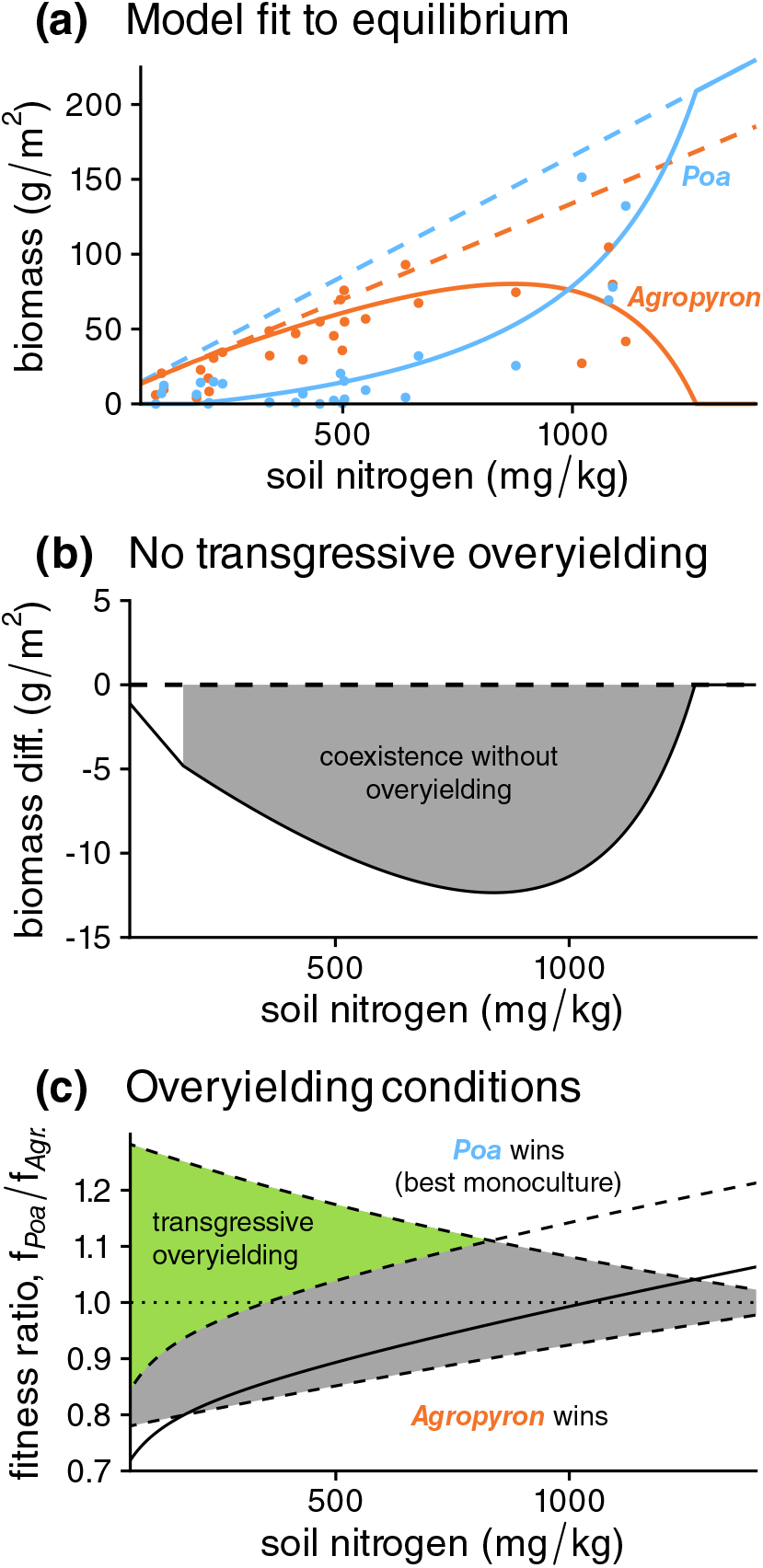
Applying functional coexistence theory to a plant competition experiment. We parameterize our resource competition model and identify drivers of pairwise community biomass using experimental data: Wedin and Tilman (1993) competed two grass species, *Poa pratensis* and *Agropyron repens*, across a soil nitrogen gradient (horizontal axis, all panels). Detailed methods are given in Appendix S7. **(a) Fitting the resource competition model**. After determining *R*^*^ and monoculture biomass of *Poa* (blue dashed line) and *Agropyron* (orange dashed line) from single-species growth, we fit our model to the plot-level equilibrium biomass of each species (points) across the soil nitrogen gradient. Model predictions (solid lines) capture the shift between *Agropyron* and *Poa* as nitrogen increases. **(b) Competitive effect on yield**. Using the fitted parameters, we calculate the equilibrium of the pairwise community and quantified transgressive overyielding (black line) as the difference between the community’s biomass and that of its highest yielding species (*Poa*); we shade the portion of this curve where the outcome was coexistence without transgressive overyielding (gray). **(c) Niche and fitness components**. We show the predictions of functional coexistence theory for this system, calculating the range of fitness ratios (vertical axis) that would allow transgressive overyielding (green) or just coexistence (gray) across the nitrogen gradient. The solid line shows the actual fitness ratio between the species; the dashed lines show the three boundaries as in Figure 4b.

##### Resource model captures yield and competitive outcomes

The model provided a close fit to monoculture yields, showing that *Poa* had higher yield than *Agropyron* (Figure 5a, dashed lines; Supplemental Figure S7.1), and that increasing nitrogen availability amplified this difference; however, species differed little in *R*^*^ (Supplemental Table S7.1). Model fits successfully predicted changes in competition biomass along the nitrogen gradient (Figure 5a, solid lines), though we found evidence that *Poa*’s sensitivity to competition from *Agropyron* (*a*_*Poa*,*Agr*._) intensified with increasing nitrogen (Supplemental Figure S7.2), a departure from the theoretical derivation in Box 3. Following observed shifts in biomass with increasing nitrogen, our model predicts a shift from competitive exclusion by *Agropyron* to coexistence with increasing dominance by *Poa*. However, this competitive shift towards the higher yielding species did not enable transgressive overyielding at any nitrogen level (Figure 5b).

##### Explaining lack of overyielding using functional coexistence theory

We explain this finding using the functional coexistence components in Figure 5c, which visualizes the fitness ratios enabling coexistence (gray) and transgressive overyielding (green) across the nitrogen gradient. The competitive shift was explained by an equalizing effect of resource availability: higher soil nitrogen increased the fitness ratio in favor of *Poa* (Figure 5c, solid line). While stabilizing niche differences would have been sufficient for transgressive overyielding at low nitrogen (green region; < ca. 700 mg/kg), the fitness–function relationship was far from optimal: the higher yielding *Poa* was competitively inferior under these conditions (solid line). Although increasing nitrogen favored *Poa*, it simultaneously amplified imbalance in the species’ intrinsic yields, thus decreasing the potential for overyielding (vertical distance of the green region). This closely corresponds to the predictions by our theoretical analysis (as simulated in Figure 4b): varying soil nitrogen did not change the relationship between actual fitness and the overyielding boundary (solid and dashed curved lines). We therefore conclude that at Cedar Creek, *Poa* lacks the excess niche difference and fitness advantage which would have allowed transgressive overyielding when it competes with *Agropyron*.

Complementing the insights available from other methodologies (e.g., selection and complementarity: Supplemental Figure S7.3), our functional coexistence analysis clarifies how competitive processes underpin the lack of transgressive overyielding in this system. Indeed, the mechanistic measurements from Wedin and Tilman (1993) indicated high similarity between *Poa* and *Agropyron*, both in terms of *R*^*^ (independently estimated by measuring ability to draw down soil nitrogen) and in resource use traits, providing ecological context for our finding that the system lacked the excess niche difference required for transgressive overyielding. Furthermore, though the dataset did not allow us to directly fit underlying resource use parameters, the authors’ independent finding that the species had a similar ability to draw down soil nitrogen corroborates our model’s *R*^*^ fits. This suggests that differences in intrinsic yield may have been driven by *Poa* experiencing less self-limitation from other factors (corresponding to lower *β*_*Poa*,*Poa*_), or by it producing more biomass from available nitrogen (higher *ε*_*Poa*_), both of which we predicted should prevent transgressive overyielding (equation 19). Thus, we highlight that, in tandem with manipulative experiments, our functional coexistence approach can identify the biological mechanisms responsible for changes in community function.

### 5 Beyond classic theory: applying functional coexistence theory to multiple functions and species

Thanks to conceptual synthesis within each field, literature on diversity–function relationships and on coexistence has addressed increasingly sophisticated questions. By integrating these two fields, our functional coexistence framework is also poised to address these contemporary research questions. Accordingly, we show that, because it explicitly predicts how competition affects individual species, functional coexistence theory can be applied to understand drivers of ecosystem multifunctionality (subsection “Highlighting the importance of niche difference for multifunctionality”). Meanwhile, we show that the niche, fitness, and productivity components we investigated in the pairwise case also provide information on function in multispecies communities, highlighting its potential as a unifying theory for biodiversity studies (subsection “Niche, fitness, and function in multispecies communities”)

#### 5.1 Highlighting the importance of niche difference for multifunctionality

Though our derivations and examples focus on processes promoting biomass production, we stress that the results of functional coexistence theory can apply to any ecosystem function (e.g., nutrient cycling: Godoy, Gómez-Aparicio, et al. 2020 or other ecosystem services: Hooper, Chapin III, et al. 2005), as we prove in Appendix S8. Moreover, going beyond previous approaches, functional coexistence theory can consider these functions simultaneously, allowing it to address an emerging synthesis considering biodiversity’s effect on *multifunctionality*, the ability for ecosystems to maintain multiple processes or services (Hector and Bagchi 2007). Accordingly, we apply our results to emphasize that just as it promotes individual functions, niche difference is also indispensable for ecosystem multifunctionality. To show this, we begin by noting that since equation 4 for species’ biomass contributions to the community can be multiplied by function per unit biomass at equilibrium φ_*i*_ to obtain functional contribution, the quantitative results of the framework can be generalized by considering Φ_*i*_ = *K*_*i*_·φ_*i*_ instead of *K*_*i*_. Under the assumption that function per unit biomass is constant, Φ_*i*_ is simply a species’ intrinsic yield *in terms of function*, instead of *biomass* yield.

With this extension, we can now consider conditions for simultaneous overyielding. In Figure 6, we add a second function (e.g., litter decomposition) to our biomass simulations (Figures 1–2) and consider the conditions promoting transgressive overyielding for both functions. In particular, we consider the case where the species follow a tradeoff between the two functions: in isolation, species 1 produces more biomass but species 2 has a higher level of the other function (*K*_1_/*K*_2_ *>* 1, but Φ_1_/Φ_2_ < 1). Accordingly, each function is maximized at a different fitness ratio (Figure 6a). Nonetheless, stabilization and equalization remain important for multifunctionality: though Figure 6b shows that lower niche difference values (1 − *ρ* = 0.15 to 0.29) only allowed transgressive overyielding for one function that corresponds to the competitively dominant species, higher niche difference (1 − *ρ* = 0.36) allowed simultaneous transgressive overyielding for both functions. Put conceptually, since competitive outcomes favor functions associated with fitter species, communities may display the same functional tradeoffs as their component species. However, niche differences in excess of those required for coexistence can overcome these tradeoffs, allowing communities to outperform individual species across multiple functions.

**Figure 6:**
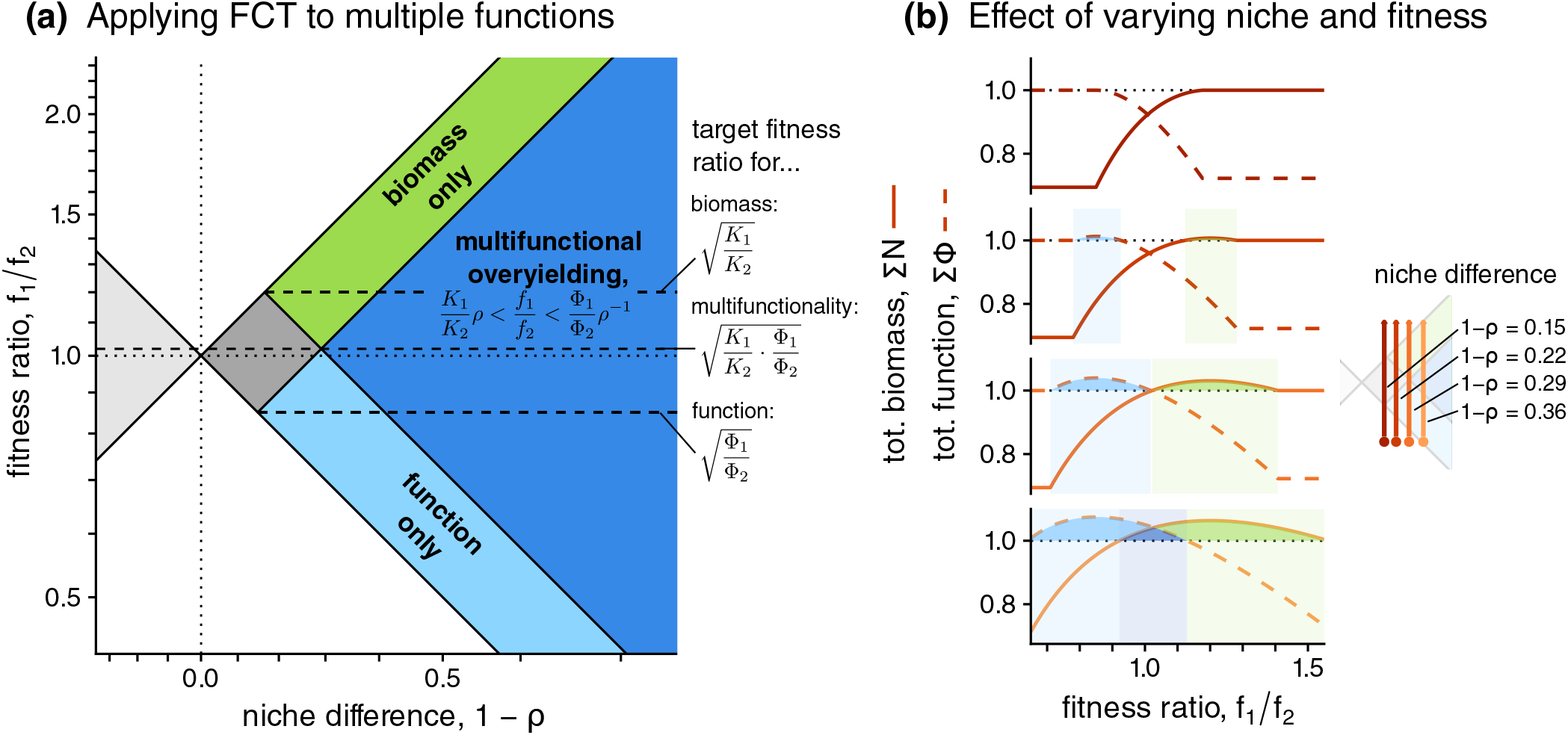
Applying the framework to multiple ecosystem functions. Using the same parameter values as Figures 1–2, we also allow the species to differ in a second function such that Φ_1_/Φ_2_ = 0.72/1, favoring species 2, as opposed to biomass where species 1 has higher intrinsic yield (*K*_1_/*K*_2_ = 1/0.694). **(a) Applying functional coexistence theory to predict multiple functions**. As in Figure 2a, we show niche–fitness combinations where species coexist without showing transgressive overyielding (dark gray region) and where species transgressively overyield in terms of biomass (green). We additionally show the new possibilities when considering a second function: transgressive overyielding in terms of that function only (light blue) or simultaneously for both functions (dark blue). We also indicate the target fitness ratio values for each form of overyielding, as derived in Appendix S1. **(b) Effect of varying niche difference and fitness ratio**. Each subpanel shows the effect of fitness ratio (horizontal axis) on biomass (solid line) and function (dashed line) under different values of niche difference, also indicated by line color; the inset shows values of niche difference and trajectories in the niche–fitness space. At lower niche differences, only one form of transgressive overyielding is possible (light blue or green shading), but at the highest niche difference, the community can simultaneously overyield in terms of both functions (dark blue shading). See Appendix S9 for parameter values.

Though it has been suggested that communities consisting of species performing different functions should show multifunctionality (Hector and Bagchi 2007), functional coexistence theory shows that this depends on stabilization and equalization between these species. Our more general theoretical analysis (summarized for two functions in Figure 6a and given in full in Appendix S8) clarifies that outcomes depend on the pair of functions showing the strongest tradeoff (i.e., with the most dissimilar yield ratios): the stronger the tradeoff between functions, the more stabilization is required for multifunctionality. More specifically, transgressive overyielding for multiple functions is possible when niche differences provide strong enough stabilization to overcome this dissimilarity, and when the fitness ratio is sufficiently equalized (i.e., close enough to the geometric mean of these two yield ratios). Indeed, in an experimental test of the relationship between coexistence components and multiple ecosystem functions, Godoy, Gómez-Aparicio, et al. (2020) found that high niche difference and similarity in fitness increased both biomass production and litter decomposition rate in diverse plant communities, emphasizing the importance of excess niche difference for multifunctionality. Agreeing with these empirical findings, our results shed light on the general importance of stabilization and equalization for ecosystem function.

#### 5.2 Niche, fitness, and function in multispecies communities

Previous debates (Loreau, Sapijanskas, et al. 2012) have highlighted a key limitation of modern coexistence theory for understanding biodiversity effects: typically, the theory has focused on small communities, in contrast to the larger numbers of species considered in many biodiversity experiments (Hooper, Chapin III, et al. 2005) and in real systems. Fortunately, recent theory increasingly provides tools for understanding the maintenance of diversity in multispecies communities (Barbier, Arnoldi, et al. 2018; Saavedra et al. 2017). We suggest that synthesizing functional coexistence theory with these emerging frameworks may allow it to address these questions. As a motivating case study, we show that our pairwise metrics also predict total function in multispecies communities.

In Figure 7, we simulate the resource interference model from Box 3 for *n* = 20 species, starting with a reference community where each mechanistic trait was drawn from a random distribution (starred points; parameter values in Appendix S9). To investigate the role of our functional coexistence components, we varied the interspecific interference parameter *β*_*ij*_ in order to vary the median niche difference (i.e., stabilization; Figure 7a) or median fitness advantage of the higher yielding species (fitness–function relationship; Figure 7b). Additionally, we manipulated functional equalization (Figure 7c) by varying the *β*_*ii*_ to change the range of intrinsic productivities in the system, while keeping the maximum fixed (since it is the reference point for transgressive overyielding). Detailed methods for these manipulations are given in Appendix S9.

**Figure 7:**
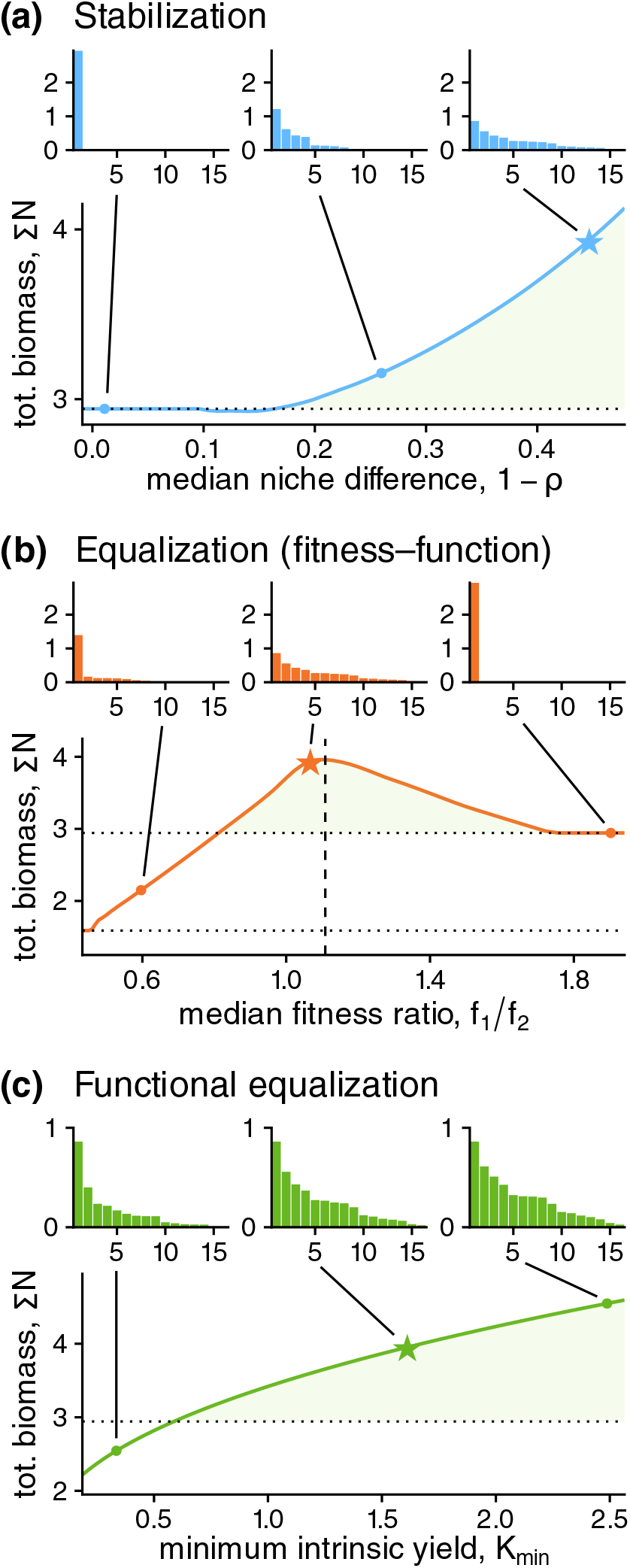
Functional coexistence components in multispecies communities. We consider a multispecies community (*n* = 20) under the mechanistic resource competition model in Box 3, with traits randomly drawn from statistical distributions (details in Appendix S9). Starting from this reference community, we vary the interference terms *β*_*ij*_ in order to manipulate **(a) stabilization**, i.e. the median pairwise niche difference, **(b) fitness–function relationship**, i.e. the median pairwise fitness ratio for a higher yielding species versus its competitor, or **(c) functional equalization**, i.e. the minimum intrinsic yield. Each main panel shows the total biomass of the community as the strength of the component is varied; the green region highlights transgressive overyielding (relative to the highest yielding monoculture, indicated with a dotted line). For each scenario, we show the community’s rank abundance curve (i.e., plotting species rank on the horizontal axis against abundance on the vertical axis) for three representative points. We indicate the reference community, which is the same in all three panels, with a star.

We quantified the effect of each process on equilibrium community structure (rank abundance curves shown in insets) and transgressive overyielding (green shaded areas in main panels). Stabilization played a similar role in the two-species and multispecies models (Figure 7a). Sufficient niche difference was required for species coexistence (panel a), but just as in the two-species case, values just enough to enable coexistence actually slightly decreased total biomass (more clearly visualized in Supplemental Figure S9.2). Nonetheless, as before, further stabilization was able to create transgressive overyielding (point 3). Meanwhile, we also confirmed the effects of the fitness–function relationship (Figure 7b). The traits in our reference model (point 2) created a positive fitness–function relationship for most species pairs (Supplemental Figure S9.1), allowing overyielding. Slightly increasing pairwise fitness ratios in favor of more productive species (dashed line: median *f*_1_/ *f*_2_ = 1.10) resulted in the highest total biomass. Meanwhile, more imbalanced fitness ratios resulted in the extinction of species from the system and reduced total biomass (points 1 and 3), resulting in competitive dominance by the lowest or highest yielding species. Finally, we found an analogous role for functional equalization (Figure 7c): bringing the minimum intrinsic yield (horizontal axis) closer to the maximum (horizontal dashed line) allowed transgressive overyielding in the system; we also confirmed this was the case when the median, not the maximum, intrinsic yield was kept constant (Supplemental Figure S9.4).

These results confirm the close link between coexistence and total function, and suggest that the measures predicting transgressive yielding in two-species systems also provide useful context for understanding multispecies systems. Indeed, there was approximate quantitative correspondence between pairwise metrics and multispecies outcomes: for instance, in the equalizing scenario (Figure 7b), coexistence of two or more species occurred when pairwise fitness was approximately within the range determined by median niche difference (Supplemental Figure S9.3) and total biomass was optimized near the value predicted from median pairwise yield ratio (median 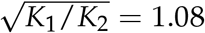; Supplemental Figure S9.1). This finding broadly agrees with other studies finding correspondence between pairwise modern coexistence theory measures and overall community outcomes (Advani et al. 2018; Carroll et al. 2011; Godoy, Gómez-Aparicio, et al. 2020). Like Carroll et al. (2011), who provided a similar multispecies analysis using the non-mechanistic Lotka– Volterra model, we emphasize the importance of stabilization for promoting overyielding; addressing the critique of Loreau, Sapijanskas, et al. (2012), we also clarify the importance of role of fitness and productivity variation, which were not considered there.

Despite the insight offered by functional coexistence theory into the results of our multispecies simulations, we caution that the calculations presented here do not yet offer a quantitative framework directly predicting multispecies yield as it does in the two-species case. Nonetheless, the pairwise theory may offer a valuable starting point from which recently developed multispecies theory can build (Advani et al. 2018; Barbier and Arnoldi 2017; Saavedra et al. 2017). Indeed, our pairwise result that competitive fitness and intrinsic yield jointly predict species’ contributions to the community (i.e., equation 4) also holds approximately for these simulations (Supplemental Figure S9.5), as it does for several theoretical multispecies models (Advani et al. 2018; Gibbs et al. 2022). Accordingly, we anticipate a growing role for multispecies theories of species coexistence in providing a predictive quantitative synthesis between community ecology and ecosystem function.

### 6 Conclusion: towards mechanistic understanding of biodiversity–function relationships

By showing fundamental links between modern coexistence theory and ecosystem function, our findings link community and ecosystem processes. We show that a simple condition predicts when coexistence increases the total function of a community: species must experience niche differences and fitness advantages in excess of those required for coexistence. Thus, our theoretical framework, which we term functional coexistence theory, explicitly identifies three processes that explain biodiversity–function relationships: stabilizing niche differences, fitness–function relationships, and functional equalization, which we demonstrate can be applied to mechanistic models and experimental data, and to multiple ecosystem functions and species.

By demonstrating the compatibility of the components of modern coexistence theory with the additive partition from the biodiversity–ecosystem function literature, our work adds to a growing shift from particular metrics to a focus on the conceptual processes encoded by these metrics (Godwin et al. 2020; Loreau and Hector 2019). For instance, recent work has highlighted that, despite apparent quantitative disagreement, different formulae for the components of modern coexistence theory generally encode shared intuition regarding how biological processes affect species’ abilities to persist (Godwin et al. 2020; Spaak, Ke, et al. 2023). Similarly, we found that complementarity measures the same conceptual process as niche difference: reduction in the amount of competition species experience from heterospecifics, quantified using the invasion growth rate (our *F*_*i*_) as a “common currency” (Box 2, Appendix S5; Grainger et al. 2019). Our findings take advantage of a more general relationship between invasion growth rate and species’ contributions to the community (Appendices S1 and S3; Arnoldi et al. 2022; Gibbs et al. 2022), which provides a general conceptual foundation for the link between coexistence and function, and allows for further quantitative extensions of the framework.

Like modern coexistence theory itself (Godwin et al. 2020), our functional coexistence framework relies on information about competitive processes which may not be available in all systems. Thus, while it offers new insights for testing the hypothesis that coexistence-promoting processes are integral to biodiversity–function relationships, it may not be applicable to the wide range of systems and datasets covered by previous quantitative methods (Bannar-Martin et al. 2018; Fox 2005; Loreau and Hector 2001). Moreover, as previously noted (Loreau, Sapijanskas, et al. 2012), these empirically-motivated methods differ fundamentally in scope from the modern coexistence theory research program and its more theoretical aims: the additive partition, for instance, aims to explain differences observed over the course of a study (which may not correspond to equilibrium dynamics; Wagg et al. 2019), while coexistence theory considers an abstractly defined system and its long-term trajectory (i.e., the stable equilibria or attractors of a system; Barabás et al. 2018).

Nonetheless, we expect that the two approaches will be complementary as considering competition is necessary in order to understand how ecosystem function may change with time or environmental context (Wan and Crowther 2022). To do so, our functional coexistence theory integrates the predictive framework of modern coexistence theory with information regarding differences in species’ intrinsic yield (i.e. level of function). Previous work has often cited yield variation as a methodological consideration for biodiversity– function experiments (de Wit 1960; Schmid et al. 2008). However, much as the stabilizing– equalizing framework highlighted differences in fitness as indispensable for predicting coexistence (Adler, HilleRisLambers, et al. 2007), our framework suggests that functional imbalance deserves increased attention in its own right as a predictor of ecosystem function. Furthermore, though it has long been noted that species with higher function in isolation may not perform better under competition (de Wit 1960; Gustafsson 1951; Montgomery 1912), our framework’s focus on fitness–function relationships highlights the quantitative consequences such tradeoffs have for community function. Thus, as a growing functional paradigm in community and ecosystem ecology highlights (Clark et al. 2018; Treseder and Lennon 2015), identifying tradeoffs between competitive processes and functional outcomes may provide a route towards more generally predicting of ecosystem function.

More broadly, we echo recent suggestions that moving forward in biodiversity–function research requires searching for the shared mechanisms that structure both communities and ecosystems (Hooper, Chapin III, et al. 2005; Loreau 2010; Mayor et al. 2024). The framework presented here bridges the questions of biodiversity–function literature with the rich theoretical foundations of the modern coexistence theory literature. Indeed, studies in hundreds of systems have quantified niche and fitness differences (Buche et al. 2022) and attributed them to specific biological mechanisms (e.g., Yan et al. 2022), often finding stabilizing and equalizing forces in excess of the requirements of coexistence (Adler, Ellner, et al. 2010; Buche et al. 2022; Levine and HilleRisLambers 2009). Our framework clarifies that these excesses—Adler, Ellner, et al. (2010)’s “embarrassment of niches”—should work to maximize the total functioning of a community. As empirical work increasingly seeks to identify the specific biological mechanisms driving ecosystem function, the modern coexistence literature can thus offer a valuable starting point (Godoy, Gómez-Aparicio, et al. 2020; Wang et al. 2024). Accordingly, we emphasize the utility of ecological theory for addressing today’s pressing challenges. By integrating established theory from community and ecosystem ecology, we repurpose well-studied tools in order to provide a fundamental understanding of the relationship between coexistence and ecosystem functioning. Adding to a growing synthesis of ecological theory across scales to address anthropogenic environmental change (Mayor et al. 2024; Wan and Crowther 2022), we hope the functional coexistence framework presented here will help build a more predictive understanding of Earth’s ecosystems and their roles in a changing world.

## Acknowledgments

We thank Tom W. N. Walker, Marcel van der Heijden, and Jukka Jokela for their comments on the study and for discussions that helped clarify our framing of the project. Additionally, we would like to acknowledge Ching-Lin Huang and Yu-Pei Tseng for feedback that helped us improve the clarity of the manuscript.

## Statements and Declarations

### Author Contributions

JW and TWC conceived the study. JW and IH designed the resource competition model, and JW and PJK designed the remaining analyses with input from all authors. JW wrote analysis code and derived the theoretical results. JW and PJK drafted the manuscript with equal contributions, and all authors contributed to revisions.

### Data Availability

No original data were presented in this manuscript. All code and inputs required for the simulations and re-analyses in this manuscript are publicly available at https://github.com/joe-wan/functional-coexistence-code/.

