## Supplemental Data 1 for "Functional coexistence theory: a mechanistic framework linking biodiversity to ecosystem function"

**S1 Overyielding in Lotka–Volterra and related models** **2**

**S2 Conditions for other forms of overyielding** **9**

**S3 Applying functional coexistence theory to nonlinear competition** **14**

**S4 Calculating the additive partition** **19**

**S5 Interpreting complementarity as a niche difference measure** **25**

**S6 Resource competition model with interference** **30**

**S7 Fitting the resource competition model to empirical data** **36**

**S8 Applying the framework to other ecosystem functions** **41**

**S9 Numerical simulations** **43**

### S1 Overyielding in Lotka–Volterra and related models

In this Appendix, we derive the conditions for overyielding for two-species systems obeying Lotka–Volterra or related dynamics. After first defining the class of models considered, we derive familiar results for yield (biomass) in isolation, equilibrium abundance, and coexistence conditions. We then use these results to identify the conditions that allow two species to show transgressive overyielding (as highlighted in the main text).

**Lotka–Volterra and related models** We consider the Lotka–Volterra model of competition, given by

$$\frac{dN_i}{dt} = r_i N_i \cdot \left(1 - \sum_{j=1}^{n} \alpha_{ij} N_j\right), \tag{S1.1}$$

where $N_i$ is the abundance of species $i$, $r_i$ is its intrinsic growth rate, and $\alpha_{ij}$ is the competitive effect of species $j$ on $i$, and $n$ is the number of species in the system. We consider the two-species case, corresponding to $n = 2$ and $i = 1, 2$. Though we use the Lotka–Volterra model for the derivations below, the method and results directly hold for any system where per-capita growth rates follow

$$h_i \left(1 - \sum_{j=1}^{n} \alpha_{ij} N_j\right), \tag{S1.2}$$

where $1 - \sum_j \alpha_{ij} N_j$ represents how growth is reduced due to competition, such that $h_i(1)$ represents no competition (i.e., the intrinsic rate of increase), and $h_i(0)$ represents the case where competition has completely suppressed population growth. Thus, we require that $h_i(\cdots)$ be any monotonically increasing function such that $h_i(0) = 0$.

The formulation of the Lotka–Volterra model used here corresponds to the simple choice of

$$h_i(x) = r_i x, \tag{S1.3}$$

i.e., simple scaling by the intrinsic growth rate $r_i$, but this class of models includes many other familiar models of competition. For instance, we can obtain the Beverton–Holt model (Beverton and Holt 1957) used to study annual plant competition (Levine and HilleRisLambers 2009) by defining

$$h_i(x) = \frac{x}{1 - x + (\lambda_i - 1)^{-1}}, \tag{S1.4}$$

where $h_i$ gives the per-capita growth between two time points, i.e., $N_i(t+1) = N_i(t) \cdot$

$\left(1 + h_i(1 - \sum_j \alpha_{ij} N_j(t))\right)$, and we can recover the original form of the equation by writing

$$\begin{aligned} N_i(t+1) &= N_i(t) \cdot \frac{1 + (\lambda_i - 1)^{-1}}{\sum_{j=1}^{n} \alpha_{ij} N_j + (\lambda_i - 1)^{-1}} \\ &= N_i(t) \cdot \frac{\lambda_i}{1 + (\lambda_i - 1) \sum_{j=1}^{n} \alpha_{ij} N_j} \\ &= N_i(t) \cdot \frac{\lambda_i}{1 + \sum_{j=1}^{n} \alpha'_{ij} N_j}, \end{aligned} \tag{S1.5}$$

where in the first line we have combined the 1 and fraction terms into one fraction, in the second line we have multiplied through by $\lambda_i - 1$, and in the last line we have let $\alpha'_{ij} = \alpha_{ij} (\lambda_i - 1)$, where $\alpha'_{ij}$ is the originally defined competition coefficient in this model.

**Yield in isolation, $K_i$** We calculate the yield of each species when it is growing alone. When all other $N_j$ are 0, the equilibrium abundance of $i$ is denoted $K_i$ (the carrying capacity) and can be found by solving the equilibrium condition associated with $i$'s growth rate (after setting all other abundances to zero, $1 = \alpha_{ii} K_i$), which gives the familiar expression for carrying capacity,

$$K_i = 1/\alpha_{ii}. \tag{S1.6}$$

**Conditions for coexistence: competitive fitness, $F_i$** Coexistence can be investigated using the invasion criterion, which states that species $i$ persists under competition if it has a positive invasion growth rate $\text{IGR}_i$, defined as the species' per-capita growth rate when its own abundance is near zero and all other species are at equilibrium with each other. For the two species system considered here, we consider each species' ability to invade an equilibrium community consisting only of its competitor; in the generic case (e.g. equation S1.2) this is

$$\text{IGR}_i = h_i \left(1 - \alpha_{ij} K_j\right) = h_i \left(1 - \alpha_{ij}/\alpha_{jj}\right). \tag{S1.7}$$

Noting that the sign of this quantity does not depend on the function $h_i$ (since by definition it does not change the sign of its argument), we can define a species' competitive fitness in this class of models in terms of $h_i^{-1}(\cdots)$, the inverse of the function $h_i$, as

$$F_i = h_i^{-1}(\text{IGR}_i) = 1 - \alpha_{ij}/\alpha_{jj}, \tag{S1.8}$$

such that coexistence between species 1 and 2 requires $F_1 > 0, F_2 > 0$, which are respectively the persistence conditions for species 1 and 2. Note that we denote this quantity with a capital $F$ to distinguish it from the fitness ratio $f_i/f_j$. Though less familiar than the fitness ratio for the models considered here, we retain both notations in order to simplify the calculations below. In fact, our $F_i$ is closely related to the relative reduction measure $S_i$ of Carroll et al. (2011), i.e., $F_i = 1 - S_i$. For the Lotka–Volterra model, it is also equivalent to choosing a scaling factor of $1/r_i$ when applying the general version of modern coexistence theory, as summarized by Barabás et al. (2018). We further discuss the relationship between $F_i$ and other metrics in Appendix S5.

**Conditions for coexistence: niche overlap $\rho$ and fitness ratio $f_i/f_j$** Noting that the persistence condition for species $i$ ($F_i > 0$) is equivalent to

$$1 - F_i < 1, \tag{S1.9}$$

we can measure the tendency of species to competitively exclude each other using the geometric mean of values of $1 - F_i$. This gives the familiar niche overlap metric for two species $i$ and $j$,

$$\begin{aligned} \rho &= \sqrt{(1 - F_i)\left(1 - F_j\right)} \\ &= \sqrt{\frac{\alpha_{ij}\alpha_{ji}}{\alpha_{ii}\alpha_{jj}}}, \end{aligned} \tag{S1.10}$$

where larger values indicate a greater tendency towards competitive exclusion; a similar formula was given by Carroll et al. (2011). Meanwhile, differences in fitness can be quantified as the degree to which values of $1 - F_i$ depart from the geometric mean; accordingly, we can define a fitness ratio between the two species as

$$\begin{aligned} \frac{f_i}{f_j} &= \frac{\rho}{1 - F_i} = \sqrt{\frac{1 - F_j}{1 - F_i}} \\ &= \sqrt{\frac{\alpha_{ji}\alpha_{jj}}{\alpha_{ij}\alpha_{ii}}}. \end{aligned} \tag{S1.11}$$

This also allows us to solve for $F_i$ in terms of niche difference and fitness ratio, giving

$$F_i = 1 - \rho\frac{f_j}{f_i}, \tag{S1.12}$$

which was also highlighted as equation 5 in the main text. We can rewrite the persistence condition for species $i$ in terms of $\rho$ and $f_i/f_j$ by taking the reciprocal of both sides of S1.9 and multiplying through by $\rho$, noting that the resulting expression is equivalent to $f_i/f_j > \rho$. Deriving the analogous expression for the persistence of $j$ and taking the reciprocal gives $f_i/f_j < \rho^{-1}$. Thus, the condition for coexistence in the system can be written as

$$\rho < \frac{f_i}{f_j} < \rho^{-1}, \tag{S1.13}$$

which requires that $\rho < 1$ (i.e., niche overlap is less than complete) in order for coexistence to be possible at some fitness ratios.

**Coexistence equilibrium,** $\hat{N}_i$ When the two species coexist, their equilibrium is given by the solution to the system of linear equations given by $1 = \sum_{j=1}^{n} \alpha_{ij}\hat{N}_j$ for all $i$, where we write $\hat{N}_i$ to denote abundance at the coexistence equilibrium. For the two-species system considered here, this corresponds to

$$\begin{aligned} 1 &= \alpha_{11}\hat{N}_1 + \alpha_{12}\hat{N}_2 \\ 1 &= \alpha_{21}\hat{N}_1 + \alpha_{21}\hat{N}_2, \end{aligned} \tag{S1.14}$$

where each equation represents the condition that one species' per-capita growth be zero. There are many equivalent ways to solve this equation; for convenience, we do so here by rewriting the system as a vector equation $\vec{1} = \mathbf{A}\vec{n}$, where

$$\mathbf{A} = \begin{bmatrix} 1 & \frac{\alpha_{12}}{\alpha_{22}} \\ \frac{\alpha_{21}}{\alpha_{11}} & 1 \end{bmatrix}, \vec{n} = \begin{bmatrix} \alpha_{11}\hat{N}_1 \\ \alpha_{22}\hat{N}_2 \end{bmatrix} = \begin{bmatrix} \hat{N}_1/K_1 \\ \hat{N}_2/K_2 \end{bmatrix}, \tag{S1.15}$$

and $\vec{1}$ is a vector where each element is one. Cramer's rule states that the $i$th element of the solution to this equation is given by

$$n_i = \frac{\det \mathbf{A}_i}{\det \mathbf{A}}, \tag{S1.16}$$

where $\mathbf{A}_i$ is the matrix obtained by replacing the $i$th column of $\mathbf{A}$ with $\vec{1}$. This gives

$$n_i = \frac{1 - \alpha_{ij}/\alpha_{jj}}{1 - \alpha_{ij}\alpha_{ji}/\alpha_{ii}\alpha_{jj}} = \frac{F_i}{1 - \rho^2}, \tag{S1.17}$$

where we have noted that the numerator and part of the denominator correspond to the expressions for $F_i$ and $\rho$ above. Then the actual abundance of the species is simply

$\hat{N}_i = K_i n_i$, which gives

$$\hat{N}_i = \frac{F_i K_i}{1-\rho^2}, \tag{S1.18}$$

stating that the abundance of each species is directly proportional to its competitive fitness $F_i$ and yield in isolation $K_i$, with a proportionality constant of $1/\left(1-\rho^2\right)$. The total abundance of the two species at equilibrium $\Sigma\hat{N}$ is simply

$$\Sigma\hat{N} = \hat{N}_1 + \hat{N}_2 = \frac{F_1K_1 + F_2K_2}{1-\rho^2}, \tag{S1.19}$$

which can also be expressed solely in terms of $F_1, F_2$ using $1-\rho^2 = 1-(1-F_1)(1-F_2) = F_1 + F_2 - F_1F_2$, giving

$$\Sigma\hat{N} = \frac{F_1K_1 + F_2K_2}{F_1 + F_2 - F_1F_2}. \tag{S1.20}$$

**Conditions for transgressive overyielding** Overyielding occurs when the species coexist and produce more biomass than some expected value. In this case, we consider *transgressive overyielding*, where the species together produce more biomass than the highest yielding species in isolation. Without loss of generality, assume that the higher yielding species is labelled species 1 (i.e., $K_1 > K_2$). Then transgressive overyielding occurs when $\Sigma\hat{N} > K_1$. Multiplying through by $1-\rho^2 = 1-(1-F_1)(1-F_2) = F_1 + F_2 - F_1F_2$, which is positive for stably coexisting species, we obtain

$$\begin{aligned} F_1K_1 + F_2K_2 &> K_1\left(F_1 + F_2 - F_1F_2\right) \\ F_1F_2K_1 &> F_2\left(K_1 - K_2\right) \\ F_1 &> 1 - \frac{K_2}{K_1}, \end{aligned} \tag{S1.21}$$

where in the last step we have divided through by $K_1$ (positive because it is a carrying capacity) and $F_2$ (positive because the species coexist). Thus, transgressive overyielding requires the higher yielding species to have sufficiently high fitness. To convert to the more familiar $\rho$ and $f_1/f_2$ metrics, we rearrange and take the reciprocal to obtain

$$\begin{aligned} \frac{1}{1-F_1} &> \frac{K_1}{K_2} \\ \sqrt{\frac{1-F_2}{1-F_1}} &> \frac{K_1}{K_2}\sqrt{(1-F_1)(1-F_2)} \\ \frac{f_1}{f_2} &> \frac{K_1}{K_2}\rho, \end{aligned} \tag{S1.22}$$

where in the second step we have multiplied through by $\rho = \sqrt{(1-F_1)(1-F_2)}$, which is positive for coexisting species. Since overyielding also requires that the fitness of species 1 is not so high that it excludes its competitor (i.e. that $f_1/f_2 < \rho^{-1}$), the complete condition for transgressive overyielding can be written

$$\frac{K_1}{K_2}\rho < \frac{f_1}{f_2} < \rho^{-1}, \tag{S1.23}$$

which is closely related to the coexistence condition (equation S1.13) except that the lower bound for fitness is multiplied by a factor of $K_1/K_2$ (which is greater than 1 since species 1 has the higher yield).

**Optimal yield** Keeping $\rho$ constant, we find the value of $f_1/f_2$ that maximizes total yield. This is equivalent to maximizing the numerator of equation $F_1K_1 + F_2K_2$, which we can further rewrite in terms of $1 - F_1$ and $\rho$ as

$$K_1 + K_2 - (1-F_1)K_1 - \frac{\rho^2}{(1-F_1)}K_2, \tag{S1.24}$$

where we have used the identity $\rho^2 = (1-F_1)(1-F_2)$. Setting the derivative of this expression with respect to $(1-F_1)$ equal to zero, we find the following condition for the optimum yield:

$$\begin{aligned} -K_1 + (1-F_1)^{-2}\rho^2 K_2 &= 0 \\ (1-F_1)^{-2}\rho^2 &= \frac{K_1}{K_2} \end{aligned} \tag{S1.25}$$

Since $(1-F_1)^{-2}\rho^2 = (1-F_2)/(1-F_1) = \left(f_i/f_j\right)^2$ and fitness is always positive, the optimal fitness ratio is simply

$$\frac{f_1}{f_2} = \sqrt{\frac{K_1}{K_2}}. \tag{S1.26}$$

Thus optimal yielding occurs when the higher yielding species in isolation has greater fitness to a precise degree (specified by the square root of the yield ratios). Since $K_1 > K_2$, we have that $1 < \sqrt{K_1/K_2} < K_1/K_2$; thus, this optimum fitness ratio satisfies the overyielding condition (equation S1.23) whenever niche overlap allows coexistence ($\rho < 1$). Furthermore, we can derive the optimal yield by expressing equation S1.19 using equation S1.12 and substituting $f_1/f_2 = \sqrt{K_1/K_2}$ to obtain that

$$\left(\Sigma\hat{N}\right)_{\max} = \frac{K_1 + K_2 - 2\rho\sqrt{K_1 K_2}}{1-\rho^2}. \tag{S1.27}$$

**Excess niche and fitness differences** The conditions for overyielding (equation S1.23) can be understood as requiring "excess" niche and fitness difference relative to the conditions for coexistence (equation S1.13). To see this, we can take the logarithm of each expression in equation S1.23 to obtain

$$\log\rho + \log\frac{K_1}{K_2} < \log\frac{f_1}{f_2} < -\log\rho \tag{S1.28}$$

Defining the quantity

$$\Delta = \frac{1}{2}\log\frac{K_1}{K_2} \tag{S1.29}$$

and subtracting it from each of the three expressions in the compound inequality, we find that

$$\log\rho + \Delta < \log\frac{f_1}{f_2} - \Delta < -\log\rho - \Delta. \tag{S1.30}$$

Noting that the outer expressions are additive inverses of each other, we can rewrite this as a single inequality using absolute values, finding

$$\begin{aligned} -\log\rho - \Delta &> \left|\log\frac{f_1}{f_2} - \Delta\right| \\ \mathrm{FD} - \Delta &> \left|\mathrm{ND} - \Delta\right|, \end{aligned} \tag{S1.31}$$

where we have noted that the logarithmic expressions $-\log\rho$ and $\log f_1/f_2$ correspond to the definitions of niche and fitness difference in Figure 1 of the main text. Thus, niche difference and the fitness advantage of the higher yielding species must both be in excess by an amount of $\Delta$ of the requirements for coexistence ($\mathrm{FD} > |\mathrm{ND}|$). Furthermore, because $\Delta$ is simply the logarithm of the optimum fitness (equation S1.26), this optimum fitness lies in the center, logarithmically speaking, of the bounds in equation S1.31.

#### S2 Conditions for other forms of overyielding

Overyielding occurs when coexisting species produce more biomass than some expected value, $Y_E$. In this Appendix, we generalize our framework to other forms of overyielding, which corresponds to results shown in Box 2 and Figure 3.

**Conditions for relative yield total** A less stringent definition of overyielding is provided by a quantity known as the relative yield total. The relative yield (Loreau 2010) is defined as the observed biomass of a species in a community divided by its biomass when growing alone. Accordingly, the relative yield total (RYT) is simply the sum of the relative yields of all species, and RYT $> 1$ corresponds to overyielding (relative to average intrinsic yield). For the case considered here, RYT can be calculated using equation S1.18 as

$$\text{RYT} = \sum_i \frac{\hat{N}_i}{K_i} = \frac{F_1 + F_2}{1 - \rho^2} \tag{S2.1}$$

Furthermore, since the denominator $1 - \rho^2$ is equal to $F_1 + F_2 - F_1 F_2$, it is always less than the numerator when species coexist ($F_1, F_2 > 0$), ensuring that RYT $> 1$. Thus, as noted previously, this form of overyielding is a necessary result of coexistence in the Lotka–Volterra and related models. This result also allows us to straightforwardly show that transgressive underyielding ($\Sigma \hat{N} < K_2$) is impossible in the system. Since species 2 is the lower yielding species ($K_2 < K_1$), we can also write

$$\begin{aligned} \frac{F_1 K_1 + F_2 K_2}{1 - \rho^2} &> \frac{F_1 K_2 + F_2 K_2}{1 - \rho^2} \\ \Sigma \hat{N} &> \text{RYT} \cdot K_2, \end{aligned} \tag{S2.2}$$

where the right-hand side is greater than $K_2$ because RYT $> 1$. Thus we find that $\Sigma \hat{N} > K_2$; that is, coexistence always causes the community to produce more biomass than its lowest yielding species would produce alone.

**Conditions for overyielding relative to average yield** More generally, overyielding $\Delta Y$ can be defined as the difference between the observed yield $Y_O = \Sigma_i N_i$ and some expected yield $Y_E$ (Loreau and Hector 2001). In experimental contexts, the expected yield is calculated by using initial relative abundances (e.g. during seeding or planting) to take a weighted average of species' yields in isolation ($K_i$). In the theoretical context here, we can simply weight species equally, defining expected yield as the mean yield in isolation

$\overline{K}$=$(K_1 + K_2)/2$ (note that other choices would give results intermediate between those shown above for transgressive overyielding and underyielding). Accordingly, we can use the expression for total abundance to derive the conditions under which $\Sigma\hat{N} > \overline{K}$; proceeding as above, we find that

$$\begin{aligned} F_1K_1 + F_2K_2 &> \frac{1}{2}(K_1+K_2)(F_1+F_2-F_1F_2) \\ \frac{1}{2}(K_1-K_2)(F_1-F_2) &> -\frac{1}{2}(K_1+K_2)F_1F_2 \end{aligned} \tag{S2.3}$$

Dividing through by the quantity $\frac{1}{2}(K_1-K_2)$ (positive since 1 is the higher yielding species), we find

$$F_1 - F_2 > -\frac{K_1+K_2}{K_1-K_2}F_1F_2; \tag{S2.4}$$

that is, overyielding (relative to average yield) is possible when the fitness of species 1 is sufficiently high relative to that of its competitor. Indeed, since the right-hand side is negative, $F_1 > F_2$ (or equivalently $f_1/f_2 > 1$) is a sufficient (but not necessary) condition for overyielding. Using $F_i = 1 - \rho\frac{f_j}{f_i}$, we can also rewrite the condition for overyielding relative to $\overline{K}$ (equation S2.4) in terms of niche overlap and fitness ratio as follows:

$$\begin{aligned} 2\overline{K} - \rho\left(\frac{f_2}{f_1}K_1 + \frac{f_1}{f_2}K_2\right) &> \overline{K}\left(1-\rho^2\right) \\ \rho\left(K_1\frac{f_2}{f_1} + K_2\frac{f_1}{f_2}\right) &< \frac{1}{2}(K_1+K_2)\left(1+\rho^2\right) \\ \left(\frac{f_1}{f_2}\right)^2 - \frac{1}{2}\left(1+\frac{K_1}{K_2}\right)\left(\frac{1}{\rho}+\rho\right)\frac{f_1}{f_2} + \frac{K_1}{K_2} &< 0, \end{aligned} \tag{S2.5}$$

where in the third line we have divided through by $\rho K_2 f_2/f_1$. Note that substituting $f_1/f_2 = 1$ satisfies this inequality; thus, species 1 having greater fitness is a sufficient (but not necessary) condition for overyielding. Solving the inequality in terms of $f_1/f_2$ gives the condition

$$B - \sqrt{B^2 - \frac{K_1}{K_2}} < \frac{f_1}{f_2} < B + \sqrt{B^2 - \frac{K_1}{K_2}}, \tag{S2.6}$$

where we write the shared expression in the lower and upper bounds as

$$B = \frac{1}{4}\left(1+\frac{K_1}{K_2}\right)\left(\frac{1}{\rho}+\rho\right). \tag{S2.7}$$

We visualize the lower and upper bounds of this condition in S2.1. As shown graphically, inequality S2.6 does not account for the prerequisite that the species coexist. Since transgressive overyielding also implies overyielding relative to average yield, the upper fitness bound in equation S1.23 must be less than or equal to the one considered here: $\rho^{-1} \leq B + \sqrt{B^2 - K_1/K_2}$. As $\rho^{-1}$ is also the maximum fitness ratio at which coexistence is possible, the complete condition for overyielding can be written

$$B - \sqrt{B^2 - \frac{K_1}{K_2}} < \frac{f_1}{f_2} < \rho^{-1}, \tag{S2.8}$$

with $B$ as above (equation S2.7). The lower bound $B - \sqrt{B^2 - \frac{K_1}{K_2}}$ delineates the lowest fitness ratio at which overyielding is possible; for complete niche overlap ($\rho = 1$), this is $f_1/f_2 = 1$, and this minimum fitness ratio decreases as the species experience increasing niche differentiation (decreasing $\rho$).

This boundary, visualized in Figure 3d in the main text and in more detail here in Supplemental Figure S2.1, is a hyperbola-like curve lying between the coexistence and transgressive overyielding conditions. To highlight the close relationship between it and the transgressive overyielding boundary explored above, we also consider the case when niche differentiation is high (i.e. $\rho \to 0$) and the term $B$ becomes large. Rewriting the boundary as $B\left(1 - \sqrt{1 - (K_1/K_2)\, B^{-2}}\right)$, we note that the term $B^{-2}$ becomes small as $\rho \to 0$, so we can apply the approximation $\sqrt{1-x} \approx 1 - x/2$ (a first-order Taylor approximation). Similarly, we can approximate $B \approx \frac{1}{4}\left(1 + \frac{K_1}{K_2}\right)\rho^{-1}$. Putting these together, we obtain the approximation

$$B - \sqrt{B^2 - \frac{K_1}{K_2}} \approx \frac{1}{2} \cdot \frac{K_1}{K_2} B^{-1} \approx \frac{2K_1}{K_2 + K_1}\rho = \frac{K_1}{\overline{K}}\rho \tag{S2.9}$$

which we note is strictly greater than the lower boundary in equation S2.8, indicating that this approximate boundary lies above the curved true boundary. Thus, we can write two sufficient (but not necessary) conditions,

$$\frac{f_1}{f_2} > \frac{K_1}{\overline{K}}\rho \text{ and} \tag{S2.10}$$

$$\frac{f_1}{f_2} > 1, \tag{S2.11}$$

which each ensure overyielding relative to average yield (provided that niche difference is sufficient for the species to coexist). Note that equation S2.10 has a similar form to the transgressive overyielding condition (equation S1.23) and thus can also be interpreted as requiring niche and fitness difference in excess of the conditions for coexistence, as we discussed previously (equation S1.31), with the degree of excess now $\Delta = \frac{1}{2}\log K_1/\overline{K}$ (with $\overline{K}$ instead of $K_2$ in the denominator). We illustrate the approximate conditions and their relationship to the full condition (equation S2.8) in Supplemental Figure S2.1.

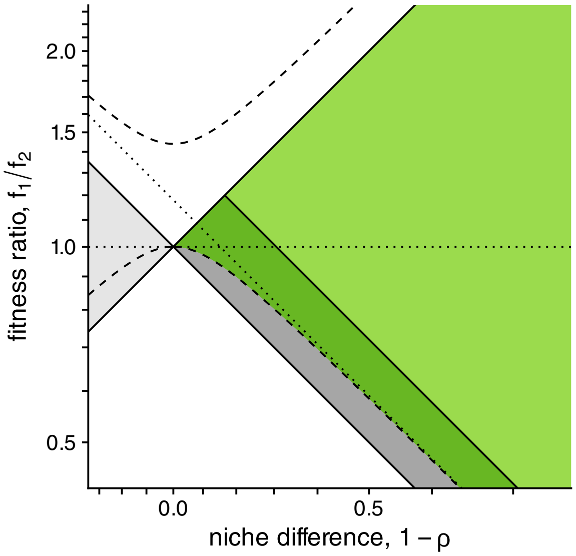


**Figure S2.1: Boundary and approximations for overyielding relative to average yield.** As in Figure 3d in the main text, we visualize the conditions for overyielding relative to $\overline{K}$ (dark + light green; corresponding to equation S2.8) alongside those for transgressive overyielding only (light green), coexistence without overyielding (dark gray), or alternative stable states (gray). We illustrate the mathematical form of the solution to the quadratic inequality by plotting the hyperbola-like pair of boundaries from equation S2.6 (dashed lines), where only the lower curve lies within the coexistence region. We also illustrate the approximations for this boundary from equation S2.10 (diagonal dotted line), which is parallel to the transgressive overyielding boundary, and equation S2.11 (horizontal dotted line), which simply requires that species 1 is relatively fitter.

**Sufficient condition for overyielding relative to any expected yield** Similarly, for any expected yield $Y_E$ between $K_1$ and $K_2$, we can compute the conditions for overyielding $F_1K_1 + F_2K_2 > \left(1-\rho^2\right) Y_E$, which we can rewrite in terms of $\rho$ and $f_1/f_2$ as

$$\left(\frac{f_1}{f_2}\right)^2 - \left(\frac{K_1+K_2-Y_E}{K_2}\rho^{-1} + \frac{Y_E}{K_2}\rho\right)\frac{f_1}{f_2} + \frac{K_1}{K_2} > 0, \tag{S2.12}$$

which then gives the minimum fitness condition,

$$\frac{f_1}{f_2} > B - \sqrt{B^2 - \frac{K_1}{K_2}}, \text{ where } B = \frac{1}{2}\left(\frac{K_1 + K_2 - Y_E}{K_2}\rho^{-1} + \frac{Y_E}{K_2}\rho\right). \tag{S2.13}$$

Applying the same approximations as previously, we find that for $\rho \to 0$, the minimum fitness for overyielding is approximately $\left(K_1 / \left(K_1 + K_2 - Y_E\right)\right)\rho$; substituting this expression for $f_1/f_2$ in equation S2.8 also shows that it defines a line above the overyielding boundary. Thus, a sufficient but not necessary condition for overyielding relative to any expected yield $Y_E$ is

$$\frac{f_1}{f_2} > \frac{K_1}{K_1 + K_2 - Y_E}\rho, \tag{S2.14}$$

which generalizes the previous results for transgressive underyielding ($f_1/f_2 > \rho$ for $Y_E = K_2$, thus the community must outperform its worst species, as shown above in this Appendix), overyielding relative to average yield ($f_1/f_2 > \left(K_1/\overline{K}\right)\rho$ for $Y_E = \overline{K}$, matching equation S2.8), and transgressive overyielding ($f_1/f_2 > \left(K_1/K_2\right)\rho$ for $Y_E = K_1$, matching equation 7 from the main text). This approximation fully generalizes our statement that overyielding requires niche and fitness differences in excess of those required for coexistence: generalizing equation S1.31, we can write that overyielding relative to $Y_E$ approximately requires that

$$\text{ND} - \Delta_{Y_E} > \left|\text{FD} - \Delta_{Y_E}\right|, \tag{S2.15}$$

where $\text{ND} = -\log\rho$ and $\text{FD} = \log f_1/f_2$ as before, and the degree of excess niche and fitness differences is given by

$$\Delta_{Y_E} = \frac{1}{2}\log\frac{K_1}{K_1 + K_2 - Y_E}, \tag{S2.16}$$

which is 0 when $Y_E = K_2$ and increases as the overyielding requirement $Y_E$ becomes increasingly stringent.

### S3 Applying functional coexistence theory to nonlinear competition

We consider a nonlinear extension of the Lotka–Volterra model.

$$\frac{dN_i}{dt} = r_i N_i \left[ 1 - \sum_j \alpha_{ij} N_j + \sum_j \gamma_{ij} \left( \alpha_{jj} N_j - \alpha_{jj}^2 N_j^2 \right) \right], \tag{S3.1}$$

where $\gamma_{ii}$ and $\gamma_{ij}$ add nonlinear responses to intra- and interspecific competition, and the model is the same as the Lotka–Volterra model in Appendix S1 when these nonlinearities are set to zero. We note that equivalent quadratic extensions of the per-capita growth rate have often been applied to study nonlinear competitive responses (Ayala et al. 1973; Gibbs et al. 2022); the parameterization here has been chosen so $\alpha_{ii}, \alpha_{ij}$ still reflect the carrying capacity and invasion properties of the system: as in the Lotka–Volterra model, $K_i \equiv \alpha_{ii}^{-1}$ is the carrying capacity of species $i$; since $\alpha_{ii} N_i = N_i / K_i$ and $\alpha_{ii} N_i - \alpha_{ii}^2 N_i^2 = (1 - N_i/K_i) \cdot N_i/K_i$, which is zero when species $i$ is absent or at carrying capacity, we can also see that $\gamma_{ii}$ and $\gamma_{ij}$ do not change the carrying capacity or scaled pairwise invasion growth $F_i = 1 - \alpha_{ij}/\alpha_{jj}$ of the model, respectively.

**Nondimensionalized model** To avoid working with so many coefficients, we can nondimensionalize this equation by defining a scaled biomass $n_j = \alpha_{jj} N_j$ which has been divided by carrying capacity (i.e., the same as its relative yield). Then the system can be written succinctly as

$$\frac{dn_i}{dt} = r_i n_i \left[ 1 - n_i + \gamma_{ii} \left( n_i - n_i^2 \right) - \sum_{j \neq i} \omega_{ij} n_j + \sum_{j \neq i} \gamma_{ij} \left( n_j - n_j^2 \right) \right] \tag{S3.2}$$

where $\omega_{ij} = \alpha_{ij}/\alpha_{jj}$. Now we solve for the coexistence outcome of a two-species system. As shown above, carrying capacity is $K_i \equiv \alpha_{ii}^{-1}$ and $F_i = 1 - \omega_{ij}$, matching the classic modern coexistence theory results. However, the coexistence equilibrium now requires solving a system of quadratic equations for $n_1, n_2$,

$$\begin{aligned} 1 &= n_1 + \omega_{12} n_2 - \gamma_{11} \left( n_1 - n_1^2 \right) - \gamma_{12} \left( n_2 - n_2^2 \right) \\ 1 &= \omega_{21} n_1 + n_2 - \gamma_{21} \left( n_1 - n_1^2 \right) - \gamma_{22} \left( n_2 - n_2^2 \right), \end{aligned} \tag{S3.3}$$

which is possible analytically, but produces expressions that are difficult to interpret.

**Perturbation analysis** Considering the case where $\gamma_{ii}, \gamma_{ij}$ are relatively small, we can instead apply perturbation theory (Otto and Day 2011; Strogatz 2015) to obtain a more biologically meaningful solution. We begin by solving the linear case ($\gamma_{ii}, \gamma_{ij} = 0$) to obtain a starting approximation. In this case, the equation is simply

$$\begin{aligned} 1 &= \tilde{n}_1 + \omega_{12}\tilde{n}_2 \\ 1 &= \omega_{21}\tilde{n}_1 + \tilde{n}_2 \end{aligned} \tag{S3.4}$$

which has the solution

$$\tilde{n}_i = \frac{1 - \omega_{ij}}{1 - \omega_{ij}\omega_{ji}} = \frac{F_i}{1 - \rho^2}, \tag{S3.5}$$

where $\rho^2 = \omega_{ij}\omega_{ji}$, corresponding to what we have previously shown for the Lotka–Volterra model, and we have denoted this solution $\tilde{n}_i$ with a tilde to indicate that it is a starting approximation. Perturbation starts with this linear solution and adds corrections for each of the small nonlinear effects.

Since we will only be interested in the linear terms of the perturbation series, we only need to consider the effect of a single $\gamma$ at a time. We illustrate this for the intraspecific nonlinear competition term $\gamma_{ii}$; including only this nonlinearity, the system of equations becomes

$$\begin{aligned} 1 &= n_i + \omega_{ij}n_j - \gamma_{ii}\left(n_i - n_i^2\right) \\ 1 &= \omega_{ji}n_i + n_j, \end{aligned} \tag{S3.6}$$

and we write the solution $n_i, n_j$ as a set of power series in $\gamma_{ii}$:

$$\begin{aligned} n_i &= \tilde{n}_i + \gamma_{ii}n_i^{(1)} + \gamma_{ii}^2 n_i^{(2)} + \cdots \\ n_j &= \tilde{n}_j + \gamma_{ii}n_j^{(1)} + \gamma_{ii}^2 n_j^{(2)} + \cdots, \end{aligned} \tag{S3.7}$$

that is, the approximate solution, plus a series of increasingly fine corrections in terms of powers of $\gamma_{ii}$. To actually obtain this correction, we solve for the first-order coefficients $n_i^{(1)}, n_j^{(1)}$. Substituting these expressions for $n_i, n_j$ into the system of equations in equation S3.6 and only considering terms up to first-order (i.e., dropping any terms that would contain $\gamma_{ii}^2, \gamma_{ii}^3, \ldots$), we obtain

$$0 = \gamma_{ii}\left[n_i^{(1)} + \omega_{ij}n_j^{(1)} - \left(\tilde{n}_i - \tilde{n}_i^2\right)\right] \tag{S3.8}$$

$$0 = \gamma_{ii}\left[\omega_{ji}n_i^{(1)} + n_j^{(1)}\right], \tag{S3.9}$$

where we have canceled out the terms corresponding to the original linear solution (equation S3.5) and factored out $\gamma_{ii}$. The key step of perturbation theory is noting that all terms of corresponding power series should match—in this case, this means the terms in square brackets are required to equal zero (as would the coefficients of higher-order terms) if the power series in equation S3.7 is to satisfy the system of equations. Thus, we can solve: from equation S3.9, we have $n_j^{(1)} = -\omega_{ji} n_i^{(1)}$, which we can substitute into equation S3.8 to obtain

$$n_i^{(1)} = \frac{\tilde{n}_i - \tilde{n}_i^2}{1-\rho^2} \quad \text{and} \quad n_j^{(1)} = -\omega_{ji} \frac{\tilde{n}_i - \tilde{n}_i^2}{1-\rho^2}, \tag{S3.10}$$

which we then substitute into the power series (equation S3.7) to obtain the first-order perturbation approximations,

$$\begin{aligned} n_i &\approx \frac{F_i}{1-\rho^2} + \frac{\gamma_{ii}\left(\tilde{n}_i - \tilde{n}_i^2\right)}{1-\rho^2} \quad \text{and} \\ n_j &\approx \frac{F_j}{1-\rho^2} - \frac{\omega_{ji}\gamma_{ii}\left(\tilde{n}_i - \tilde{n}_i^2\right)}{1-\rho^2}. \end{aligned} \tag{S3.11}$$

The first term in each expression is the linear solution (expected from classic modern coexistence theory) and the second term indicates that, if $\gamma_{ii}$ is positive, nonlinear effects due to $i$'s abundance benefit itself, ultimately increasing $n_i$ at the expense of $n_j$ (or, if $\gamma_{ii} < 0$, increasing $n_j$ at the expense of $n_i$). An analogous analysis for the interspecific nonlinearity $\gamma_{ij}$ gives corresponding expressions

$$\begin{aligned} n_i &\approx \frac{F_i}{1-\rho^2} + \frac{\gamma_{ij}\left(\tilde{n}_j - \tilde{n}_j^2\right)}{1-\rho^2} \\ n_j &\approx \frac{F_j}{1-\rho^2} - \frac{\omega_{ji}\gamma_{ij}\left(\tilde{n}_j - \tilde{n}_j^2\right)}{1-\rho^2}, \end{aligned} \tag{S3.12}$$

which now account for the effect of $j$'s nonlinearity on $i$.

**Full first-order perturbation solution** Accounting for the first-order perturbation terms of all four nonlinearities (following equations S3.11 and S3.12), we find:

$$\begin{aligned} n_i &= \frac{F_i}{1-\rho^2} + \frac{\gamma_{ii} - \omega_{ij}\gamma_{ji}}{1-\rho^2}\left(\tilde{n}_i - \tilde{n}_i^2\right) + \frac{\gamma_{ij} - \omega_{ij}\gamma_{jj}}{1-\rho^2}\left(\tilde{n}_j - \tilde{n}_j^2\right) \\ &= \frac{1}{1-\rho^2}\left[F_i + \psi_{ii}\left(\tilde{n}_i - \tilde{n}_i^2\right) + \psi_{ij}\left(\tilde{n}_j - \tilde{n}_j^2\right)\right] \end{aligned} \tag{S3.13}$$

where each coefficient $\psi_{ik} = \gamma_{ik} - \omega_{ij}\gamma_{jk}$ (for $j \neq i$) measures how much $i$ benefits due to nonlinearities resulting from $k$, and each of the $\tilde{n}_i - \tilde{n}_i^2$ terms approximate the degree of these nonlinearities.

**Expression for community biomass** Since abundance is $N_i = n_i K_i$, we can write the total abundance at equilibrium as

$$\hat{N}_1 + \hat{N}_2 = \frac{1}{1-\rho^2}\left[F_1K_1 + F_2K_2 + \delta_1\left(\tilde{n}_1 - \tilde{n}_1^2\right) + \delta_2\left(\tilde{n}_2 - \tilde{n}_2^2\right)\right], \tag{S3.14}$$

where the term

$$\begin{aligned}\delta_i &= \psi_{ii}K_i + \psi_{ji}K_j \\ &= \left(\gamma_{ii} - \omega_{ij}\gamma_{ji}\right)K_i + \left(\gamma_{ji} - \omega_{ji}\gamma_{ii}\right)K_j \\ &= \gamma_{ii}\left(K_i - \omega_{ji}K_j\right) + \gamma_{ji}\left(K_j - \omega_{ij}K_i\right),\end{aligned} \tag{S3.15}$$

measures how total biomass is affected by nonlinear responses to species $i$, and $\tilde{n}_i = F_i / \left(1-\rho^2\right)$ as given above. Thus, we see that both the classic coexistence theory prediction (given by $\left(F_1K_1 + F_2K_2\right) / \left(1-\rho^2\right)$, as explored in the main text), as well as the effects of nonlinearity (terms controlled by $\delta_i$) can be expressed in terms of the intrinsic yields $K_i$ and the scaled invasion growth rates $F_i$ (and thus the niche and fitness differences, as in the classical case explored throughout this manuscript). We can rewrite $F_i$ in terms of $\tilde{n}_i$ (using equation S3.5) to obtain that total biomass is

$$\begin{aligned}&\hat{N}_1 + \hat{N}_2 \\ =&\frac{1}{1-\rho^2}\left[\left(\left(\delta_1 + \left(1-\rho^2\right)K_1\right)\tilde{n}_1 - \delta_1\tilde{n}_1^2\right) + \left(\left(\delta_2 + \left(1-\rho^2\right)K_2\right)\tilde{n}_2 - \delta_2\tilde{n}_2^2\right)\right]\end{aligned} \tag{S3.16}$$

which, completing the square using $bx - ax^2 = -a\left(x - b/2a\right)^2 + a\left(b/2a\right)^2$ and simplifying, becomes

$$\hat{N}_1 + \hat{N}_2 = \frac{1}{\left(1-\rho^2\right)}\left[\delta_1\tilde{n}_1^{*2} + \delta_2\tilde{n}_2^{*2} - \delta_1\left(\tilde{n}_1 - \tilde{n}_1^*\right)^2 - \delta_2\left(\tilde{n}_2 - \tilde{n}_2^*\right)^2\right], \tag{S3.17}$$

where we define

$$\tilde{n}_i^* \equiv \frac{1}{2} + \frac{\left(1-\rho^2\right)K_i}{2\delta_i}, \tag{S3.18}$$

which we can see is the value optimizing (i.e. maximizing if $\delta_i > 0$ or minimizing if $\delta_i < 0$) yield. Furthermore, since $\tilde{n}_i = F_i / \left(1-\rho^2\right)$, we can write this in terms of $F_1, F_2$ as

$$\hat{N}_1 + \hat{N}_2 = \frac{1}{\left(1-\rho^2\right)^3}\left[\delta_1 F_1^{*2} + \delta_2 F_2^{*2} - \delta_1 \left(F_1 - F_1^*\right)^2 - \delta_2 \left(F_2 - F_2^*\right)^2\right], \tag{S3.19}$$

where

$$F_i^* \equiv \tilde{n}_i^* \left(1-\rho^2\right) = \frac{1}{2}\left(1-\rho^2\right) + \frac{K_i}{2\delta_i}\left(1-\rho^2\right)^2 \tag{S3.20}$$

is the value optimizing yield.

**General overyielding condition** Generally, we can show that overyielding relative to some arbitrary $Y_E$ requires

$$\frac{1}{\left(1-\rho^2\right)}\left[F_1 K_1 + F_2 K_2 + \delta_1\left(\tilde{n}_1 - \tilde{n}_1^2\right) + \delta_2\left(\tilde{n}_2 - \tilde{n}_2^2\right)\right] > Y_E, \tag{S3.21}$$

where as above, the terms with $\delta_i$ create departure from the classical case. We can also rewrite $F_i$ in terms of $\tilde{n}_i$ (using equation S3.5)

$$\delta_1 \tilde{n}_1^{*2} + \delta_2 \tilde{n}_2^{*2} - \left(1-\rho^2\right) Y_E > \delta_1 \left(\tilde{n}_1 - \tilde{n}_1^*\right)^2 + \delta_2 \left(\tilde{n}_2 - \tilde{n}_2^*\right)^2, \tag{S3.22}$$

whose boundary is a conic section in terms of $\tilde{n}_2, \tilde{n}_2$ (an ellipse if $\delta_1, \delta_2$ share a sign; otherwise a hyperbola) centered on $\tilde{n}_1^*, \tilde{n}_2^*$. Note that this interpretation is most useful when $\delta_i$ is not too small, since the center of the conic section diverges as the boundary approaches the classical case ($\delta_i \to 0$).

**Condition for transgressive overyielding** Using $Y_E = K_2$ in equation S3.21 and rearranging shows that transgressive overyielding occurs when species 1 has sufficient fitness:

$$F_1 > 1 - \frac{K_2}{K_1} - \left[\delta_1\left(\tilde{n}_1 - \tilde{n}_1^2\right) + \delta_2\left(\tilde{n}_2 - \tilde{n}_2^2\right)\right] F_2^{-1} K_1^{-1}, \tag{S3.23}$$

which is sufficient to predict transgressive overyielding from coexistence theory metrics and the degree of nonlinearity in the system. As we have discussed above, the boundary approaches the classical case derived in Appendix S1 when the system is nearly linear ($\delta_1, \delta_2 \approx 0$); furthermore, nonlinearities will become less important when species' abundances are closer to competitive exclusion ($\tilde{n}_1 - \tilde{n}_1^2, \tilde{n}_2 - \tilde{n}_2^2 \approx 0$).

#### S4 Calculating the additive partition

The additive partition of biodiversity effects separates the degree of overyielding into two components, selection and complementarity. Loreau and Hector (2001) measure these components using the concept of relative yield, $\mathrm{RY}_i = N_i^*/K_i$, i.e., a species' abundance relative to its abundance when growing alone. Applying this notation, the additive partition can be written in terms of means ($\overline{\cdots}$) and covariance ($\mathrm{cov}\,(\ldots)$) as

$$\Delta Y = \underbrace{n \cdot \overline{\Delta \mathrm{RY}} \cdot \overline{K}}_{\text{complementarity}} + \underbrace{n \cdot \mathrm{cov}\,(\Delta \mathrm{RY}, K)}_{\text{selection}}, \tag{S4.1}$$

where $\Delta Y = \sum_i N_i^* - \sum_i \mathrm{RY}_{E,i} K_i$ is overyielding (the difference between observed and expected yield), $n$ (originally denoted $N$ in Loreau and Hector 2001) is the number of species, $\Delta \mathrm{RY}_i = N_i^*/K_i - \mathrm{RY}_{E,i}$ is the difference between observed and expected relative yield, and $K_i$ ($M_i$ in the original notation) is species $i$'s yield when growing alone. As discussed above, we follow previous treatments (Carroll et al. 2011) in calculating overyielding relative to the average yield when growing alone, corresponding to the case where expected relative yield is simply $1/n$ for all species.

**Calculating the additive partition** For the model considered here, relative yield can be calculated from equation S1.18 as simply $F_i/\left(1-\rho^2\right)$. Accordingly, we can calculate the partition in equation S4.1 using $\Delta \mathrm{RY_i} = F_i/\left(1-\rho^2\right) - 1/2$, giving

$$\Delta Y = \underbrace{\left(\frac{F_1 + F_2}{1-\rho^2} - 1\right)\overline{K}}_{\text{complementarity}} + \underbrace{\frac{2\,\mathrm{cov}\,(F, K)}{1-\rho^2}}_{\text{selection}}, \tag{S4.2}$$

where subtraction of the constant $1/2$ does not affect the covariance term. Applying the identity $1-\rho^2 = F_1 + F_2 - F_1F_2$, we can show that the bracketed term in the selection effect term is simply $F_1F_2/\left(1-\rho^2\right)$; meanwhile, applying the definition of covariance and denoting $\Delta F = F_1 - F_2$, $\Delta K = K_1 - K_2$, we find that

$$\begin{aligned}
\mathrm{cov}\,(F, K) &= \frac{1}{2}F_1K_1 + \frac{1}{2}F_2K_2 - \overline{F}\cdot\overline{K} \\
&= \frac{1}{2}\left(\overline{F} + \frac{1}{2}\Delta F\right)\left(\overline{K} + \frac{1}{2}\Delta K\right) + \frac{1}{2}\left(\overline{F} - \frac{1}{2}\Delta F\right)\left(\overline{K} - \frac{1}{2}\Delta K\right) - \overline{F}\cdot\overline{K} \\
&= \frac{1}{4}\Delta F \cdot \Delta K.
\end{aligned} \tag{S4.3}$$

Accordingly, we can write out the additive partition as

$$\Delta Y = \underbrace{\frac{F_1 F_2}{1-\rho^2} \cdot \frac{K_1 + K_2}{2}}_{\text{complementarity, CE}} + \underbrace{\frac{F_1 - F_2}{1-\rho^2} \cdot \frac{K_1 - K_2}{2}}_{\text{selection, SE}}, \tag{S4.4}$$

where we can see that complementarity (CE) is positive for coexisting species and selection (SE) is positive when the higher yielding species 1 also has greater fitness. Using the identity $F_i = 1 - \rho \left( f_j / f_i \right)$, this could also be written purely in terms of niche overlap and fitness ratio,

$$\Delta Y = \frac{1 + \rho^2 - \rho \left( \frac{f_1}{f_2} + \frac{f_2}{f_1} \right)}{1 - \rho^2} \cdot \overline{K} + \frac{\rho \left( \frac{f_1}{f_2} - \frac{f_2}{f_1} \right)}{1 - \rho^2} \cdot \frac{K_1 - K_2}{2}. \tag{S4.5}$$

We can now relate the additive partition to the functional coexistence theory components by considering the change in complementarity or selection when niche, fitness, and function are changed (i.e., by taking the partial derivative). We summarize these results in Table S4.1 and provide full proofs of these results below: niche differentiation is consistently positively related to complementarity, fitness difference is consistently related to selection, and yield imbalance had a direct effect only on the magnitude of selection.

**Table S4.1: Effects of niche, fitness, and function on the additive partition components.** We show the effects of varying each functional coexistence process (left column) on the complementarity (middle) and selection effects (right). Note that increasing niche differentiation actually involves decreasing $\rho$.

| Functional coexistence process | Effect on complementarity | Effect on selection |
|---|---|---|
| increasing niche difference, $\rho \to 0$ | $+$ | SE $\to 0$ ($-$ if SE $> 0$; $+$ if SE $< 0$) |
| increasing fitness of higher yielding species, $f_1/f_2 \to \infty$ | inconsistent ($+$ only if $f_1/f_2 \to 1$) | $+$ |
| equalizing yields in isolation, $K_1 - K_2 \to 0$ | no direct effect | SE $\to 0$ ($-$ if SE $> 0$; $+$ if SE $< 0$) |

**Niche differentiation increases complementarity** We first consider the effect of increasing niche differentiation (decreasing $\rho$) on the complementarity effect CE. To do so, we consider the sign of $-\partial\text{CE}/\partial\rho$, where the negative sign reflects the inverse relationship between $\rho$ and niche differentiation. Since $\overline{K}$ is positive and independent of $\rho$, we only need to consider the first term in the complementarity expression from equation S4.4,

$$\begin{aligned}\frac{\text{CE}}{\overline{K}} = \frac{F_1F_2}{(1-\rho^2)} &= \frac{F_1F_2}{F_1+F_2-F_1F_2} \\ &= (F_1^{-1}+F_2^{-1}-1)^{-1},\end{aligned} \tag{S4.6}$$

where we have used the identity $1-\rho^2 = F_1+F_2-F_1F_2$ and divided the numerator and denominator through by $F_1F_2$. While we could directly rewrite this equation in terms of $\rho$ and $f_1/f_2$ and take the derivative, we can simplify the derivation by noting that since $0 < F_i < 1$ as a consequence of coexistence in the competitive model, the reciprocal of the expression in equation S4.6 has a strictly decreasing relationship to the complementarity effect. Thus, the sign of $-\partial\text{CE}/\partial\rho$ is the same as the sign of the derivative of this reciprocal with respect to $\rho$,

$$\frac{\partial}{\partial\rho}\left[F_1^{-1}+F_2^{-1}-1\right] = \frac{f_2}{f_1}\cdot F_1^{-2} + \frac{f_1}{f_2}\cdot F_2^{-2}, \tag{S4.7}$$

where we have applied the chain rule and used the fact that $\partial F_i/\partial\rho = -f_j/f_i$. We note that equation S4.7 is always positive since $F_i$ and $f_i/f_j$ are always positive for coexisting species. Thus, we conclude that $-\partial\text{CE}/\partial\rho > 0$. Stated conceptually, *increasing niche differentiation (i.e., decreasing ρ) always increases the complementarity effect.*

**Fitness has a variable effect on complementarity** Similarly, we consider the effect of fitness on complementarity by finding the sign of $\partial\text{CE}/\partial(f_1/f_2)$ i.e., the effect of increasing the fitness of species 1 (the higher yielding species) on the complementarity effect. Rewriting the expression in equation S4.6 in terms of $\rho$ and $f_1/f_2$ (as in equation S4.5), we find that

$$\begin{aligned}\frac{\partial}{\partial(f_1/f_2)}\left[\frac{1+\rho^2-\rho\left(\frac{f_1}{f_2}+\frac{f_2}{f_1}\right)}{1-\rho^2}\right] &= -\frac{\rho}{1-\rho^2}\cdot\frac{\partial}{\partial(f_1/f_2)}\left[\frac{f_1}{f_2}+\frac{f_2}{f_1}\right] \\ &= -\frac{\rho}{1-\rho^2}\left(1-\left(\frac{f_1}{f_2}\right)^{-2}\right),\end{aligned} \tag{S4.8}$$

which is positive for $f_1/f_2 < 1$ but negative if $f_1/f_2 > 1$. Thus, we conclude that $\partial\text{CE}/\partial(f_1/f_2) > 0$ only when $f_1/f_2 < 1$. Thus, *the direction of the effect of fitness on* *complementarity is inconsistent*, but bringing fitness difference closer to 1 always increases complementarity.

**Niche differentiation has a variable effect on selection** Turning to the selection component, we consider how increasing niche differentiation changes the selection effect. Similarly to above, since niche differentiation involves decreasing niche overlap $\rho$, we are interested in the sign of $-\partial \mathrm{SE}/\partial\rho$. Note that the term $(K_1 - K_2)/2$ in equations S4.4 and S4.5 is always positive because we defined species 1 as having higher intrinsic yield. Thus, we only need to consider the term

$$\mathrm{SE} \cdot \frac{2}{K_1 - K_2} = \frac{F_1 - F_2}{1-\rho^2} = \frac{\rho\left(\frac{f_1}{f_2} - \frac{f_2}{f_1}\right)}{1-\rho^2}, \tag{S4.9}$$

Then we can find the sign of the effect of interest by considering

$$\begin{aligned} -\frac{\partial}{\partial\rho}\left[\frac{\rho\left(\frac{f_1}{f_2} - \frac{f_2}{f_1}\right)}{1-\rho^2}\right] &= -\left(\frac{f_1}{f_2} - \frac{f_2}{f_1}\right) \cdot \frac{\partial}{\partial\rho}\left[\frac{\rho}{1-\rho^2}\right] \\ &= -\left(\frac{f_1}{f_2} - \frac{f_2}{f_1}\right) \cdot \frac{1+\rho^2}{\left(1-\rho^2\right)^2}, \end{aligned} \tag{S4.10}$$

which always has the opposite sign as the selection effect: the term $f_1/f_2 - f_2/f_1$ also determines the sign of the selection effect (equation S4.9), while the last term is always positive. Thus, $-\partial \mathrm{SE}/\partial\rho > 0$ only when $\mathrm{SE} < 0$. In other words, *the direction of the effect of niche differentiation on selection is inconsistent*, but increasing niche differentiation (i.e., decreasing $\rho$) always decreases the magnitude of the selection effect ($\mathrm{SE} \to 0$)

**Fitness is consistently related to selection** To understand how fitness affects the selection effect, we consider the sign of the partial derivative $\partial \mathrm{SE}/\partial\left(f_1/f_2\right)$. As above, we can simply consider the sign of the derivative with respect to the scaled expression in equation S4.9, finding that

$$\frac{\partial}{\partial\left(f_1/f_2\right)}\left[\frac{\rho\left(\frac{f_1}{f_2} - \frac{f_2}{f_1}\right)}{1-\rho^2}\right] = \frac{\rho}{1-\rho^2} \cdot \frac{\partial}{\partial\left(f_1/f_2\right)}\left[\frac{f_1}{f_2} - \frac{f_2}{f_1}\right] \tag{S4.11}$$

$$= \frac{\rho}{1-\rho^2}\left(1 + \left(\frac{f_1}{f_2}\right)^{-2}\right), \tag{S4.12}$$

which is always positive because $f_1/f_2$ is positive. Thus, $\partial\text{SE}/\partial(f_1/f_2) > 0$ always holds as a consequence of coexistence; that is, *increasing the fitness ratio in favor of the higher yielding species always makes the selection effect more positive.*

**The effect of yield** Finally, we can consider the effect of changing the imbalance between the two species' biomass yields when growing alone ($K_1$ vs. $K_2$). According to equation S4.4, the complementarity effect depends only on $\overline{K}$ (mean biomass when growing alone); thus, *difference in yield has no direct effect on complementarity*, though concomitant changes in average yield can affect it (as seen in Figure 3c in the main text). Only the selection effect reflects inequality between the two species' yields; in particular, the effect of increasing $K_1 - K_2$ has the same sign as $f_1/f_2 - f_2/f_1$. Thus, the *direction of the effect of yield difference on selection is inconsistent*, but increasing the yield of the species with higher fitness always positively affects selection.

**Summary: overyielding versus the additive partition** We now summarize our theoretical results for coexistence and transgressive overyielding (Appendix S1), other forms of overyielding (Appendix S2), and the additive partition (the present Appendix). In Figure S4.1, we illustrate all possible combinations of overyielding and additive partition outcomes for two-species competition under the class of models considered here. Coexistence theory predicts whether niche and fitness differences are sufficient for coexistence (I–III) or whether one species excludes its competitor (IV–IV). Functional coexistence theory then predicts whether the coexisting community exhibits transgressive overyielding (I), overyielding relative to average intrinsic yield (II), or neither (III). Finally, according to S4.4, we can see that complementarity is always positive for coexisting species, but that the sign of the selection effect depends on fitness: positive if the higher yielding species is fitter ($f_1/f_2 > 1$; I–IIa) or negative otherwise (I–IIb). We summarize these outcomes in Table S4.2. While complementarity is positive for all forms of overyielding (a consequence of coexistence), both positive or negative selection effects are possible for each of the two forms of overyielding considered here.

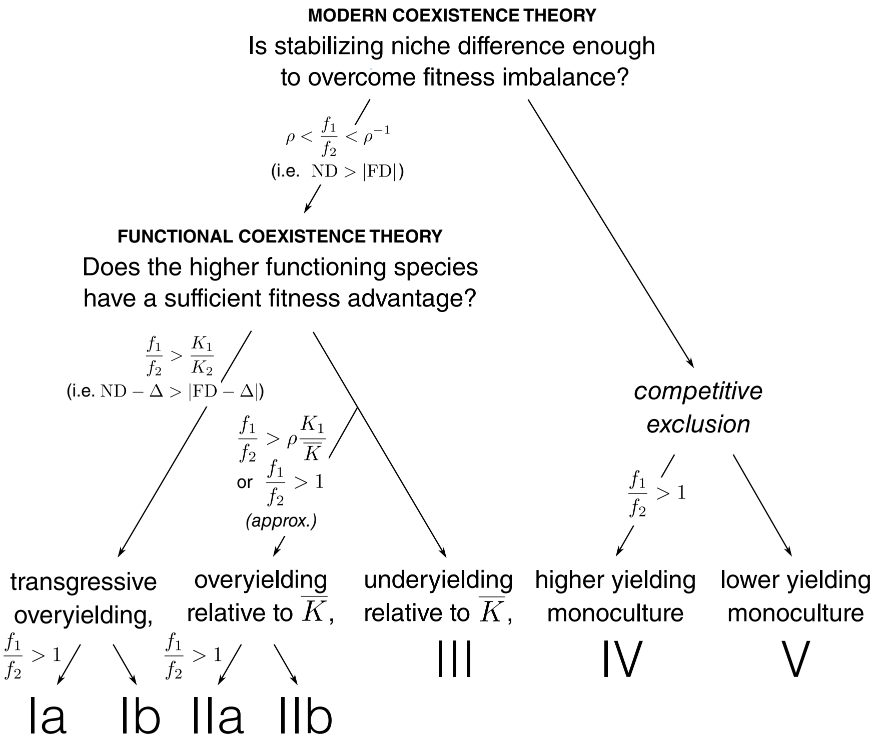


**Figure S4.1: Predicting outcomes using niche, fitness, and function.** We illustrate the results from the present Appendix together with those from Appendices S1 and S2. As described in greater detail in the text, the outcomes of modern coexistence theory (top: I–III vs. IV–V) and functional coexistence theory (middle left: I vs. II vs. III) as well as the signs of additive partition components (bottom left: I–IIa vs. I–IIb) can be predicted using niche, fitness, and/or function (conditions given on relevant arrows).

**Table S4.2: Predicting the signs of overyielding and additive partition components.** Roman numerals and letters correspond to the categorization in Figure S4.1; for each, we give the signs of differences in yield (above) and the additive partition components (below). Since I–IIa differ from I–IIb only in the sign of the selection effect (positive vs. negative), we show each pair together and indicate the sign as $+/-$.

| | | Ia/b | IIa/b | III | IV | V |
|---|---|---|---|---|---|---|
| overyielding relative to... | lowest yield | $+$ | $+$ | $+$ | $+$ | 0 |
| | average yield | $+$ | $+$ | $-$ | $+$ | $-$ |
| | best yield | $+$ | $-$ | $-$ | 0 | $-$ |
| additive partition... | complementarity | $+$ | $+$ | $+$ | 0 | 0 |
| | selection | $+/-$ | $+/-$ | $-$ | $+$ | $-$ |

### S5 Interpreting complementarity as a niche difference measure

In this Appendix, we discuss the calculation of niche and fitness difference in order to show that complementarity can be interpreted as a niche difference measure. To do so, we focus on the Lotka–Volterra model and review two previously used sets of metrics: the first consists of the widely-used niche overlap $\rho$ and fitness ratio $f_1/f_2$ and is termed "geometric" by Spaak, Ke, et al. (2023); the second corresponds to more general versions of modern coexistence theory and is termed "arithmetic". We show that complementarity encodes many of the same intuitions as these definitions, and indeed that it is closely related to the arithmetic definition of niche difference. Accordingly, we illustrate how it can also be used with a corresponding fitness metric to define conditions for coexistence.

**Geometric definition of niche and fitness** Throughout this exposition, we have used the widely-adopted niche overlap $\rho$ and fitness ratio $f_1/f_2$ metrics. Originally from Chesson and Kuang (2008), this method and its extensions (Carroll et al. 2011; Spaak and De Laender 2020) calculate niche and fitness as a geometric mean of species' scaled reduction in invasion growth rates: for the Lotka–Volterra model, and using our notation from Appendices S1–S4 of $F_i = \mathrm{IGR}_i/r_i$ (where $r_i$ is species $i$'s intrinsic rate of increase and $\mathrm{IGR}_i$ is its invasion growth rate), we can write that

$$\rho = \sqrt{(1 - F_1)(1 - F_2)} \tag{S5.1}$$

$$\frac{f_1}{f_2} = \sqrt{\frac{1 - F_2}{1 - F_1}}, \tag{S5.2}$$

as stated in Box 1; this can be verified by substituting $F_i = 1 - \alpha_{ij}/\alpha jj$ to obtain the original definitions of these quantities. Since equation S5.1 is a geometric mean of the quantities $1 - F_i$, Spaak, Ke, et al. (2023) termed it the "geometric definition" of niche and fitness differences. These ratios can be converted into differences by taking logarithms to define

$$\begin{aligned} \mathrm{ND_{geo}} &= -\log \rho \\ &= \frac{1}{2}\left[-\log(1 - F_1) - \log(1 - F_2)\right] \end{aligned} \tag{S5.3}$$

$$\begin{aligned} \mathrm{FD}_{\mathrm{geo}} &= \log \frac{f_1}{f_2} \\ &= \frac{1}{2}\left[-\log\left(1-F_1\right)+\log\left(1-F_2\right)\right], \end{aligned} \tag{S5.4}$$

such that the coexistence condition is $\mathrm{ND} > |\mathrm{FD}|$, as depicted throughout the main text (e.g. Figure 1). Note that these are respectively the average and the difference from the average of the species-specific quantities $-\log(1-F_i)$; in the competitive Lotka–Volterra model, this quantity is positive for a species if and only if its invasion growth rate is positive ($F_i > 0$).

**Arithmetic definition of niche and fitness** As several reviews of the modern coexistence theory framework have noted (Godwin et al. 2020; Song et al. 2019; Spaak, Ke, et al. 2023), there are numerous alternative methods for quantifying niche and fitness difference. One family of methods uses arithmetic, not geometric, means to define niche difference and was thus termed the "arithmetic definition" by Spaak, Ke, et al. (2023). For our purposes, we provide a set of arithmetic definitions as

$$\begin{aligned} \mathrm{ND}_{\mathrm{ari}} &= \overline{F} \\ &= \frac{1}{2}\left(F_1+F_2\right) \end{aligned} \tag{S5.5}$$

$$\begin{aligned} \mathrm{FD}_{\mathrm{ari}} &= F_1-\overline{F} \\ &= \frac{1}{2}\left(F_1-F_2\right), \end{aligned} \tag{S5.6}$$

which correspond exactly to the definitions proposed by Chesson (2003) and Barabás et al. (2018), notated there as $A$ and $f_i - A$, when the intrinsic growth rate $r_i$ is chosen as the scaling factor (Barabás et al. 2018; Johnson and Hastings 2022). As above, $\mathrm{ND} > |\mathrm{FD}|$ is a necessary and sufficient condition for coexistence. Spaak, Ke, et al. (2023) and Zhao et al. (2016) originally provided a slightly different formulation in which the invasion growth rates $\mathrm{IGR}_i$ are used without scaling, and the fitness definition does not directly predict coexistence.

**General properties of niche and fitness metrics** The arithmetic and geometric definitions considered here share certain properties due to the general principle that any set of niche and fitness metrics should predict the outcome of competition (Spaak, Ke, et al. 2023). One consequence of this requirement is that niche difference should be a necessary condition for coexistence: in other words, niche differentiation should be positive whenever species can coexist (i.e., ND $> 0$). Another consequence is that fitness difference should reflect competitive hierarchy in the system: stated more precisely, any process that shifts the system from competitive dominance by species $i$ to competitive dominance by species $j$ should involve a shift in sign of the fitness metric (e.g., from $\mathrm{FD}_i > 0$ to $\mathrm{FD}_i < 0$). We illustrate these commonalities using a series of simulations. In Figure S5.1a, we consider processes (lines 1–3) that purely affect either niche or fitness under the geometric definition and calculate the corresponding arithmetic niche and fitness (center plot); similarly, in Figure S5.1b we consider processes (lines 4–6) that purely affect the arithmetic metrics (center plot) and illustrate the corresponding geometrically defined ones (left plot). In both cases, coexistence (dark gray region) is associated with positive niche difference ($\mathrm{ND}_{\mathrm{geo}} = -\log\rho > 0$ or $\mathrm{ND}_{\mathrm{ari}} = (F_1 + F_2)/2 > 0$). Indeed, increasing either geometric or arithmetic niche difference (3 or 6, respectively) also increases the other definition of niche difference. Similarly, in both the geometric (1–2) or arithmetic (4–5) cases, going from competitive exclusion by species 1 to coexistence then to competitive exclusion by species 2 involves going from positive to negative fitness difference (change of sign in $\mathrm{ND}_{\mathrm{geo}} = \log f_1/f_2$ or $\mathrm{ND}_{\mathrm{ari}} = (F_1 - F_2)/2$, respectively). Indeed, the geometric and arithmetic fitness metrics share signs: we note that fitness difference is positive (e.g., along 1 and 4) or negative (e.g., along 2 and 5) in the same cases regardless of the choice of metrics.

**Scaled complementarity in the Lotka–Volterra model** We will show that complementarity encodes similar information as the existing geometric and arithmetic fitness metrics. As a measure of complementarity, we divide the standard definition from Loreau and Hector (2001, i.e., $n \cdot \overline{\Delta\mathrm{RY}} \cdot \overline{K}$, as defined in S4.1) by the average intrinsic yield $\overline{K}$ to obtain a unitless scaled complementarity effect. For two coexisting species, we can derive this from equation S4.2 by substituting $1 - \rho^2 = F_1 + F_2 - F_1 F_2$ to obtain

$$\frac{\mathrm{CE}}{\overline{K}} = \mathrm{RYT} - 1 = \frac{F_1 + F_2}{F_1 + F_2 - F_1 F_2} - 1 = \frac{F_1 F_2}{F_1 + F_2 - F_1 F_2}, \tag{S5.7}$$

where RYT is the relative yield total, as defined in S2.1. Carroll et al. (2011) calculated the same scaled complementarity for the Lotka–Volterra model; expressing this in terms of (geometric) niche overlap $\rho$, they stressed the quantitative incompatibility between complementarity and this measure of niche differentiation. However, despite this difference, we note that like the geometric and arithmetic niche definitions, this metric is positive whenever species coexist (i.e., $F_1, F_2 > 0$ implies $\mathrm{RYT} - 1 > 0$).

**Complementarity-based niche and fitness metrics** Accordingly, we can extend complementarity into a set of niche and fitness metrics that capture the same information as the geometric and arithmetic ones. Defining a non-negative scaling coefficient,

$$C = 2\left|\frac{1}{F_1 + F_2 - F_1 \cdot F_2} - \frac{1}{F_1 + F_2}\right| = 2\left|\frac{F_1 \cdot F_2}{(F_1 + F_2)(F_1 + F_2 - F_1F_2)}\right|, \tag{S5.8}$$

we can define a complementarity-based niche difference as

$$\mathrm{ND}_{\mathrm{com}} \equiv C \cdot \frac{1}{2}(F_1 + F_2), \tag{S5.9}$$

i.e., the arithmetic niche difference $\mathrm{ND}_{\mathrm{ari}}$ scaled by $C$. Noting that the quantity inside the absolute value in the definition for $C$ (equation S5.9) is positive when species coexist ($F_1, F_2 > 0$), we can multiply through to find that $\mathrm{ND}_{\mathrm{com}} = \mathrm{RYT} - 1$ for coexisting species. The corresponding fitness difference can then be found by scaling $\mathrm{FD}_{\mathrm{ari}}$ by $C$ to obtain

$$\mathrm{FD}_{\mathrm{com}} = C \cdot \frac{1}{2}(F_1 - F_2), \tag{S5.10}$$

such that the coexistence condition is still $\mathrm{ND} > |\mathrm{FD}|$.

**Interpreting the complementarity-based metrics** In Figure S5.1, we visualize how the complementarity-based niche and fitness metrics $\mathrm{ND}_{\mathrm{com}}$ and $\mathrm{FD}_{\mathrm{com}}$ (rightmost plot in each panel) are related to the geometric (left) and arithmetic (center) definitions. The results are comparable regardless of whether we consider processes purely affecting the geometric components (panel a) or the arithmetic ones (panel b). Though the relationship between the metrics is somewhat complex, they give the same conclusions regarding coexistence: increasing geometric or arithmetic niche difference while fixing fitness difference (3 and 6, respectively; in blue) initially decreases complementarity-based niche difference, but eventually results in coexistence with increasingly strong complementarity. Similarly, decreasing the geometric or arithmetic fitness difference while fixing niche difference (1–2 and 4–5, respectively; in red) also decreases the complementarity-based fitness difference in favor of species 1 until coexistence is reached; eventually, species 1 becomes so competitive that it competitively excludes species 2. Indeed, positive or negative $\mathrm{FD}_{\mathrm{com}}$ (indicating species 1 is the superior competitor) also implies that the other fitness differences ($\mathrm{FD}_{\mathrm{geo}}$ or $\mathrm{FD}_{\mathrm{ari}}$) are also positive or negative, respectively. These close relationships emerge because $\mathrm{ND}_{\mathrm{com}}$ and $\mathrm{FD}_{\mathrm{com}}$ are simply the arithmetic metrics scaled by a non-negative constant. Thus, overall, we find that the complementarity metrics satisfy the same roles as the geometric and arithmetic ones: coexistence requires positive $ND_{com}$, and reversing the outcome of competitive exclusion requires a change in the sign of $FD_{com}$.

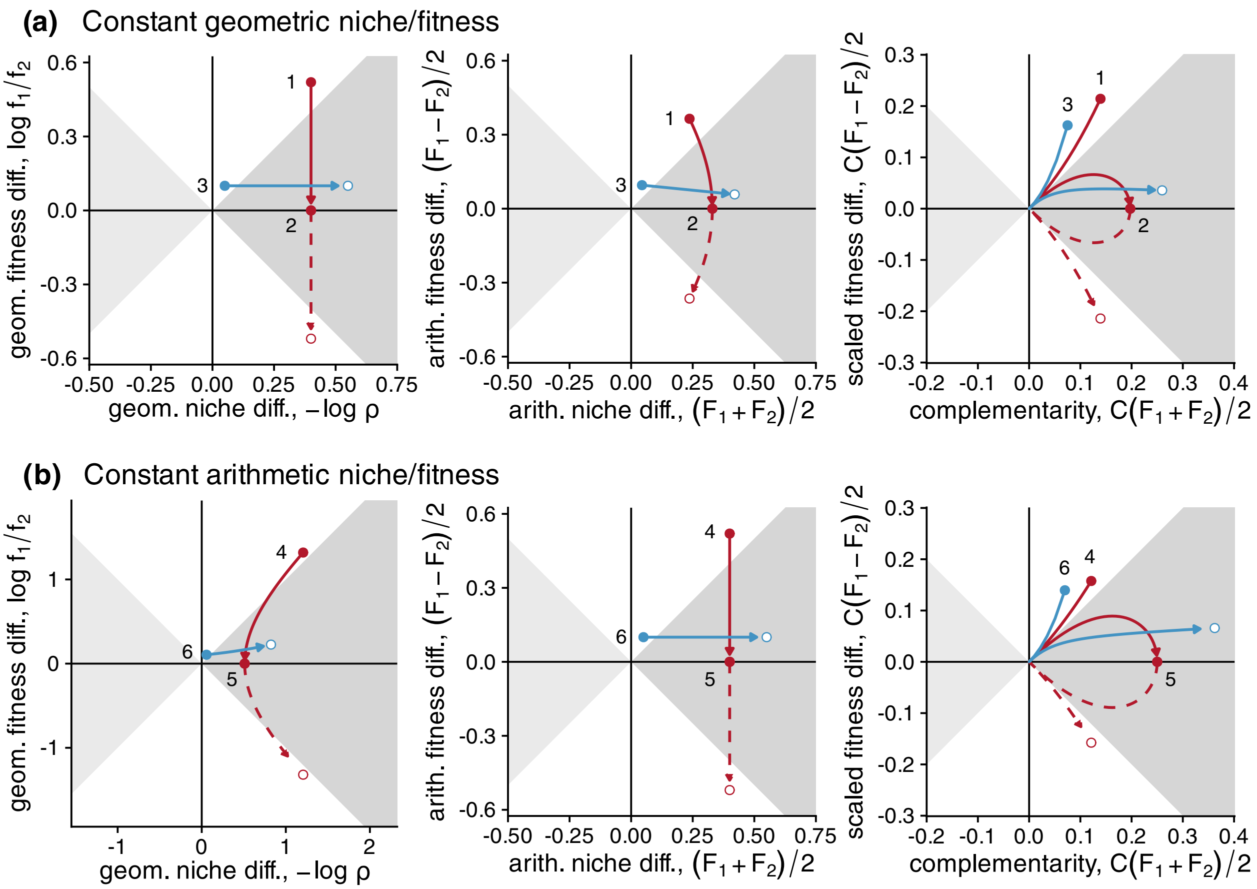


**Figure S5.1: Comparing different definitions of niche and fitness differences.** Focusing on the Lotka–Volterra model of competition, we compare three definitions of niche and fitness differences. In each panel, we visualize the geometric (left plot), arithmetic (middle plot), and complementarity-based (right plot) niche and fitness metrics calculated for the same simulated processes. Arrows from solid to open points indicate the direction of each process; solid lines indicate positive fitness difference and dashed lines indicate negative fitness difference. In each case, the niche difference metric is on the horizontal axis and the fitness difference on the vertical axis. We follow the conventions of Figure 1a in the main text; for instance, coexistence (dark gray) requires $ND > |FD|$. **(a) Constant geometric niche or fitness difference.** We vary one of the geometric metrics (left plot) while keeping the other constant: decreasing fitness difference (red; 1: decreasing $FD_{geo}$ from $0.52 \to 0$ (solid line) and 2: $0 \to -0.52$ (dashed line), with constant $ND_{geo} = 0.4$) or increasing niche difference (blue; 3: $FD_{geo}$ from $0.05 \to 0.55$, with constant $FD_{geo} = 0.1$). **(b) Constant arithmetic niche or fitness difference.** We now vary the arithmetic metrics (center plot): decreasing fitness difference (red; 4–5) or increasing niche difference (blue; 6); parameter values are the same as in (a) except using $ND_{ari}, FD_{ari}$ instead of $ND_{geo}, FD_{geo}$.

### S6 Resource competition model with interference

We generalize a one-resource competition model from Tilman (1982). We consider $n$ species, each with biomass $N_i$, and a single limiting resource, with abundance $R$. The dynamics of the general model are given by the following equations:

$$\frac{dN_i}{dt} = N_i \left[\varepsilon_i u_i \left(R, N_1, \ldots, N_n\right) - m_i \left(N_1, \ldots, N_n\right)\right] \tag{S6.1}$$

$$\frac{dR}{dt} = g\left(R\right) - \sum_{i=1}^{n} N_i u_i \left(R, N_1, \ldots, N_n\right) + \sum_{i=1}^{n} \varphi_i N_i m_i \left(N_1, \ldots, N_n\right) \tag{S6.2}$$

This general treatment of $m_i$ (the mortality function of species $i$) follows Gross (2008); meanwhile, $u_i$ (uptake rate) is a function of not only resource abundance $R$ but also species abundances, allowing for interference effects: $u_i\left(R, N_1, N_2, \ldots, N_n\right)$ (as in Amarasekare 2002). Finally, as in the classic model, resource dynamics are governed by resource supply $g$ and by uptake (where species $i$ has resource use efficiency $\varepsilon_i$). However, we add a term representing return of the resource from dead biomass (where $\varphi_i$ is the amount of resource that returns from one biomass unit of species $i$).

**Specific model** Now we consider a specific case of the general model, representing interference competition between species competing for a fixed pool of resources, closely related to the lottery model for plant competition.

$$m_i\left(N_1, \ldots, N_n\right) = \mu_i \tag{S6.3}$$

$$u_i\left(R, N_1, \ldots, N_i\right) = \frac{v_i R}{1 + \sum_{j=1}^{n} \beta_{ij} N_j} \tag{S6.4}$$

$$\varphi_i = \varepsilon_i^{-1} \tag{S6.5}$$

$$g\left(R\right) = 0 \tag{S6.6}$$

Since $\varphi_i = \varepsilon_i^{-1}$ (i.e. dead individuals return all resources to the pool) and $g\left(R\right) = 0$ (i.e. there is no resource input), the system is closed and the total amount of resource in the resource pool $R$ and in species biomass (i.e., $\varepsilon_i^{-1} N_i$) is fixed: $dR/dt + \sum_{i=1}^{n} \varepsilon_i^{-1} \cdot dN_i/dt = 0$. We let $R_0$ denote this total resource quantity:

$$R_0 \equiv R + \sum_{i=1}^{n} \varepsilon_i^{-1} N_i \tag{S6.7}$$

Then we can rewrite the system without the state variable for the resource pool using the relationship $R = R_0 - \sum_{i=1}^{n} \varepsilon_i^{-1} N_i$. Then we have:

$$\frac{dN_i}{dt} = N_i \left[ \frac{\varepsilon_i v_i \left( R_0 - \sum_{j=1}^{n} \varepsilon_j^{-1} N_j \right)}{1 + \sum_{j=1}^{n} \beta_{ij} N_j} - \mu_i \right] \tag{S6.8}$$

We show the dynamics of this model in Supplemental Figure S6.1 for $n = 1$ (panel a), 2 (panel b), or 20 species (panel c).

**Relationship to Tilman's $R^*$** This model is an extension of the one-resource model of Tilman (1982). Just as in that model, we can define a quantity $R_i^*$, the lowest resource concentration at which species $i$ can maintain nonnegative population growth. Assuming all species are at low abundance and setting the per-capita growth rate (the bracketed term in equation S6.8) to zero and allowing resource level to vary arbitrarily, the condition for $R_i^*$ is $\varepsilon_i v_i R_i^* - \mu_i = 0$, thus giving that

$$R_i^* \equiv \frac{\mu_i}{\varepsilon_i v_i}, \tag{S6.9}$$

corresponding exactly to the expression given by Tilman (1982). However, we note that this expression must be interpreted slightly differently: since species interfere with themselves and others, the actual resource level at equilibrium $\hat{R}$ (where the hat $\hat{\cdot}$ represents equilibrium) will be greater than $R_i^*$ if the consumer is at some nonzero equilibrium abundance (e.g., Supplemental Figure S6.1a). In other words, $R_i^*$ summarizes resource competition ability due to mortality, resource uptake, and use efficiency, but does not account for the effect of interference ($\beta_{ii}, \beta_{ij}$).

**Calculating competition coefficients** We note that the specific model belongs to the class considered in Appendix S1. To demonstrate this, we can show that the per-capita growth rate (the bracketed term in equation S6.8) is a case of equation S1.2. Multiplying the per-capita growth rate by the positive quantity $\varepsilon_i^{-1} v_i^{-1} \left( 1 + \sum_{j=1}^{n} \beta_{ij} N_j \right)$ and noting that $\mu_i \varepsilon_i^{-1} v_i^{-1} = R_i^*$ gives

$$\varepsilon_i^{-1} v_i^{-1} \left( 1 + \sum_{j=1}^{n} \beta_{ij} N_j \right) \cdot \frac{1}{N_i} \cdot \frac{dN_i}{dt} = R_0 - \sum_{j=1}^{n} \varepsilon_j^{-1} N_j - R_i^* \left( 1 + \sum_{j=1}^{n} \beta_{ij} N_j \right) \tag{S6.10}$$

$$= (R_0 - R_i^*) - \sum_{j=1}^{n} \left( R_i^* \beta_{ij} + \varepsilon_j^{-1} \right) N_j, \tag{S6.11}$$

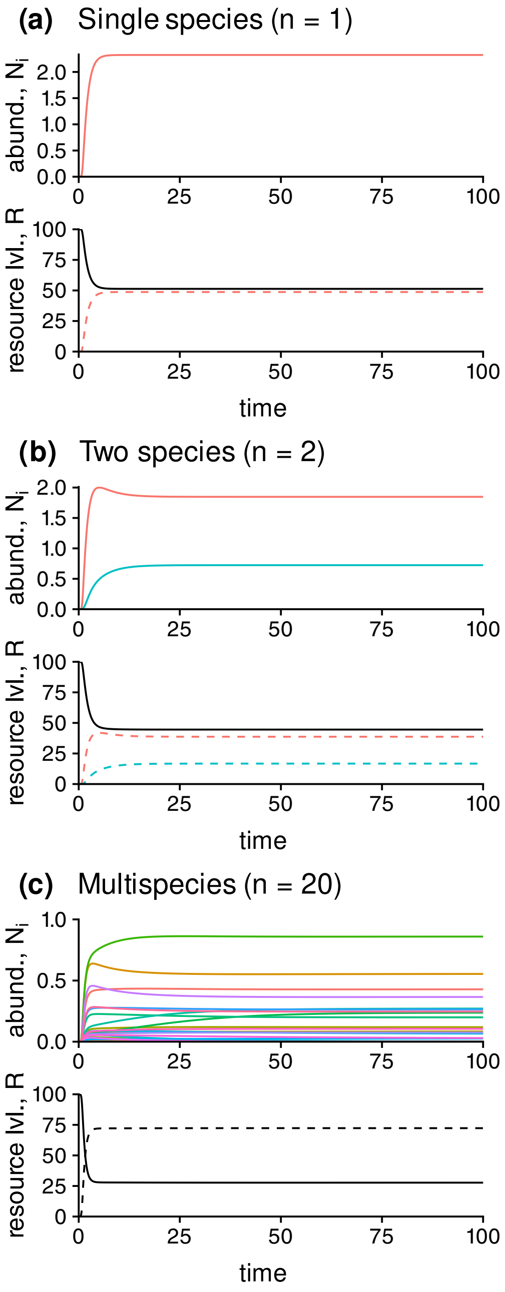


**Figure S6.1: Time series for the resource competition model.** We illustrate dynamics of species abundances $N_i$ (top plot in each panel: solid lines) and resource level $R$ (bottom plot: black lines) for **(a) a single species** (i.e., a monoculture), **(b) two competing species**, or **(c) multispecies competition** ($n = 20$). To show that total resource level is conserved, we also illustrate the resources contained in each species' biomass $N_i/\varepsilon_i$ (panels a–b, bottom plot: colored dashed lines) or the total in the multispecies case (panel c, bottom plot: black dashed line). Parameters match the reference multispecies model used in main text Figure 7 and described in Appendix S9, taking one, two, or all species as needed.

which responds linearly to the species abundances $N_j$. Thus, further dividing by $\left(R_0 - R_i^*\right)$ produces an expression with an intercept of 1, matching equation S1.2, and we can define

$$h_i^{-1}\left(\frac{1}{N_i}\cdot\frac{dN_i}{dt}\right) \equiv \frac{1+\sum_{j=1}^{n}\beta_{ij}N_j}{\varepsilon_i v_i\left(R_0 - R_i^*\right)}\cdot\frac{1}{N_i}\cdot\frac{dN_i}{dt} \tag{S6.12}$$

$$= 1 - \sum_{j=1}^{n}\alpha_{ij}N_j, \tag{S6.13}$$

where $h_i^{-1}$ is the inverse function from Appendix S1 that converts a per-capita growth rate to the quantity $1 - \sum_j \alpha_{ij}N_j$, and

$$\alpha_{ij} \equiv \frac{R_i^*\beta_{ij} + \varepsilon_j^{-1}}{R_0 - R_i^*} \tag{S6.14}$$

is the effective competition coefficient, accounting for species $j$'s total impact on species $i$ due to interference and removal of resources from the pool. We note that the numerator of this expression is independent of the actual total resource level $R_0$, thus determining the contribution of species traits and interaction parameters to interference ($R_i^*\beta_{ij}$) and resource monopolization ($\varepsilon_j^{-1}$), while the denominator scales this effect according to the portion of the resource pool available to species $i$ ($R_0 - R_i^*$). Thus, we can define a resource availability–independent competition coefficient

$$b_{ij} \equiv R_i^*\beta_{ij} + \varepsilon_j^{-1}, \tag{S6.15}$$

such that $\alpha_{ij} = b_{ij}/\left(R_0 - R_i^*\right)$.

**Calculating coexistence components** Applying the definitions in equations S1.10 and S1.11, we find that niche overlap is given by

$$\rho = \sqrt{\frac{\left(R_1^*\beta_{12}+\varepsilon_2^{-1}\right)\left(R_2^*\beta_{21}+\varepsilon_1^{-1}\right)}{\left(R_1^*\beta_{11}+\varepsilon_1^{-1}\right)\left(R_2^*\beta_{22}+\varepsilon_2^{-1}\right)}} = \sqrt{\frac{b_{12}b_{21}}{b_{11}b_{22}}}, \tag{S6.16}$$

which is completely independent of resource level $R_0$, while fitness ratio is

$$\frac{f_1}{f_2} = \frac{R_0 - R_1^*}{R_0 - R_2^*}\sqrt{\frac{\left(R_2^*\beta_{22}+\varepsilon_2^{-1}\right)\left(R_2^*\beta_{21}+\varepsilon_1^{-1}\right)}{\left(R_1^*\beta_{11}+\varepsilon_1^{-1}\right)\left(R_1^*\beta_{12}+\varepsilon_2^{-1}\right)}} = \frac{R_0 - R_1^*}{R_0 - R_2^*}\sqrt{\frac{b_{22}b_{21}}{b_{11}b_{12}}}, \tag{S6.17}$$

where the term outside the square root represents how resource level affects the resource competition hierarchy: $(R_0 - R_1^*) \,/\, (R_0 - R_2^*)$ favors the species with lower $R_i^*$, but approaches 1 when $R_0$ is significantly large. Note that this expression is analogous to the case for other models where some traits appear in the fitness ratio but not the niche overlap: for instance, Letten et al. (2017) found similar expressions for Tilman (1982)'s model of two substitutable resources, and intrinsic fecundity affects fitness ratio but not niche overlap in the Beverton–Holt model for annual plants (Adler et al. 2010).

**Approximation of more complex resource dynamics** Note the system defined by equation S6.8 can also approximate a model with more complex resource dynamics. As an illustration, we consider a model where supply $g(R)$ and return $\varphi_i$ are such that $dR/dt$ decreases monotonically with $R$; note however that the argument could be generalized further. We treat the case where resource dynamics are fast relative to consumer dynamics (i.e., parameters are such that resources are under tight biotic control). Accordingly, at any given time point, $R$ is very near a value determined only by consumer populations at that time, known as the *quasi-equilibrium*. We can solve for this by treating consumer populations as fixed and solving $dR/dt = 0$ to find

$$R \approx f^{-1}\left(0; N_1, \ldots, N_n\right) \equiv \hat{R}_q, \tag{S6.18}$$

where $f(R; N_1, \ldots, N_n)$ is the rate of change in resource as a function of resource level, for some given fixed abundance of $N_1, \ldots, N_n$, and $f^{-1}$ is its inverse; note that the semicolon separates the argument of the function (resource level $R$) from parameters. Assuming the system is close to its equilibrium $\hat{R}, \hat{N}_1, \ldots, \hat{N}_n$ (i.e., for the resource with all its consumers), we can further approximate resource level using a Taylor expansion of $\hat{R}_q$ around the equilibrium as

$$R \approx \hat{R} + \sum_{i=1}^{n} k_i \left(N_i - \hat{N}_i\right) = \left(\hat{R} - \sum_{i=1}^{n} k_i \hat{N}_i\right) + \sum_{i=1}^{n} k_i N_i, \tag{S6.19}$$

where $k_i = \left(\partial \hat{R}_q / \partial N_i\right)\big|_{\hat{N}_1,\ldots,\hat{N}_n}$ is the partial derivative of quasi-equilibrium resource value with respect to species $i$, evaluated near the equilibrium. Then we can write $R \approx \tilde{R}_0 - \sum_{i=1}^{n} \tilde{\varepsilon}_i^{-1} N_i$, where

$$\tilde{\varepsilon}_i = -\left(\left.\frac{\partial \hat{R}_q}{\partial N_i}\right|_{\hat{N}_1,\ldots,\hat{N}_n}\right)^{-1} \quad \text{and} \quad \tilde{R}_0 = \hat{R} + \sum_{i=1}^{n} \tilde{\varepsilon}_i^{-1} \hat{N}_i. \tag{S6.20}$$

Accordingly, we can see that our approximation to the full model has the same form as the simple model in equation S6.8:

$$\frac{dN_i}{dt} = N_i \left[ \frac{\tilde{\varepsilon}_i \tilde{v}_i \left( \tilde{R}_0 - \sum_{i=1}^{n} \tilde{\varepsilon}_i^{-1} N_i \right)}{1 + \sum_{j=1}^{n} \beta_{ij} N_j} - \mu_i \right], \tag{S6.21}$$

where furthermore

$$\tilde{v}_i = \frac{v_i \varepsilon_i}{\tilde{\varepsilon}_i} \tag{S6.22}$$

and the tildes $\tilde{\varepsilon}_i, \tilde{R}, \tilde{v}_i$ indicate that these are the effective parameters in the simplified model that approximate the dynamics of the more complex model.

### S7 Fitting the resource competition model to empirical data

We used data from Wedin and Tilman (1993) for the species pair *Agropyron repens* and *Poa pratensis* to fit the competition model. We digitized biomass data from the original publication figures using WebPlotDigitizer v4.6 (Rohatgi 2022) to obtain biomass data for plots of these species growing individually (Wedin and Tilman 1993, Figure 1) and in competition (Wedin and Tilman 1993, Figure 6a) along a soil nitrogen gradient, in the fifth year of the experiment. Since the authors observed that community composition was relatively stable at this point in the experiment, we assumed this represented the equilibrium of our dynamic model, and fit the parameters $R_i^*$,$b_{ij}$ (where $i, j = Poa$ or $Agr.$ and $i = j$ for intraspecific competition) to these equilibrium biomasses. To better fit the data, we allowed competition parameters to vary with soil nitrogen, a departure from our original model, according to

$$b_{ij} = f_{ij} + g_{ij} \cdot \text{nitrogen}, \tag{S7.1}$$

where $f_{ij}, g_{ij}$ are the intercept and slope of the nitrogen relationship, respectively, and subscripts indicate that this is species $i$'s competitive sensitivity to species $j$. We obtained intrinsic parameters $R_i^*, b_{ii}$ from monoculture biomass and $b_{ij}$ from competition biomass using nonlinear least-squares regression (NLS); since NLS requires suitable starting values from which to optimize parameters, we first obtained rough estimates for each regression.

**Monoculture fits** We first used monoculture data to fit the single-species parameters $R^*_{Poa}, R^*_{Agr.}$ and $b_{Poa,Poa}, b_{Agr.,Agr.}$ by fitting the relationship,

$$\text{biomass}_i \sim \frac{\text{nitrogen} - R_i^*}{f_{ii} + g_{ii} \cdot \text{nitrogen}}, \tag{S7.2}$$

which correspond to equation 17 in the main text. To obtain starting estimates of the parameters, we used linear regressions: since $R^*$ in our model is the value of nitrogen for which biomass is zero, we obtained a rough estimate using the $x$-intercept of the linear regression fit $\text{biomass}_i \sim b_0 + b_1 \cdot \text{nitrogen}$, which is $R_i^* \approx -b_0/b_1$. Next, we estimated the coefficients $f_{ii}, g_{ii}$ by rearranging equation S7.2 to obtain

$$\frac{\text{nitrogen} - R_i^*}{\text{biomass}_i} \sim f_{ii} + g_{ii} \cdot \text{nitrogen} \tag{S7.3}$$

and performing linear regression using the rough value for $R_i^*$. Final fits for $R_i^*, f_{ii}, g_{ii}$ were obtained simultaneously by starting from the rough values and directly fitting equation S7.2 using NLS; these are shown in Supplemental Table S7.1, visualized alongside the original data in Supplemental Figure S7.1, and the resulting values of $b_{ii}$ are plotted across the nitrogen gradient in Supplemental Figure S7.2. Values of $g_{ii}$ inferred by NLS were 0 for both species, suggesting that the original model provides a good description of intrinsic yield in the system.

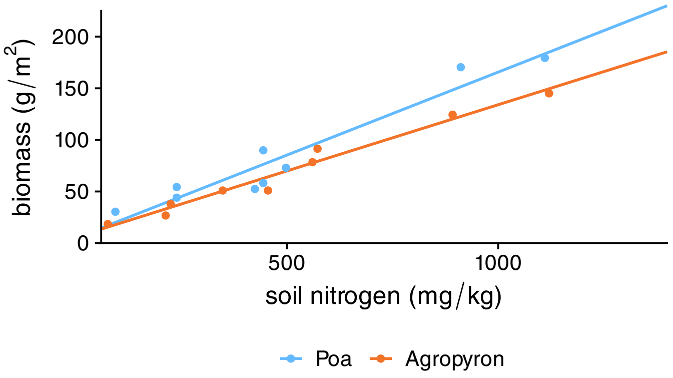


**Figure S7.1: Monoculture fits.** Points show the biomass (vertical axis) of monoculture plots of *Poa* (blue) and *Agropyron* (orange) at different levels of soil nitrogen (horizontal axis) in the experiment of Wedin and Tilman (1993); we visualize our NLS model fits with lines in the corresponding color.

**Competition fits** We then fit the competitive coefficients $b_{Agr,Poa}$ and $b_{Poa,Agr.}$ using the monoculture-derived parameters and the data from competition plots, assuming that abundances at the end of the experiment approximated the coexistence equilibrium of our model. First, we calculated the biomass at equilibrium of species $i$ (= *Agr*. or *Poa*) as

$$\text{biomass}_i \sim \frac{\left(\text{nitrogen} - R_i^*\right) b_{jj} - \left(\text{nitrogen} - R_j^*\right) b_{ij}}{b_{ii}b_{jj} - b_{ij}b_{ji}}, \tag{S7.4}$$

which is simply calculated from equation 4 in the main text using the model-specific definitions in equations S7.1 (in this appendix) and 15–17 (in Box 3 of the main text). We fit the model for biomass of both species simultaneously using NLS. To improve convergence, we fit the data in two passes. Since it was difficult to find initial estimates for $f_{ij}, g_{ij}$

allowing for convergence, we first constrained $f_{ij}$ according to:

$$f_{ij} = \frac{\left(w_i - R_i^*\right)\left(f_j + g_j w_i\right)}{\left(w_i - R_j^*\right)} - g_{ij} w_i, \tag{S7.5}$$

where $w_i$ is a fixed parameter representing the nitrogen level at which the invasion growth of $i$ is zero (i.e. $i$'s persistence boundary), such that NLS only fit one parameter per interaction (i.e., $g_{ij}$). For this first pass, we used $g_{Agr,Poa} = g_{Poa,Agr.} = 0.001$ as initial values and enforced fixed values $w_{Agr.} = 1250$, $w_{Poa} = 0$. Next, we removed this constraint by performing a second round of NLS, starting from the rough fit and also allowing the persistence boundaries $w_{Agr.}$ and $w_{Poa}$ to vary. Final fits, after transforming from $w_i$ back to $f_{ij}$, are shown together with monoculture fits in Supplemental Table S7.1 and visualized in Figure S7.2 in the main text; resulting values of $b_{ij}$ are plotted across the nitrogen gradient in Supplemental Figure S7.2.

**Additive partition components** Using the fitted model, we calculated the additive partition components across the resource gradient (Appendix Figure S7.3). First, we used the definition in Appendix S4, which partitions the overyielding effect (solid line, dark green) relative to average intrinsic yield (top panel). Next, we adjusted the choice of expected relative yields to partition the relative degree of transgressive overyielding (solid line, light green) by choosing $\mathrm{RY}_{Poa} = 1$ and $\mathrm{RY}_{Agr.} = 0$ (bottom panel). As expected, complementarity (dotted lines) was independent of this choice and positive for all nitrogen values allowing coexistence. Meanwhile, reflecting the low fitness of the more productive species *Poa*, selection (dashed line) was negative across most (relative definition) or all (transgressive definition) of the nitrogen gradient. As a result, both forms of overyielding were impossible at most nitrogen levels used in the experiment, though overyielding relative to average yield was possible at high nitrogen.

**Table S7.1: Fitted parameters for the Wedin and Tilman (1993) dataset.**

| parameter | | species | value | units |
|---|---|---|---|---|
| monoculture | $R_i^*$ | *Poa* | $-30.877$ | $\frac{\text{mg N}}{\text{kg}}$ |
| | | *Agr.* | $-45.293$ | |
| | $f_{ii}$ | *Poa* | 6.223 | $\frac{\text{m}^2\cdot\text{mg N}}{\text{kg}\cdot\text{g bio.}}$ |
| | | *Agr.* | 7.798 | |
| | $g_{ii}$ | *Poa* | 0.000 | $\frac{\text{m}^2}{\text{g bio.}}$ |
| | | *Agr.* | 0.000 | |
| competition | $f_{ij}$ | *Poa*, *Agr.* | 7.313 | $\frac{\text{m}^2\cdot\text{mg N}}{\text{kg}\cdot\text{g bio.}}$ |
| | | *Agr.*, *Poa* | 3.932 | |
| | $g_{ij}$ | *Poa*, *Agr.* | $-0.0001539$ | $\frac{\text{m}^2}{\text{g bio.}}$ |
| | | *Agr.*, *Poa* | 0.001859 | |

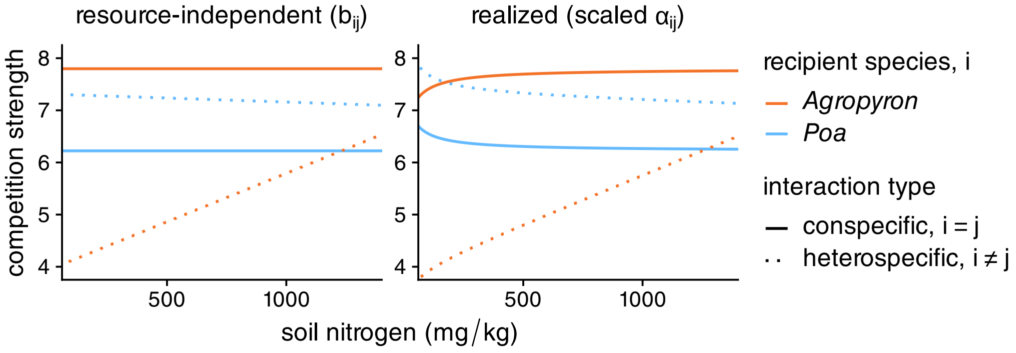


**Figure S7.2: Inferred competition coefficients vary with soil nitrogen.** We visualize the competitive coefficients as a function of soil nitrogen according to the fits in S7.1. We directly plot the intraspecific (solid lines) and interspecific (dotted lines) resource level–independent coefficients, $b_{ij}$ (left panel). To visualize how competition was affected by resource level, we scaled the actual competition coefficients to facilitate comparison across resource levels: we multiplied all $\alpha_{ij}$s by the geometric mean of the quantity $(\text{nitrogen} - R_i^*)$ for the two species.

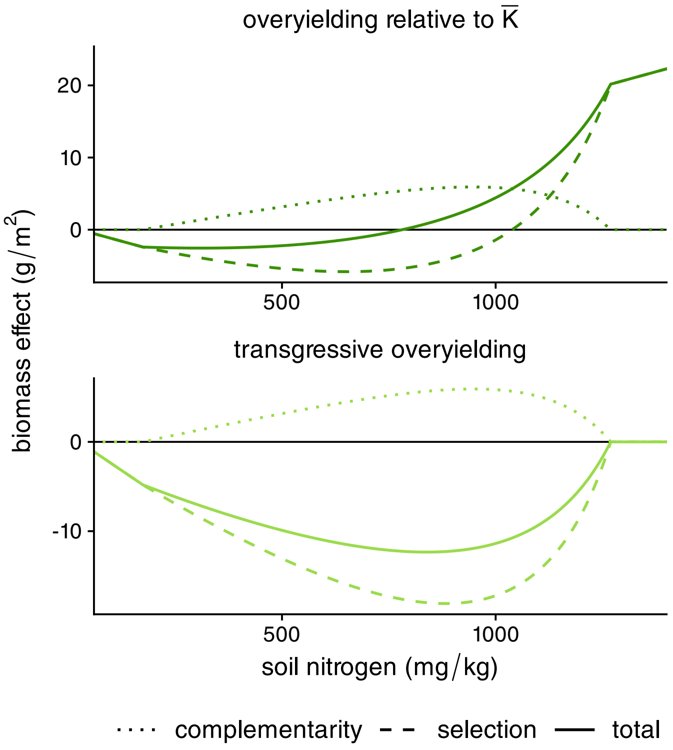


**Figure S7.3: Additive partition components calculated for the fitted biomasses.** For varying nitrogen levels (horizontal axis), we used the fitted competition parameters to predict total biomass (solid line) and calculate the additive partition components (selection: dashed line, complementarity: dotted line), all in units of biomass (vertical axis). We chose expected relative yield either according to the approach elsewhere in the text, which partitions overyielding relative to $\overline{K}$ (top panel, dark green), or according to the expectation that the higher yielding species dominates, which is equivalent to partitioning the degree of transgressive overyielding (bottom panel, light green).

#### S8 Applying the framework to other ecosystem functions

While we have chosen to show results for biomass for simplicity, we show in this Appendix how our framework straightforwardly generalizes to other functions. For some arbitrary function $x$, let $\varphi_i^{(x)}$ represent the level of this function *per unit biomass* of species $i$ at equilibrium, we can extend equation 4 from the main text to write the *overall* functional contribution of species $i$ as $\varphi_i^{(x)}\hat{N}_i = F_i \cdot \varphi_i^{(x)} K_i / \left(1-\rho^2\right)$. As all subsequent results are derived from this equation, we can simply replace $K_i$ with $\Phi_i^{(x)} \equiv \varphi_i^{(x)} K_i$ in equations 6–8 from the main text to obtain their analogues for function $x$. With the further assumption that function per unit biomass is constant for each species, $\Phi_i^{(x)}$ can simply be interpreted as species $i$'s intrinsic yield for function $x$.

**Two functions** We consider the case where each species has a higher yield for a different function; for simplicity, we call these $K$ (biomass productivity) and $\Phi$. The boundaries are

$$\frac{f_1}{f_2} > \rho \cdot \frac{K_1}{K_2} \quad \text{and} \quad \frac{f_2}{f_1} > \rho \cdot \frac{\Phi_2}{\Phi_1} \tag{S8.1}$$

Thus

$$\rho \cdot \frac{K_1}{K_2} < \frac{f_1}{f_2} < \rho^{-1} \frac{\Phi_1}{\Phi_2} \tag{S8.2}$$

When $\rho$ is large, it is not possible to satisfy this equation; the maximum $\rho$ for which the inequality can hold can be found by solving $\rho \cdot K_1/K_2 = f_1/f_2 = \rho^{-1} \cdot \Phi_1/\Phi_2$ to obtain

$$\rho = \sqrt{\frac{\Phi_1}{\Phi_2} \cdot \frac{K_2}{K_1}}, \tag{S8.3}$$

which can also be rewritten by taking logarithms to obtain $-\log \rho > \frac{1}{2} \log\left(K_1/K_2\right) - \frac{1}{2} \log\left(\Phi_1/\Phi_2\right)$. That is, niche difference must be sufficient to overcome the strength of the tradeoff between productivity and the second function. The fitness at which this occurs, the target fitness ratio for multifunctionality, is $f_1/f_2 = \rho \cdot K_1/K_2$, which gives

$$\frac{f_1}{f_2} = \sqrt{\frac{\Phi_1}{\Phi_2} \cdot \frac{K_1}{K_2}} \tag{S8.4}$$

i.e., the geometric mean of the biomass and function yield ratios.

**General conditions for multifunctionality** In the most general case, if we have any number of functions $\Phi_i^{(1)}, \Phi_i^{(2)}, \ldots$, relevant bounds are the most restrictive ones, e.g.,

$$\rho \cdot \max\left(1, \frac{\Phi_1^{(1)}}{\Phi_2^{(1)}}, \frac{\Phi_1^{(2)}}{\Phi_2^{(2)}}, \cdots\right) < \frac{f_1}{f_2} \tag{S8.5}$$

and thus the condition for all functions to show transgressive overyielding is

$$\rho \cdot \max\left(1, \frac{\Phi_1^{(1)}}{\Phi_2^{(1)}}, \frac{\Phi_1^{(2)}}{\Phi_2^{(2)}}, \cdots\right) < \frac{f_1}{f_2} < \rho^{-1} \cdot \min\left(1, \frac{\Phi_1^{(1)}}{\Phi_2^{(1)}}, \frac{\Phi_1^{(2)}}{\Phi_2^{(2)}}, \cdots\right). \tag{S8.6}$$

The $\rho$ at which simultaneous transgressive overyielding becomes possible is

$$\rho = \sqrt{\frac{\min\left(1, \frac{\Phi_1^{(1)}}{\Phi_2^{(1)}}, \frac{\Phi_1^{(2)}}{\Phi_2^{(2)}}, \cdots\right)}{\max\left(1, \frac{\Phi_1^{(1)}}{\Phi_2^{(1)}}, \frac{\Phi_1^{(2)}}{\Phi_2^{(2)}}, \cdots\right)}} \tag{S8.7}$$

and the target fitness for equalization is

$$\frac{f_1}{f_2} = \sqrt{\min\left(1, \frac{\Phi_1^{(1)}}{\Phi_2^{(1)}}, \frac{\Phi_1^{(2)}}{\Phi_2^{(2)}}, \cdots\right) \cdot \max\left(1, \frac{\Phi_1^{(1)}}{\Phi_2^{(1)}}, \frac{\Phi_1^{(2)}}{\Phi_2^{(2)}}, \cdots\right)}. \tag{S8.8}$$

### S9 Numerical simulations

#### S9.1 General model simulations

To simulate the effect of varying niche, fitness, and functional imbalance, we applied equation S1.19 to calculate the total biomass at equilibrium for representative values of niche difference $1-\rho$, fitness ratio $f_1/f_2$, and intrinsic yield $K_i$ (Table S9.1), corresponding to an underlying Lotka–Volterra model.

**Table S9.1: Parameters for general simulations.**

| parameter | species | value(s) |
|---|---|---|
| intrinsic yield, $K_i$ | 1 | 1.0 |
| | 2 | 0.694 |
| niche difference, $1-\rho$ | — | 0.29 (0.15, 0.22, 0.36) |
| fitness ratio, $f_1/f_2$ | — | 1.13 (0.93, 1.03, 1.23) |
| intrinsic (function) yield, $\Phi_i$ | 1 | 0.7225 |
| | 2 | 1.0 |

**Niche–fitness space diagrams** To produce the niche–fitness space diagram in Figure 1b (main text), we used the fixed values of $K_i$ and visualized the conditions for alternative stable states ($\rho^{-1} < f_1/f_2 < \rho$; light gray), coexistence ($\rho < f_1/f_2 < \rho^{-1}$; dark gray or green), and transgressive overyielding (main text equation 7; green) for different values of niche and fitness. To match the linear boundaries of the conceptual diagram (Figure 1a), the figure plots ND $= -\log\rho$ on the horizontal and FD $= \log f_1/f_2$ on the vertical axis, but indicates the more familiar $1-\rho$ and $f_1/f_2$ on the axis scales. Main text Figure 3d and Supplemental Figure S2.1 show the same analysis, but with an additional boundary for overyielding relative to average yield (equation S2.8).

**Effect of varying individual components** We simulated the effect of varying each component individually in Figure 2 (main text) by using the baseline values in Table S9.1 and continuously varying a focal component: niche difference $1-\rho$ (panel a), fitness ratio $f_1/f_2$ (panel b), or intrinsic yield of the lower yielding species $K_2$ (panel c). For the niche and fitness simulations, the other metric (respectively fitness and niche) was varied between four values to produce the different lines (main and parenthesized values in Table S9.1); in all cases, remaining parameters were fixed (main values in table).

**Additive partition** We repeated the above simulations for our additive partition analyses (main text Figure S4a–c); note, however, that the analysis of functional equalization (panel c) adds multiple lines for different fitness ratios, unlike the analysis above. To calculate complementarity and selection, we calculated species biomasses from main text equation 4 and applied the definitions in main text equation 10.

**Multifunctionality** To analyze multifunctionality, we additionally considered each species' intrinsic yield in terms of (non-biomass) function, $\Phi_i$ (also given in Table S9.1). For the same values of the other components, and following the same methods as above, we plotted outcomes on the niche–fitness space (Figure 6a) and the effect of varying fitness ratio (Figure 6b).

#### S9.2 Consumer–resource model simulations

For the remaining simulations, we used the consumer–resource model described in main text Box 3 and Appendix S6.

**Table S9.2: Parameters for two-species consumer–resource model simulations.**

| parameter | | species | value | description |
|---|---|---|---|---|
| aggregate values | $R_i^*$ | 1 | 134.0 | Tilman's $R^*$ |
| | | 2 | 2.0 | |
| | $a_{ii}$ | 1 | 4.320 | intraspecific competition |
| | | 2 | 9.0 | |
| | $a_{ij}$ | 1, 2 | 4.80 | interspecific competition |
| | | 2, 1 | 5.184 | |
| underlying traits | $\beta_{ii}$ | 1 | 0.0239 | intraspecific interference |
| | | 2 | 3.50 | |
| | $\beta_{ij}$ | 1, 2 | 0.0209 | interspecific interference |
| | | 2, 1 | 2.036 | |
| | $\varepsilon_i$ | 1 | 0.90 | resource use efficiency |
| | | 2 | 0.50 | |
| | $\mu_i$ | 1 | 5.0 | mortality rate |
| | | 2 | 1.0 | |
| | $v_i$ | 1 | 0.0415 | uptake ability |
| | | 2 | 1.0 | |

**Two-species consumer–resource simulation** To illustrate our theoretical results (Appendix S6), we plotted the effect of varying total resource level $R_0$ in main text Figure 4b.

We used the model-specific definitions of $\rho$ and $f_1/f_2$ in Box 3 and the general expression in equation 4 (main text) to calculate species biomasses for continuously varying $R_0$; values of the aggregated parameters $R_i^*$ and $a_{ij}$ and corresponding underlying parameters are given in Table S9.2.

**Multispecies simulations: reference model** For our multispecies analyses, we generated an $n = 20$ species community (the *reference model*) with traits drawn from random distributions: $\beta_{ii}, \beta_{ij}, \mu_i, v_i$ should be greater than zero for a model of competition with interference, so we drew them from log-normal distributions, while $\varepsilon_i$ is a resource use efficiency and thus should be between 0 and 1, so we drew it from a beta distribution. All parameter values and distributions are given in Table S9.3. The reference model is indicated with a star ($\star$) in main text Figure 7a–c, and we visualize the distribution of its pairwise functional coexistence metrics (niche, fitness, and intrinsic yield, calculated according to main text Box 3) in Supplemental Figure S9.1a–c.

**Table S9.3: Parameters for multispecies consumer–resource reference model.**

| parameter | | (mean) value | distribution |
|---|---|---|---|
| general parameters | $n$ | 20 | — |
| | $R_0$ | 100.0 | — |
| species traits/interactions | $\beta_{ii}$ | 1.0 | log-normal; cv = 0.1 |
| | $\beta_{ij}$ | 5.0 | log-normal; cv = 0.1 |
| | $\varepsilon_i$ | 0.05 | beta; cv = 0.1 |
| | $\mu_i$ | 0.5 | log-normal; cv = 0.1 |
| | $v_i$ | 2.0 | log-normal; cv = 0.1 |

**Multispecies simulations: varying niche, fitness, and function** There is no simple relationship between the distribution of pairwise niche and fitness metrics and the trait distributions in Supplemental Table S9.3; i.e., changing any trait distribution should affect both niche and fitness distributions. To vary niche, fitness, and intrinsic yield independently, we started from the randomly-drawn reference model (i.e., those in Supplemental Figure S9.1). Using the exact system resulting from this random draw, we directly modified the observed pairwise niche and fitness metrics, and back-calculated sets of underlying parameter values that exactly yielded the desired niche and fitness values. While this could be done in many ways, we chose to keep all parameter values constant except interference ($\beta_{ii}$ and/or $\beta_{ij}$; see below). For niche difference $1 - \rho$ (main text Figure 7a and Supplemental Figure S9.2), we multiplied all pairwise niche overlaps ($\rho_{ij} = \sqrt{\alpha_{ij}\alpha_{ji}/\alpha_{ii}\alpha_{jj}}$) by the same

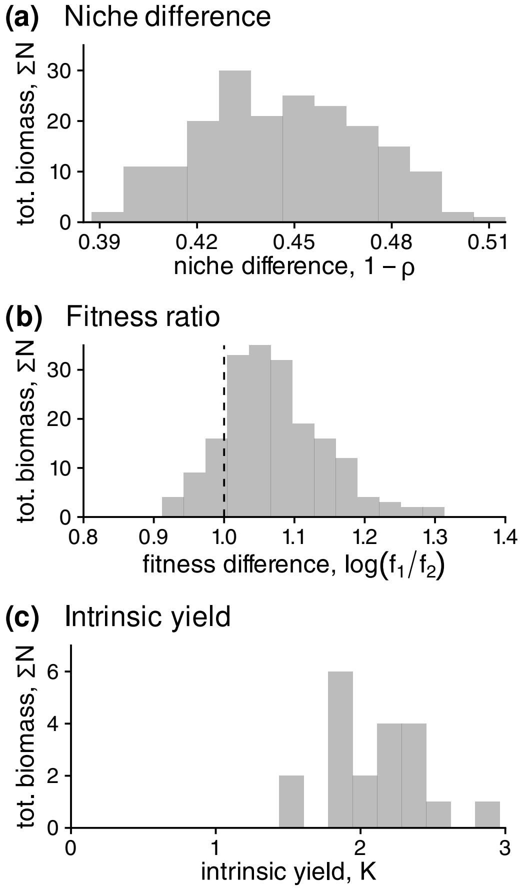


**Figure S9.1: Multispecies simulation: pairwise functional coexistence metrics for the reference model.** Histograms show the distribution of pairwise **(a) niche difference** and **(b) fitness ratio**, as well as **(c) intrinsic yield** for the reference model. For fitness ratio, species subscript 1 indicates the higher yielding species in isolation; note that for most species pairs, this ratio was greater than one (vertical dotted line), indicating the higher yielding species also had higher fitness.

factor (ranging from 0.86 to 1.82) and back-calculated the $\beta_{ij}$ necessary to obtain the exact modified distribution of niche overlaps, with all other parameters as in the reference model. We then compared the median value of $1 - \rho$ to the observed total community biomass. Similarly, for fitness ratio $f_1/f_2$ (main text Figure 7b and Supplemental Figure S9.3), we multiplied all pairwise fitness ratios $f_1/f_2$ by the same factor (0.41 to 1.82; where subscript 1 indicates the higher yielding species) and back-calculated $\beta_{ij}$; we used median $f_1/f_2$ for the analysis. Finally, for intrinsic yield $K_i$ (main text Figure 7b and Supplemental Figure S9.4), we adjusted each value by scaling the difference between its logarithm ($\log K_i$) and the logarithm of the maximum ($\log K_{\max}$) or median ($\log K_{\text{med}}$, shown in Supplemental Figure S9.4) by some factor $c$ (e.g., $K_{i,\text{new}} = \exp\left[\log K_{\max} + c \cdot (\log K_i - \log K_{\max})\right]$, with $c$ ranging between $\pm 1.5$). We then calculated the necessary $\beta_{ii}$ to achieve these yields for the same resource use parameters, and finally the $\beta_{ij}$ necessary to keep the pairwise $\rho$ and $f_1/f_2$ distributions the same as those of the reference model.

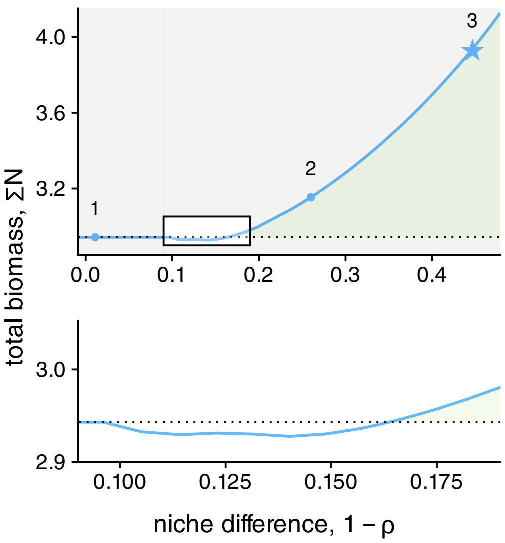


**Figure S9.2: Multispecies simulations: stabilization can decrease total biomass.** Top panel corresponds to Figure 7a in the main text; highlighted region is magnified in the lower panel.

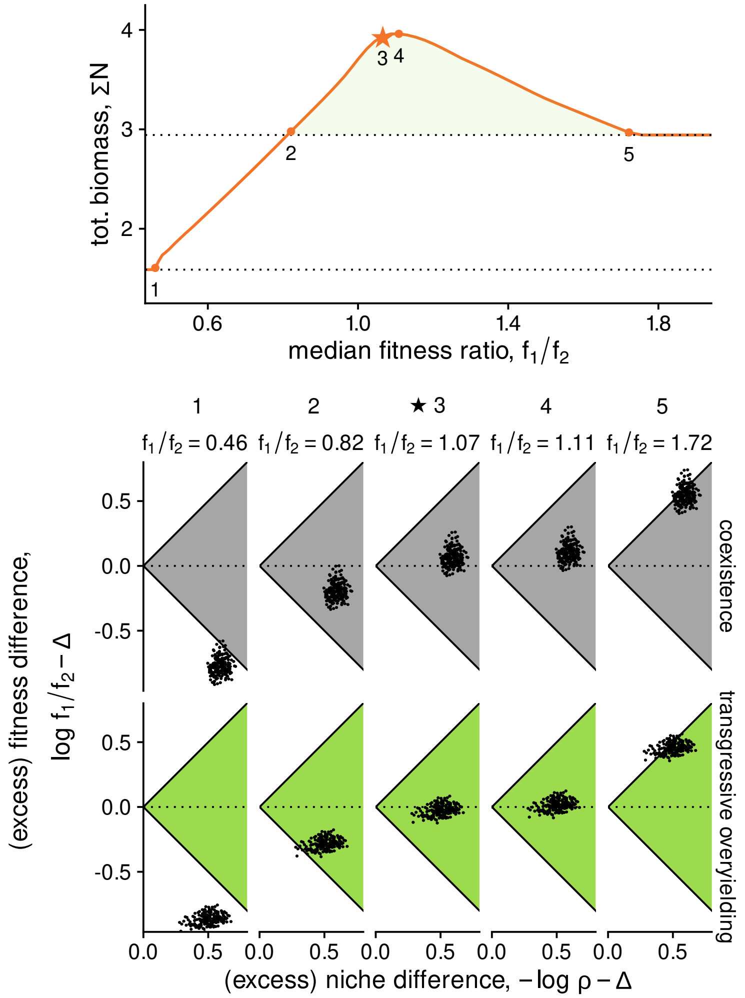


**Figure S9.3: Multispecies simulations: pairwise metrics under functional equalization scenario.** Top panel corresponds to Figure 7b in the main text, with two additional highlighted points (1–5); point 4 is the community with maximum biomass. Bottom panel shows pairwise (excess) niche and fitness differences, $-\log \rho - \Delta$ and $-\log f_1/f_2 - \Delta$, where $\Delta = 0$ for coexistence (ND $>$ |FD|; top row) or $\Delta = \frac{1}{2} \log K_1/K_2$ for overyielding (ND $- \Delta >$ |FD $- \Delta$| as given in Box 1 and equation S1.31; bottom row). Note that the pairwise results generally agree with the outcomes in the multispecies community: coexistence occurs when the distribution of pairwise niche and fitness overlaps the boundaries (points 1, 5; solid boundaries in top row of scatter plots), while overyielding occurs when the excess niche and difference distributions overlap the appropriate boundaries (points 2, 5; solid boundaries in bottom row) and is maximized when the pairwise distribution is nearly centered around FD $- \Delta = 0$ (point 4; horizontal dotted line in bottom row).

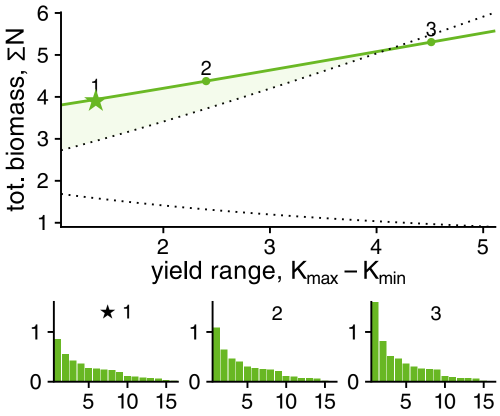


**Figure S9.4: Multispecies simulations: alternate analysis for functional equalization.** The plot follows Figure 7c in the main text. However, we vary the range in intrinsic yield $K_{\text{max}} - K_{\text{min}}$ (horizontal axis and distance between the two dotted lines) while keeping the median fixed, in contrast to the previous analysis which only varied $K_{\text{min}}$. Here, the highest single species yield (upper dotted line) changes as the range is changed. Nonetheless, functional equalization (moving leftwards) increases the degree of overyielding (green shaded region).

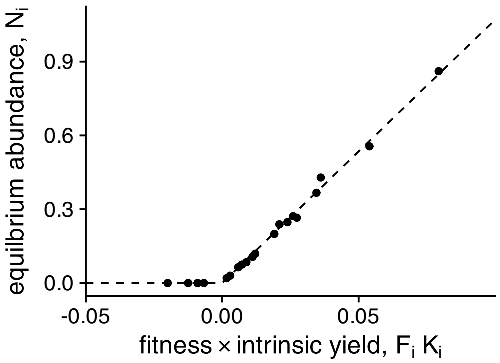


**Figure S9.5: Multispecies simulations: fitness and intrinsic yield predict abundance.** We use our multispecies model to depict a multispecies analogue of our two-species result (equation 4 in Box 1 from the main text) that equilibrium abundance ($\hat{N}_i$, vertical axis) is well determined by $F_i K_i$ i.e., by fitness and intrinsic yield (horizontal axis). For the multispecies model, we calculated $F_i = \text{IGR}_i / r_i$ by simulating the resident equilibrium for each species as invader (i.e., allowing all other species to come to an equilibrium), calculating the invasion growth rate $\text{IGR}_i$ of the focal species, and dividing it by an estimate of $r_i$ obtained by evaluating the growth rate of $i$ in the absence of competition.
